# Wayfarer: A multiscale framework for spatial analysis of tumor progression

**DOI:** 10.64898/2026.02.16.706245

**Authors:** Lambda Moses, Aurelie Herault, Lauriane Cabon, Bianca Dumitrascu

## Abstract

Spatial biology spans multiple length scales, from intracellular organization to tissue-level architecture. Spatial transcriptomics captures this structure, yet most analyses operate at a single spatial resolution, implicitly assuming that biological organization is scale-consistent. In practice, spatial autocorrelation and co-localization are functions of scale, and conclusions can depend on arbitrary aggregation choices.

Here we present *Wayfarer*, a multiscale framework for spatial -omics that tracks how spatial association metrics evolve across nested spatial aggregations, enabling statistical comparison of multiscale structure across biological conditions. Using Xenium data from lung adenocarcinoma (LUAD), we show that spatial patterns often co-exist at fine and coarse scales and that progression is accompanied by reproducible shifts in scale–response profiles. These include increased fine-scale coherence of ERBB2-high tumor regions and coarse-scale clustering of immune-associated markers that are not apparent at a single resolution. *Wayfarer* converts spatial aggregation from a confounder into a diagnostic signal and is implemented as an R package to be released through Bioconductor.

## Introduction

Spatial phenomena exist in a wide range of length scales. For example, in biology, at the molecular level, protein folding is crucial to the functioning of enzymes and chromatin folding regulates gene expression. At the subcellular level, transcript localization plays a critical role in cellular function ^1,2^. At the cellular level, macrophages reside in every tissue neighboring other cell types, not only for innate immunity, but also for a variety of tissue-specific roles ^3^. At the tissue level, hepatocytes perform different metabolic roles depending on their locations relative to the portal triad and central vein ^4^. The tissues form organs which are arranged spatially in the organisms, which are then distributed spatially in their ecosystems.

As proper spatial structures at various scales are important to biological function, spatial -omics technologies have been developed to quantify biomolecules in space to gain insights to biological function. Commonly used spatial -omics technologies have a wide range of spatial resolutions. For example, in spatial transcriptomics, laser capture microdissection (LCM) can profile regions from a centimeter scale as used in the Allen Human Brain Atlas ^5^ to single cells ^6^. Spatial resolutions of sequencing-based technologies range from 100 μm between spot centers in Visium to 2 μm in Visium HD ^7^ and 0.5 μm in Stereo-seq ^8^, while in practice, Visium HD and Stereo-seq data are binned into coarser resolutions for downstream analyses ^9–11^. Imaging-based technologies such as MERFISH, Xenium, and CosMX have single cell and single molecule resolution. Meanwhile, these technologies have different capture efficiencies, which may affect the observed spatial autocorrelation of genes ^11–13^.

Despite the varying spatial scales of biological phenomena and spatial -omics technologies, most experimental studies collecting new data do not consider implications of spatial scales on downstream analyses. For example, when multiple bin sizes can be generated in Stereo-seq and Visium HD, implications of using different bin sizes is usually not considered ^10,14,15^, with some exceptions ^16^. Furthermore, when cell type co-localization is studied, co-localization is often defined by one radius or two in the same order of magnitude from each cell of interest ^17–20^, although cell type co-localization with a wider range of radii is sometimes considered ^21^.

Furthermore, these analyses are premised on cell type labels, which can be contentious due to the quality of clustering, differential expression, and cell type label transfer which are commonly used to label cells in newly collected data. The cell type label is further complicated by hierarchies of cell types and continuous variations in cell states ^22^.

Spatial scales have long been considered in geography and spatial ecology, in methods such as the correlogram ^23^, quadrat variance methods ^24,25^, Moran eigenvector spatial filtering ^26^, Ripley’s K ^27^, and multi-scale pattern analysis ^28^. In addition, modifiable areal unit problem (MAUP) has long been recognized in areal data analysis: using different spatial units to aggregate geographical data can lead to different conclusions in statistical analyses ^29^. MAUP includes the scale problem, when different spatial scales of the aggregating units impact downstream analyses. Methods have also been developed to study spatial scales in spatial -omics, such as cell communication across scales in MISTy ^30^, cell type co-localization across scales ^31–33^, and graph signal processing on the spatial neighborhood graph ^34–36^. Methods to find spatially variable genes (SVGs) that use Gaussian process (GP) models estimate the scale parameter of the GP kernel or fit multiple models with different scale parameters in order to estimate the length scale of SVGs ^37–39^.

In quadrat variance methods, observed values are aggregated by spatial quadrats or bins into longer and longer length scales and aggregated values at neighboring bins are compared for each bin size. Spatial aggregation has also been used in SERaster ^40^ to speed up computation to find spatially variable genes. This paper presents an exploratory spatial data analysis (ESDA) of lung adenocarcinoma (LUAD) data across spatial scales with spatial aggregation at different resolutions. Spatial analysis results are compared across spatial scales and LUAD stages. Having multiple patients per stage also makes it possible to distinguish between changes among stages and biological differences among patients in the same stage. This study shows the relevance of MAUP to spatial -omics and how a multi-scale analysis can convert it from a problem to an asset: First, spatial patterns at fine and coarse scales can co-exist, but the co-existence would not be apparent if the analyses are only performed at one spatial scale.

Second, some biological differences between the LUAD stages are much more pronounced in some scales than others, so the differences would be missed if the analyses are only performed at one spatial scale.

## Results

### Overview

In this study, we applied ESDA methods on Xenium data aggregated at different spatial scales to see how the results change through scales; such multi-scale behaviors are then compared across stages of EGFR-driven LUAD to gain unique insights on LUAD progression. From early to late, stages of LUAD are adenocarcinoma *in situ* (AIS) which includes Noguchi types A (AIS A) and B (AIS B), minimally invasive adenocarcinoma (MIA, Noguchi type C, or MIA C here), and invasive adenocarcinoma (IA)^41^. The IA samples are from late invasive tumors. Visium and Xenium (302 genes) data from adjacent sections^41,42^ across stages of LUAD were used in this study. To illustrate the technical implications of Visium vs. Xenium on spatial analyses, the Visium data is first spatially aligned to Xenium from the adjacent section with histology. Then the Xenium transcript spots are aggregated by the aligned Visium spot polygons to create pseudo-Visium (Figure 1A). To more systematically investigate the impact of spatial scales on spatial analyses, a square grid is created, and the number of Xenium transcript spots from each gene falling in each bin on the grid is counted (Figure 1A). For each spatial aggregation, Moran’s I, Lee’s L and MULTISPATI were performed, and how the results change with bin size or scale can be visualized as a curve for each sample (Figure 1B). As these analyses were performed on multiple samples in each stage, multiple curves were analyzed (Figure 1C-D). Changes of Moran’s I and Lee’s L through scales were compared across the stages with linear mixed models (LMM), distinguishing between variation within and between stages (Figure 1D).

**Figure 1:**
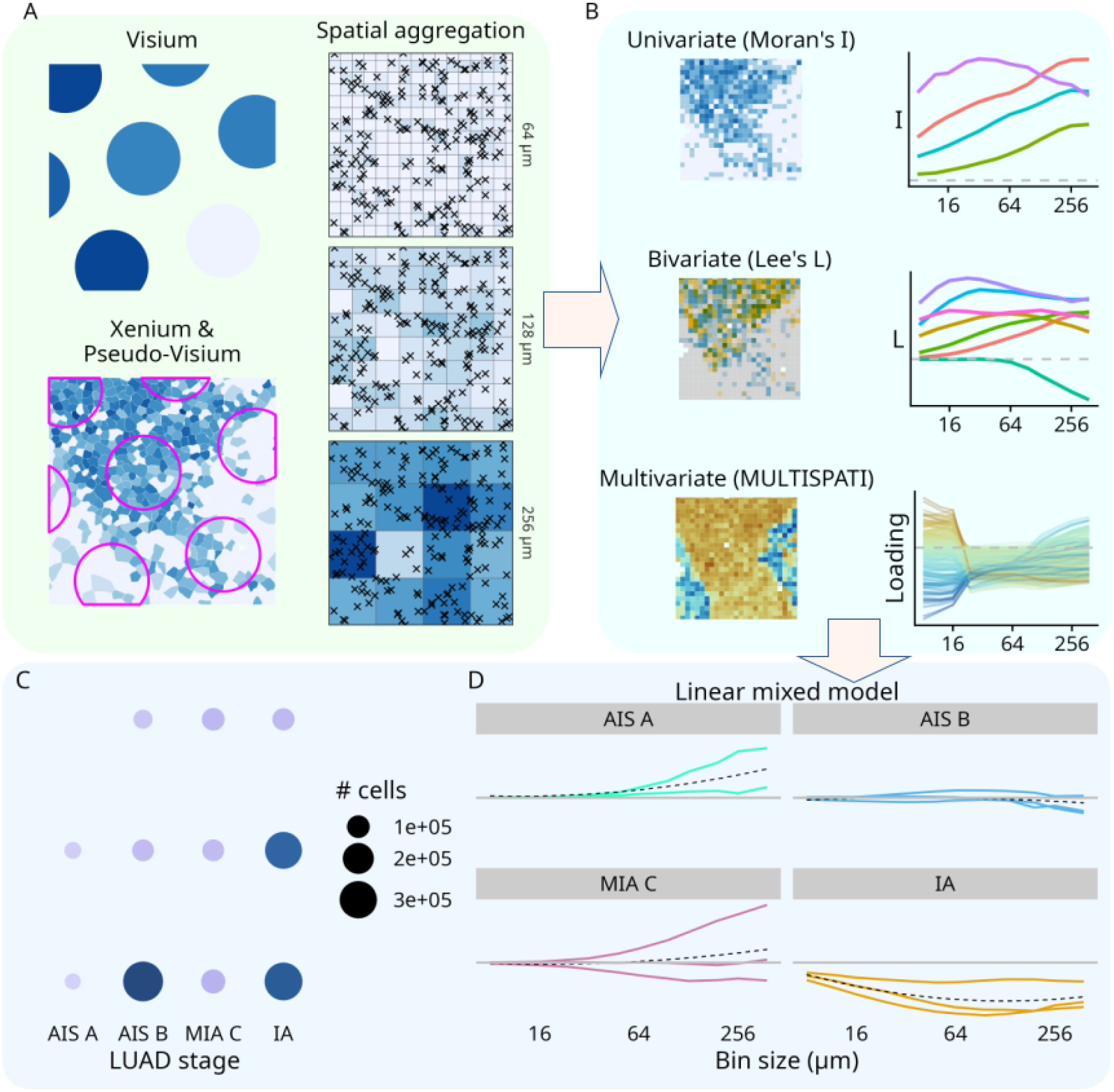
Overview of this study. A) Schematics of Visium, Xenium, pseudo-Visium, and spatial aggregation with square bins of varying sizes. B) Overview of spatial analyses performed on data aggregated with different bin sizes, including how Moran’s I, Lee’s L, and MULTISPATI change with spatial scales as shown in the curves. C) Number of cells per Xenium sample across LUAD stages; each dot corresponds to a single patient sample. D) Example of linear mixed model used to see whether Moran’s I or Lee’s L varies with bin sizes in different ways in different LUAD stages. The black dotted line shows the stage-specific spline fit which is used to determine whether the trend differs among stages.

### Moran’s I

Moran’s I is a commonly used metric of spatial autocorrelation^43^. While new methods to detect spatially variable genes in spatial -omics have been developed, Moran’s I has competitive performance compared to the new methods^44^ and is sometimes used to benchmark SVG methods^45^. To justify using Moran’s I instead of a newer SVG method for this study, we benchmarked Moran’s I against two published SVG methods, SpatialDE^37^ and nnSVG^39^. The set of genes deemed “spatially variable” is largely consistent in Moran’s I, SpatialDE, and nnSVG (Figure 2A). Moran’s I values also positively correlate with the likelihood ratio between the spatial and non-spatial models from nnSVG, indicating that Moran’s I and nnSVG agree on the magnitude of spatial variability (Figure 2B). However, with increasing numbers of bins at finer spatial resolutions, SpatialDE and nnSVG use significantly more memory than Moran’s I, limiting their utility for finer resolutions (Figure 2C).

**Figure 2:**
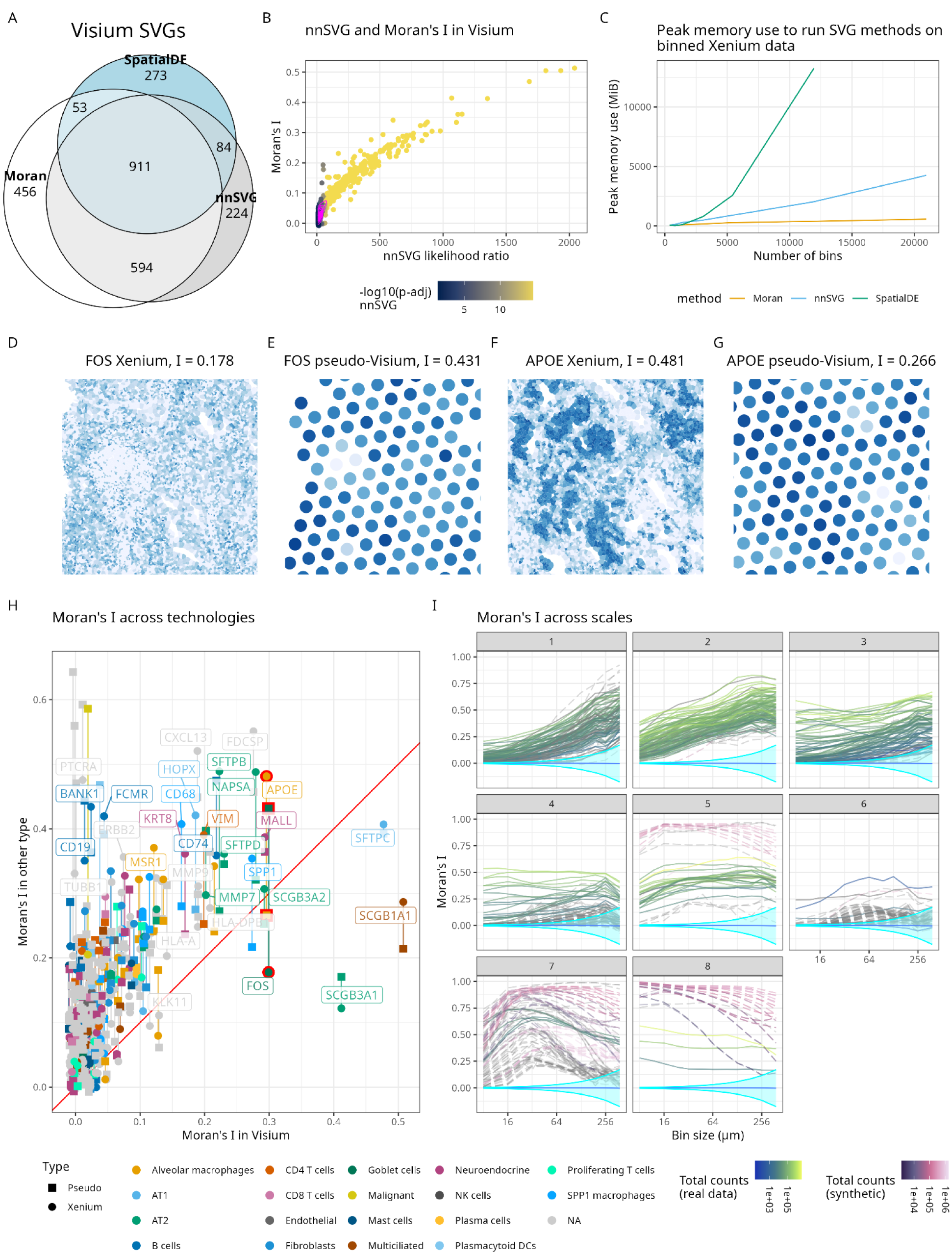
Moran’s I across technologies and scales in sample TSU-21. A) Euler diagram showing the number of SVGs common and unique to Moran’s I, SpatialDE, and nnSVG in TSU-21 Visium. B) Scatter plot of Moran’s I vs. nnSVG likelihood ratios between spatial and non-spatial models. The points are colored by -log_10_ of adjusted p-values from nnSVG. Point density is plotted in pink contours, showing that most Visium HVGs are not spatially variable according to both Moran’s I and nnSVG. C) Peak RAM used to use Moran’s I, nnSVG, and SpatialDE vs. number of bins. The SpatialDE curve ends where it used all available memory. D) Cell polygons in a small bounding box from TSU-21 are colored by log normalized expression values of FOS in Xenium, with Moran’s I=0.178. E) Visium spot polygons in the same bounding box are colored by log normalized expression of FOS in pseudo-Visium, with Moran’s I=0.431. F) Log normalized expression of APOE in a different bounding box in Xenium is plotted, with I=0.418. G) Log normalized expression of APOE in the same bounding box as in F in pseudo-Visium is plotted, with I=0.266. H) Scatter plot of Moran’s I in Xenium and pseudo-Visium vs. Moran’s I in Visium. Square points are used for pseudo-Visium and round points for Xenium, and points for the same gene are connected by a straight line. The points are colored by cell types the genes are markers of, and gray denotes genes that are not markers of the annotated cell types. FOS and APOE are highlighted with red outlines. The red line is y = x. I) Moran’s I of each gene at each bin size. Curves in green and yellow solid lines are from real TSU-21 data, and curves in purple dashed lines are from synthetic data. Each facet is a cluster based on the curves. The horizontal blue line is the expected value of Moran’s I under randomization null hypothesis, and the cyan ribbon is 2.5 and 97.5 percentiles under null hypothesis, multiplied by the number of genes in Xenium and number of bin sizes for Bonferroni correction.

Moran’s I was computed for genes detected in both Visium and Xenium in TSU-21 (AIS B). Spatial aggregation and lower Visium capture efficiency drastically impact Moran’s I. For the vast majority of genes, Moran’s I is higher in both Xenium and pseudo-Visium than in Visium (Figure 2H). This is due in part to the greater sparsity in Visium, which in some cases such as BANK1 completely obfuscated patterns clearly discernible in Xenium (Figure 2H, Supplementary Figure 1A-B). Aggregating Xenium into pseudo-Visium increased Moran’s I for some genes, such as FOS, because aggregation smoothes over local sparsity and heterogeneity (Figure 2D, E, H). However, aggregation decreases Moran’s I for some genes, such as APOE, because small-scale spatial structures get obfuscated by aggregation (Figure 2F-H).

More systematic spatial aggregations of Xenium data reveal that while aggregation increases Moran’s I for most genes (Figure 2I, clusters 1-3), some genes have Moran’s I’s that level off in a range of scales (Figure 2I, cluster 5), and some peak and decrease (Figure 2I, clusters 6 and 7). Some remain flat across scales (Figure 2I, cluster 4), while some decrease with aggregation (Figure 2I, cluster 8). To better understand these behaviors, we created synthetic data with simple patterns which recapitulate some of the behaviors observed in real data (Figure 2I). In synthetic data, each pattern was sparsified to varying extents (Methods) to illustrate the impact of sparsity on spatial analyses. In both real and synthetic data, genes that are more highly expressed tend to have higher Moran’s I (Figure 2I, Supplementary Figure 1C), and sparsifying synthetic data decreases Moran’s I (Supplementary Figures 2-3).

Both real and synthetic data reveal that when aggregation smoothes over local sparsity and heterogeneity, Moran’s I increases (Supplementary Figures 4.1). When aggregation blurs out smaller scale spatial structures and there isn’t a strong large scale spatial structure, Moran’s I decreases (Supplementary Figures 4.7). The peak indicates the scale of the spatial structure, such as spots in synthetic data and tertiary lymphoid structures (TLS) in real data; Moran’s I first increases when local sparsity and heterogeneity are smoothed over, and then decreases when bin size is exceeding the radii of the spatial structures as those structures are blurred out (Supplementary Figures 4.3, 4.5, 4.6). When Moran’s I is high and level, it means that spatial structures are present across multiple scales (Supplementary Figures 4.6, 6). This is in contrast to the single length scale parameter estimated by nnSVG.

A particularly interesting case is when the Moran’s I curve is bimodal (Figure 3). In order to understand this behavior, we created a synthetic pattern with dense small spots and sparse large spots, so only one of the scales would dominate the spatial pattern at a given bin size (Figure 3D-E). The resulting Moran’s I curve is bimodal; the peak at 32 μm highlights the small spots whose radii are randomly selected between 20 and 50 μm, and peak at 256 μm highlights the large spots whose radii are randomly selected between 500 and 1000 μm that are too sparse with smaller bins (Figure 3A, D, E). Moran’s I dropped at 64 μm when the small spots are blurred out while the large spots are too sparse. Examples in real data also show co-existence of spatial patterns at a finer and a coarser scale. For HMGA1, the first plateau after 32 μm shows some local spatial structure which becomes more blurred out at 64 μm (Figure 3C), but a different and broader spatial pattern is highlighted at 256 μm (Figure 3B). Another relevant example with bimodal Moran’s I curve is APOE, which is expressed in alveolar macrophages; higher Moran’s I at 8 μm arises from local alveolar structure, which gets blurred out with larger bins, making Moran’s I drop (Supplementary Figure 5). However, with bins larger than 64 μm, a larger scale structure across the entire section gets more highlighted, making Moran’s I rise again while the local alveolar structure is no longer visible (Supplementary Figure 5). The broader region lower in APOE is a region with alveolar collapse^41^.

**Figure 3:**
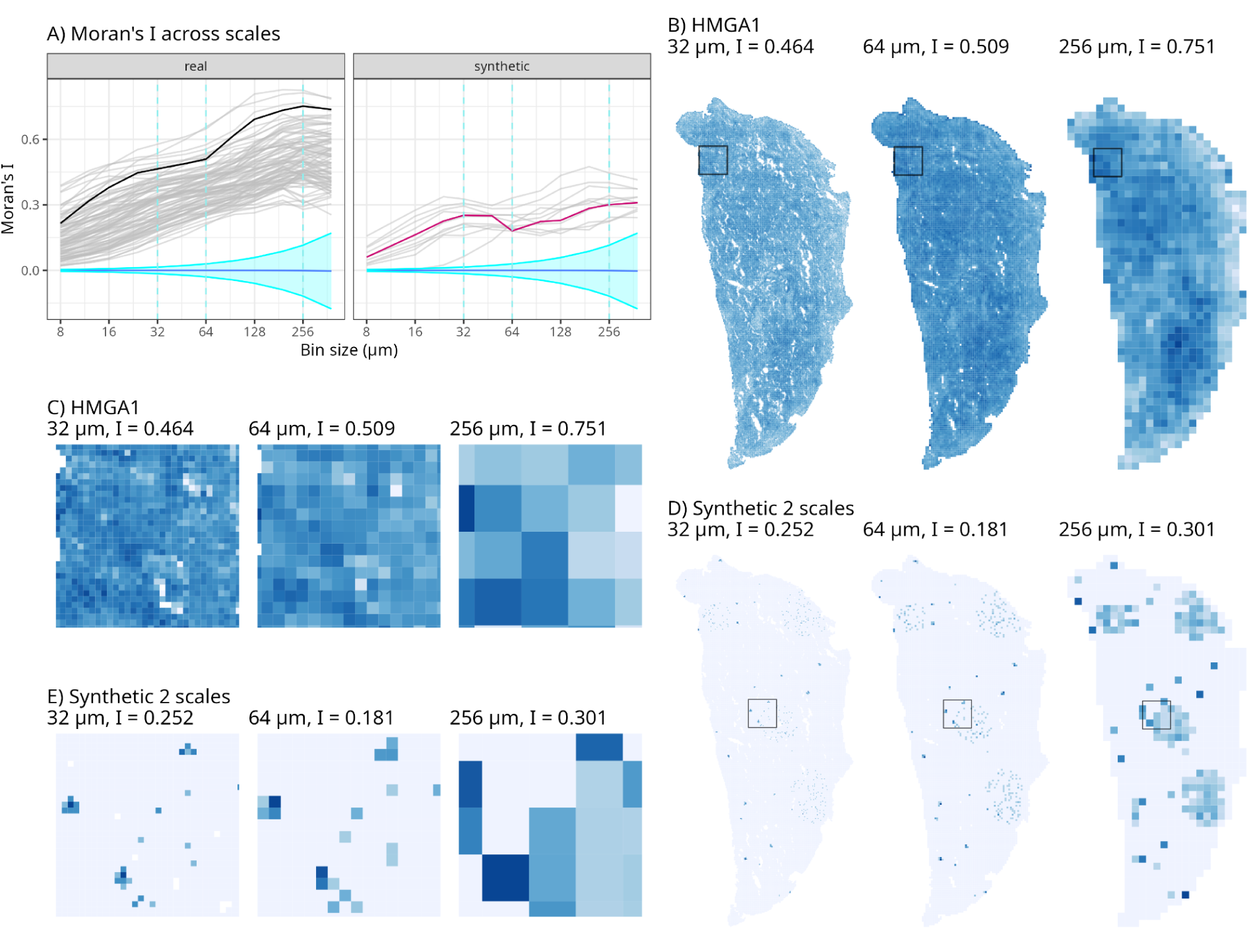
Example of co-existence of spatial structures at two different scales from cluster 2 in Figure 2I. A) Moran’s I vs. bin size for each gene; curves from real data and synthetic data are plotted in separate facets. The curves not in gray are from the features plotted spatially in panels B-E. The horizontal blue line is the expected value of Moran’s I under randomization null hypothesis, and the cyan ribbon is 2.5 and 97.5 percentiles under null hypothesis, multiplied by the number of genes in Xenium and number of bin sizes for Bonferroni correction. B) Global pattern of a gene from real data, plotted at bin sizes 32, 64, and 256 μm. C) Zooming into the box shown in B. D) Global pattern of a synthetic feature, plotted at bin sizes 32, 64, and 256 μm. E) Zooming into the box shown in D.

### Lee’s L

Lee’s L is a spatially informed correlation metric that combines Moran’s I and Pearson correlation. Two variables can attain high Lee’s L only when they are both spatially structured and correlated^46^. Lee’s L was computed for all pairwise combinations of genes detected in both Visium and Xenium. As in Moran’s I, Lee’s L tends to have greater magnitude in Xenium and pseudo-Visium compared to Visium, and in pseudo-Visium compared to Xenium (Supplementary Figure 7). An example that illustrates this tendency is HMGA1 and PTCRA. HMGA1 is involved in cancer progression^47,48^, and PTCRA plays a critical role in αβ T cell development^49,50^. In Visium, both genes are sparsely detected and they are not widely co-detected in the same spots, so their Lee’s L is close to 0 (Supplementary Figure 7). In contrast, in pseudo-Visium, the two genes are far less sparse and are more commonly co-detected, giving a higher Lee’s L (Supplementary Figure 7). In Xenium, while the two genes are commonly co-detected in the same cells, many cells have high expression of one but not the other, leading to a lower Lee’s L than in pseudo-Visium, but aggregating these cells with neighboring cells that has higher expression of the other genes increases Lee’s L in pseudo-Visium (Supplementary Figure 7).

To explore how Lee’s L varies across a wider range of spatial scales, we computed Lee’s L for all pairwise combinations of Xenium genes though only gene pairs with Lee’s L greater than 0.2 in magnitude in at least one bin size are analyzed. These Lee’s L curves were clustered in a similar manner as the Moran’s I curves (Supplementary Figure 8, Methods). Across scales, Lee’s L may monotonically increase from near 0 or from a higher value, peak and decrease, monotonically decrease from 0 to a negative value or from a more positive value to a value closer to 0, or remain flat (Supplementary Figures 8, 9). Having two variables complicates behaviors of Lee’s L curves. To better understand these behaviors in Lee’s L, a multivariate synthetic dataset with two “gene programs” was constructed and Lee’s L was computed across scales for each pair of features (Supplementary Figure 10). Features in the “gene programs” have various strengths of correlation, and 8 μm bins were assigned to each gene program in different spatial patterns (Methods).

In synthetic data, as in Moran’s I, increasing sparsity decreases Lee’s L across scales (Supplementary Figure 10). As in Moran’s I, spatial aggregation brings cells together in sparse regions, highlighting spatial patterns of co-expression at larger scales; in both real and synthetic data, when one or both of the features are sparse but a larger scale pattern is present, the magnitude of Lee’s L increases with spatial aggregation (Supplementary Figures 9.1A-C, 10.3). However, when either feature is sparse and a negative correlation exists at a larger scale, Lee’s L increases in magnitude with spatial aggregation but becomes more negative (Supplementary Figure 9.4). As in Moran’s I, when both genes co-express in a spot-like pattern as in TLS, the Lee’s L curve shows a peak (Supplementary Figure 9.5, 9.6, 10.3).

An interesting pattern is when Lee’s L remains flat across different bin sizes (Figure 4). Upon inspection, different spatial structures of co-expression are observed at small, medium, and large scales (Figure 4). As in Moran’s I, bimodal curves have been observed, such as in APOE-CD68, indicating co-existence of different spatial co-expression structures at different scales (Supplementary Figure 9.1A, D, E), and the same synthetic dataset that produced bimodal Moran’s I curves produced bimodal Lee’s L curves for genes in the same gene program (Supplementary Figure 10.1). The bimodality can also be inverted, when negative correlation between two genes occurs at two different length scales but less so at an intermediate scale (Supplementary Figure 9.2A, D, E).

**Figure 4:**
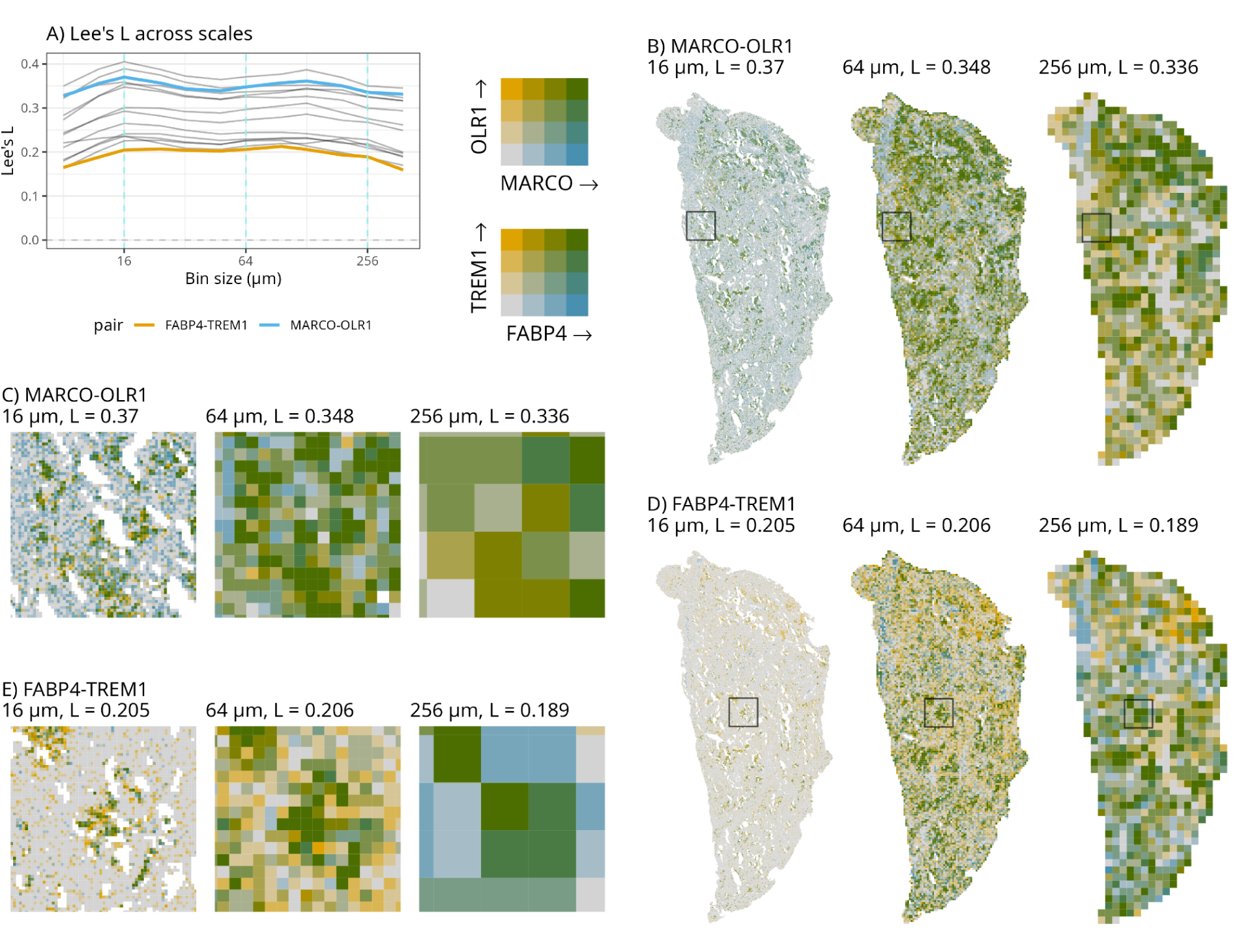
Example of co-existence of spatial structure in gene co-expression, from cluster 7 in Supplementary Figure 8. A) Lee’s L vs. bin size; curves for two example gene pairs are highlighted. Vertical dashed lines mark bin sizes where examples are plotted spatially in B-E. B) Spatial plot of the first example (MARCO-OLR1) for the whole section of TSU-21. C) Zooming into the box marked in B. D) Spatial plot of the second example (FABP4-TREM1) for the whole section of TSU-21. E) Zooming into the box marked in D. In the bivariate palette, blue bins are high in gene 1 and low in gene 2, yellow bins are high in gene 2 and low in gene 1, green bins are high in both, and gray bins are low in both.

### MULTISPATI PCA

Since spatial aggregation affects detection of gene co-expression, it may lead to different results in multivariate analyses as well. MULTISPATI PCA is a method of spatially informed PCA, derived from the matrix expression of Moran’s I^51,52^. While in PCA, the eigenvalues are variance explained, in MULTISPATI, the eigenvalues are the product of Moran’s I and variance explained. In TSU-21, with smaller bin sizes (8-16 μm), PC1 separates alveolar and cancer cells from immune cells (Supplementary Figure 11A). However, there is a “phase shift” at 24 μm, where the meaning of PC1 becomes very different (Supplementary Figure 11C-D). This is because bins 24 μm or larger bring neighboring alveolar and immune cells that are separate in Xenium and smaller bins together into the same bins (Supplementary Figure 11B), so differences between them is no longer variability between bins that PCA maximizes. Similar “phase shifts” are observed in other PC’s of TSU-21 and in other samples (Supplementary Figure 11C, Supplementary Figure 12). However, it’s not straightforward to compare these results across LUAD stages.

### The hidden progressions of LUAD: spatial differences between stages

So far the analyses have been performed in one sample, TSU-21. We ask whether the way Moran’s I and Lee’s L vary across spatial scales may differ between LUAD stages. For each gene or gene pair, there is one curve for each sample, and we ask whether these curves differ between stages. So we fitted a LMM with spline terms to model these curves. The random effects include a random intercept and a random slope for the spline terms; each stage has its own intercept and slope, so each stage has its own “average” curve between the individual patients (Figure 1D). To see if the curves differ among stages, either the entire random effect term or only the random slope was dropped to create a reduced model. The full and reduced models were then compared with a likelihood ratio test to see if the random effect or random slope is significant, to determine whether the curve differs among stages (see Methods for details).

While differences between patients within the same stage are commonly observed, over 80% of Xenium genes have significant random effect in Moran’s I (Supplementary Table 1) and over 70% of all gene pairs have significant random effect in Lee’s L (Supplementary Table 2). This is not surprising given that the Xenium panel was curated to include genes relevant to LUAD. Considering that sparsity decreases Moran’s I and Lee’s L as discussed earlier, we asked whether different expression levels across stages confound differences in Moran’s I curves. We performed pseudo-bulk differential expression (DE) and found that most genes with significant random effects or random slopes in Moran’s I are not significant in pseudo-bulk DE (Figure 5A, Supplementary Table 3), suggesting that there are spatial differences across scales in most genes across stages that are not due to overall expression levels. Examples from genes of special interests to LUAD are plotted in Supplementary Figures 13 and 15; all of them have significant random effects and some have significant random slopes. Only SPP1 is significantly DE between stages in pseudobulk (Supplementary Table 3, Supplementary Figure 14), and its increased expression level in IA is associated with increased Moran’s I in IA compared to earlier stages (Supplementary Figures 13, 15.1-2).

**Figure 5:**
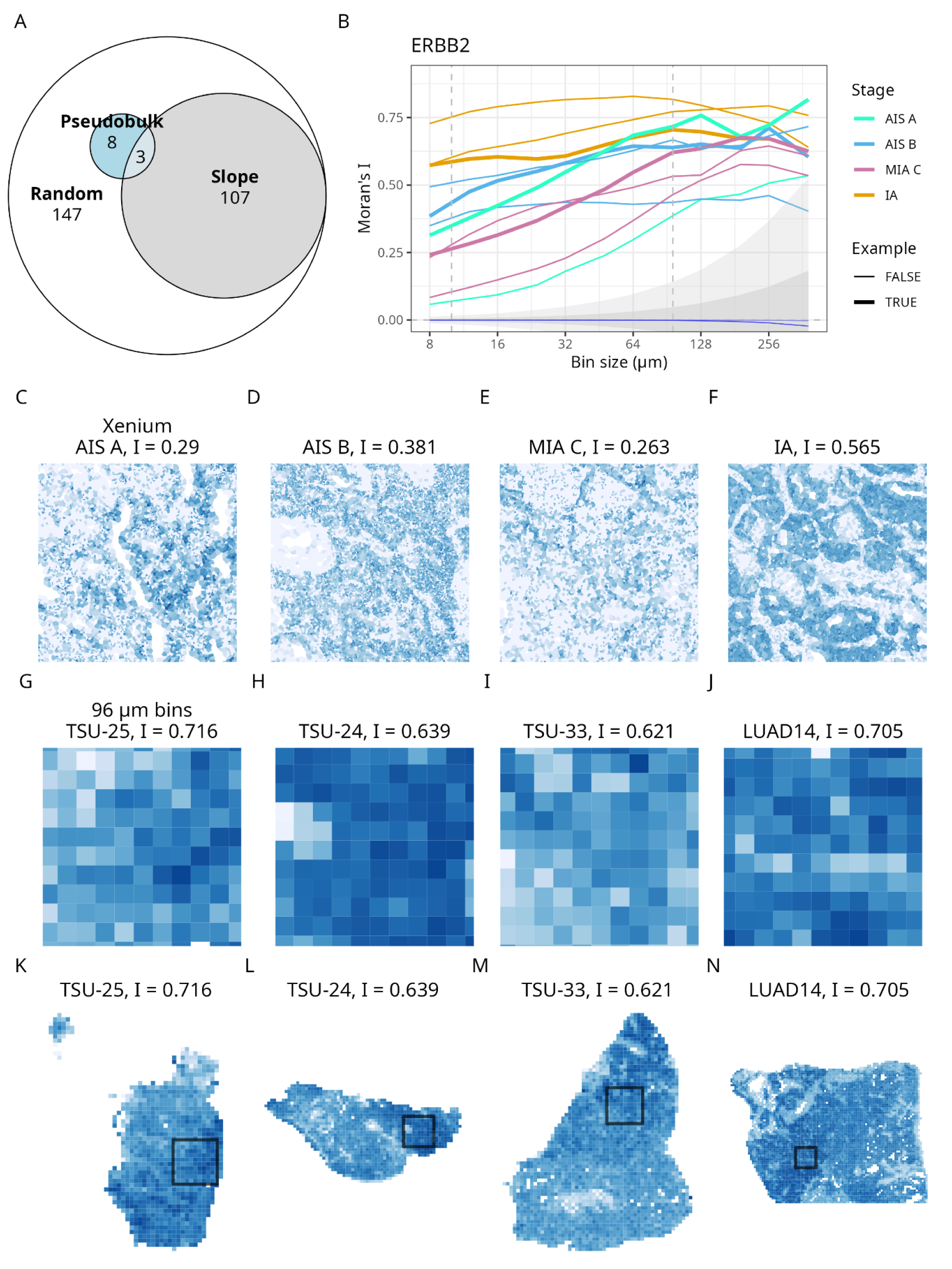
Moran’s I curves differing across LUAD stages. A) Euler diagram showing the number of genes significant in pseudobulk DE between stages (Pseudo), genes with significant random effects (Random), and genes with significant random slopes (Slope). B) Moran’s I curves for ERBB2 as an example; the curves are colored by LUAD stage, and thicker curves correspond to samples shown in panels C-N. Vertical dashed lines mark 10 μm (approximately the size of Xenium cells) and 96 μm which are shown in examples in C-N. The blue line is the expected value of Moran’s I under randomization null hypothesis in the sample with the fewest cells (TSU-20). The light ribbon is the 2.5 and 97.5 percentiles under null hypothesis, multiplied by the number of genes in Xenium, number of bin sizes, and number of samples for Bonferroni correction computed from TSU-20. The dark ribbon shows the same percentiles from the sample with the most cells (TSU-21). C-F) Local expression patterns of ERBB2 at single cells in Xenium in 1000 μm x 1000 μm boxes in the samples highlighted in B. Moran’s I values are for Xenium single cells. G-J) Same as C-F but in 96 μm bins. Moran’s I values are for 96 μm bins.K-N) Plot of whole sections for samples shown in C-J, showing 96 μm bins. The boxes mark the 1000 μm x 1000 μm regions shown in C-J.

We use the example of ERBB2 to illustrate spatial differences across stages. ERBB2 is a driver of cancer progression and the ERBB/HER family is targeted in cancer therapy^53^. In pseudo-bulk DE, ERBB2 is not significant after correcting for multiple testing (adjusted p = 0.138), but has highly significant random effect (adjusted p = 1.92e-9) and random slope (adjusted p = 7.98e-5) in Moran’s I, which means Moran’s I changes with scale in different ways in different stages. At finer scales, Moran’s I is much higher in IA than in earlier stages, but the difference shrinks at coarser scales (Figure 5B). At the single cell resolution, the earlier stages have many cells with high expression of ERBB2, but there’s more local heterogeneity and many cells with low ERBB2 co-localize with those with high expression (Figure 5C-E). In contrast, in IA, cancer cells highly expressing ERBB2 form coherent and homogeneous blocs (Figure 5F). With 96 μm bins, at a resolution comparable to Visium, this difference is not observed and all samples have regions with higher ERBB2 expression (Figure 5G-N). In the original publication of this dataset, Visium data was first analyzed in depth and Xenium data was later used to validate findings from Visium^41,42^; analyzing Visium first might miss this spatial difference across stages.

The observation of significant random slopes for T cell associated markers, specifically ITGAE and GZMB, and immunotherapy marker CXCL9 reveals a fundamental shift in the immune geography of LUAD as it transitions from in situ to invasive stages (Supplementary Figure 13). Biologically, ITGAE (CD103) is a hallmark of tissue-resident memory (TRM) T cells, which are essential for localized antitumor surveillance. At the single cell level, Moran’s I of ITGAE is minimal in all stages, but at a larger scale, Moran’s I is higher in IA than in earlier stages. This suggests that while TRM cells may be intimately intermingled with malignant cells at the single-cell level in early AIS or MIA stages, they become spatially restricted in IA stages (Supplementary Figures 13, 15.3-4). However, this behavior can also be due in part to scarcity of TRM T cells in the tissue. Similarly, the shifting slope of GZMB suggests that active cytotoxicity zones transition from broad, fine-scale infiltration to localized, coarse-scale pockets of activity at the boundary of invasive regions. Cytotoxicity is largely excluded from the invasive regions, potentially reflecting the formation of the coherent, immune-excluded blocs observed in invasive tumors (Supplementary Figures 15.5-6). For CXCL9, a chemokine that attracts T cells and has been associated with better prognosis in cancer^54^, Moran’s I exhibits an opposite stage-dependent trend. CXCL9 has moderate Moran’s I in all but one sample at fine scales. With spatial aggregation, Moran’s I decreases in pre-invasive stages, reflecting diffuse CXCL9 expression pattern, but increases in IA, due to clustering at invasive boundaries (Supplementary Figures 13, 15.7-8). This shift from diffuse to boundary-localized clustering is consistent with T cell exclusion shown in the spatial pattern of GZMB and ITGAE in IA.

From a therapeutic standpoint, these multi-scale signatures suggest that T-cell density alone may be an insufficient biomarker for predicting immunotherapy response. In invasive stages where GZMB and ITGAE exhibit purely coarse-scale patterns, the therapeutic challenge shifts from simply activating T-cells to overcoming the physical and spatial barriers that prevent fine-scale infiltration into the tumor core. This highlights the potential for using spatial slope as a biomarker to identify patients who may require combination therapies aimed at normalizing the tumor microenvironment to restore single-cell level immune-tumor interactions.

Beyond T-cell exclusion, the multi-scale analysis reveals a significant restructuring of the suppressive myeloid landscape across LUAD stages. SPP1 is a gene frequently associated with pro-tumorigenic, ’M2-like’ macrophages. As expected, SPP1 is more highly expressed in IA than in earlier stages and more lowly expressed in AIS A; such difference in expression level corresponds to higher Moran’s I in IA (Supplementary Figures 14, 15.1-2). Furthermore, the transition of MDSC-related markers such as ITGAM (CD11b) and PTPRC (CD45) across scales highlights the evolving nature of the pre-metastatic niche. However, for some myeloid markers such as CD68 and CD163, the significant random effects seem to be driven by lower Moran’s I in MIA C, and it is confounded by increased technical sparsity in two MIA C samples which lowered Moran’s I at fine scales (Supplementary Figures 14, 16).

### Gene conversations change with distance

As most gene pairs have significant random effects, we used CellChat database (CellChatDB)^55^ and genes of interest to narrow down to cases that may be biologically relevant. Twenty-five gene pairs with significant random effects are known interactions in CellChatDB, and four KEGG pathways are represented: ERBB signaling (hsa0401), cell adhesion molecule interactions (hsa04514), extra-cellular matrix (ECM)-receptor interaction (hsa04512), cytokine-cytokine receptor interaction (hsa04060), and neuroactive ligand-receptor interaction (hsa04080) (Supplementary Table 4, Supplementary Figures 17-18). However, since the co-expression here is not only at the level of cells, but also at the level of cellular neighborhoods, gene pairs that are not known molecular interactions can still be biologically relevant as they may arise from co-localization of cells playing different roles.

An example that shows the relevance of multi-scale analysis is ERBB2 and PRG4. PRG4 encodes an ECM protein proteoglycan 4. It has been shown to inhibit cancer progression and is associated with better cancer prognosis^56,57^. At a fine scale, Lee’s L is very low in magnitude in all samples, because PRG4 is sparsely distributed (Figure 6A, F-I). The low magnitude would most likely not bring attention to this co-expression at the single cell level. However, stronger co-expression becomes apparent with spatial aggregation; while the earlier stages show positive correlation, negative correlation becomes more apparent in IA (Figure 6A-E). Aggregated data shows a large region of ERBB2-high cancer cells excluding PRG4 while earlier stages have larger regions of co-expression (Figure 6B-E). At the single cell level, cells expressing PRG4 seem to intermingle with ERBB2-high cancer cells in earlier stages, while in IA, ERBB2-high regions exclude PRG4 (Figure 6F-I). This is consistent with the known properties of PRG4. Note that one of the IA samples (LUAD17) goes in a different direction from the other two in many co-expressions (Figure 6A, Supplementary Figure 14). While an analysis at a Visium-like resolution may pick up this difference, with the rise of Visium HD which is often analyzed as 8 μm bins and is much sparser than Xenium, this difference may be missed if analysis is not also performed with coarser aggregation.

**Figure 6:**
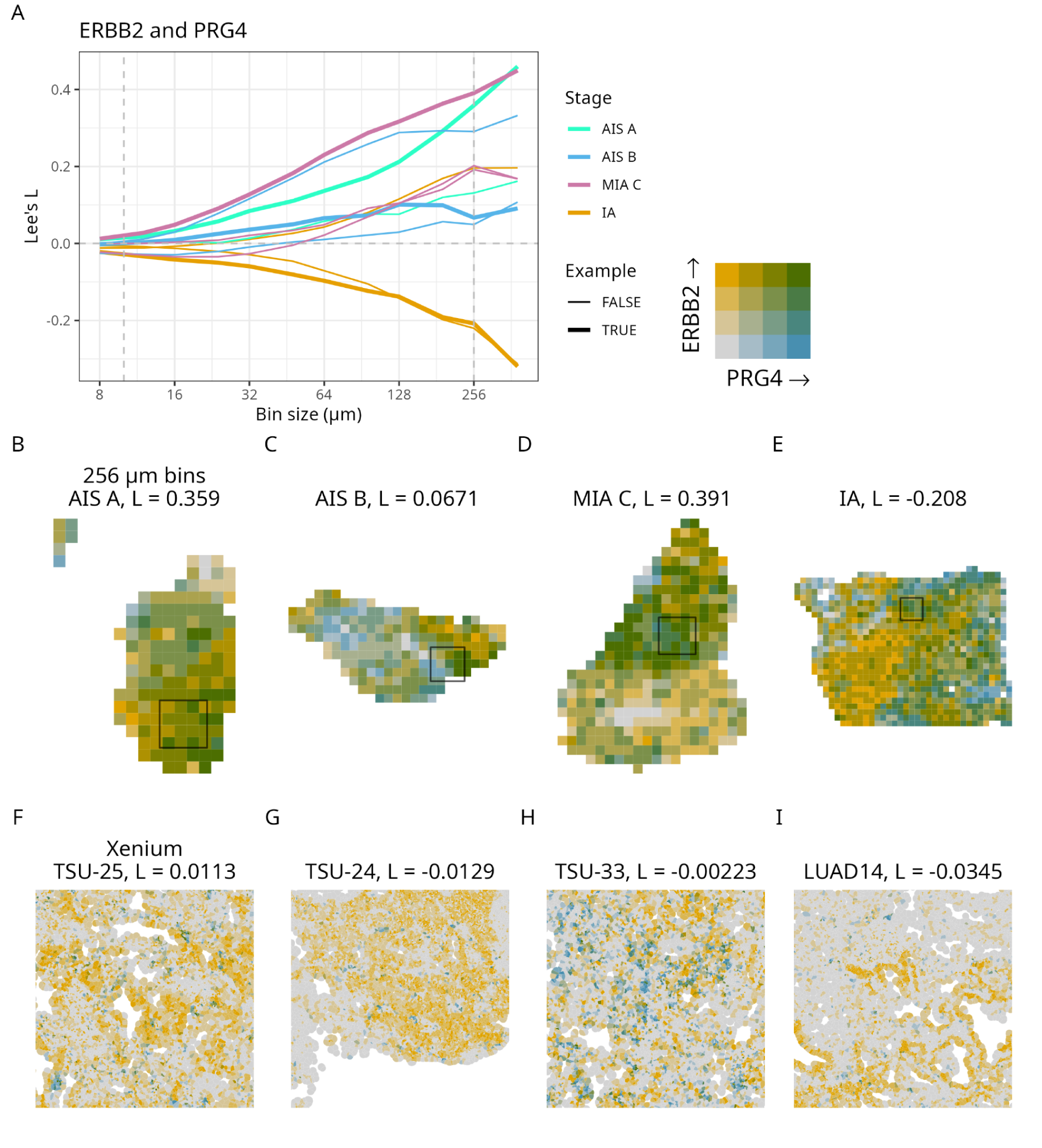
Lee’s L curves differing across LUAD stages. A) Lee’s L curves for ERBB2 and PRG4 as an example; the curves are colored by LUAD stage, and thicker curves correspond to samples shown in panels B-I. Vertical dashed lines mark 10 μm (approximately the size of Xenium cells) and 96 μm which are shown in examples in B-I. B-E) Plot of whole sections of samples highlighted in A, showing co-expression of the two genes in 96 μm bins. Boxes mark 1000 μm x 1000 μm regions shown in F-I. Lee’s L values are for 96 μm bins. F-I) Expression of the two genes in Xenium cells in 1000 μm x 1000 μm regions. Panel F corresponds to B, G corresponds to C, and so on. Lee’s L values are for Xenium single cells.

We also investigated some gene pairs associated with immunosuppression and T cell exclusion, to further expand on the observations on Moran’s I. APOE+ macrophages also contribute to an immunosuppressive tumor microenvironment. In clear cell renal carcinoma (ccRCC), APOE+ macrophages are found to form a physical barrier between the malignant region with high SPP1 expression and the region with CD8+ T cells in Visium, and that tumor-derived SPP1 drives APOE+ macrophages to establish an immunosuppressive tumor microenvironment ^58^. In LUAD, SPP1 and APOE exhibit weak spatial correlation at a fine scale with Lee’s L increasing with progression from AIS to MIA and IA, but the correlation increases with spatial aggregation in most pre-invasive samples and decreases in two IA samples (Supplementary Figures 19, 20.1-2). In IA, SPP1 is highly expressed in invasive regions while APOE is more expressed outside these regions, consistent with the observation in ccRCC in^58^. However, the spatial co-localization of SPP1 and APOE at a fine scale in all stages and the transition from positive to negative correlation from MIA to IA suggest further roles of SPP1 and APOE in establishing an immunosuppressive tumor microenvironment over the course of LUAD progression.

Spatial correlations between SPP1 and CXCL9, ITGAE, and GZMB highlight T cell exclusion in IA but less so in pre-invasive stages and the importance of follow-up analysis with local spatial statistics (Supplementary Figures 19-21). For T cell markers ITGAE and GZMB, Lee’s L with SPP1 is minimal in all stages at a fine scale, and a a broader scale, Lee’s L tends to be lower in IA than in earlier stages due to T cell exclusion from the invasive regions in IA (Supplementary Figures 19, 20.5-8). In earlier stages, cancerous regions do not exclude ITGAE (TRM T cells), resulting in moderate Lee’s L (Supplementary Figure 20.6), but exclusion of GZMB (cytotoxic T cells) is found in some AIS B and MIA C samples but none of the AIS A samples (Supplementary Figure 20.8). For CXCL9, at a fine scale, co-localization with SPP1 is not apparent in pre-invasive stages and exclusionary zones are apparent in IA upon visual inspection (Supplementary Figure 20.9). However, at a coarse scale, AIS B and IA seem to have comparable Lee’s L values despite visually different spatial patterns of correlation (Supplementary Figure 20.10). This is because in IA, there is a component of positive correlation between SPP1 and CXCL9, as CXCL9 is highly expressed at the boundaries of invasive regions high in SPP1 (Supplementary Figure 20.10); the boundaries and regions where both are low contributed to a positive Lee’s L value. Local Lee’s L and local bivariate Moran’s I show each bin’s contribution to the global spatial correlation; they show significant regions of exclusion as well as regions where both SPP1 and CXCL9 are low in IA, but only significant regions of co-expression in AIS B (Supplementary Figure 21.1). Co-existence of local positive and negative spatial correlation between SPP1 and GZMB in IA also explains why Lee’s L is not more negative at a large scale, and AIS A similarly shows only regions of positive correlation that are significant (Supplementary Figure 21.2). Co-existence of local positive and negative correlation canceled out in NAPA and VIM in TSU-21, leading to a Lee’s L near 0 (Supplementary Figure 22).

## Discussion

In this study, we demonstrated the relevance of analyzing spatial transcriptomics data across multiple spatial scales. First, we showed how the lower capture efficiency and spatial resolution of Visium compared to Xenium impact spatial patterns, consistent with the comparison between Visium (with vs. without CytAssist) and Xenium in^11^. CytAssist was not mentioned in^41,42^; not using CytAssist might have negatively impacted the quality of Visium data here. Because sequencing based technologies are transcriptome-wide but mostly don’t have single cell resolution, while imaging based technologies have single cell resolution but have limited gene panels, the two types of technologies are often used in conjunction as they complement each other. For example, imaging based spatial transcriptomics is often used to verify findings from sequencing based spatial transcriptomics regarding spatial niches^16,41,42,59,60^. In addition, with single cell resolution, imaging based data is often used to characterize cell type co-localization in spatial niches^17–21^. With transcriptome-wide coverage, sequencing based data is sometimes used to quantify ligand-receptor interactions ^41,59^ and copy number variation (CNV) in cancer^18,60^.

Second, we showed that spatial patterns at a finer scale can co-exist with spatial patterns at a coarser scale, both in individual genes and gene co-expression. This means that there might not be one “optimal” spatial binning or resolution with which to perform the analyses, as suggested in some methods proposed to mitigate MAUP ^61^. The “proper” resolution would depend on the spatial scale relevant to the question of interest, which can be single cells or the level of neighborhoods. Third, we found multi-scale differences between LUAD stages that may not be apparent at a single scale. In summary, performing spatial analyses across multiple scales can lead to insights missed if the analyses were performed at a single scale. Visium HD, Stereo-seq, and Xenium 5k bring simultaneous single cell resolution and transcriptome wide coverage within reach. This study suggests that the lower capture efficiency of Visium HD can obscure spatial autocorrelation, which is confirmed in ^11^. Applying the kind of analysis performed on this study to Visium HD, Stereo-seq, and Xenium 5k can enable further insights that are impractical with a panel of 302 genes, such as gene set and pathway enrichment in different types of multi-scale behaviors. If survival information is available, multi-scale spatial patterns may be associated with survival.

A limitation of spatial binning is that edge effect has led to spurious Moran’s I for densely expressed features, i.e. a spatial pattern caused by bins that are not fully covered by tissue, that are at the edge or holes (Supplementary Figure 23B-H). The Moran’s I and Lee’s L curves might not be easy to interpret, as different spatial patterns with similar spatial scales can give rise to similar curves. Furthermore, the curve is a global summary over the entire tissue section, obscuring local regions that can have different spatial patterns. Hence this analysis should be complemented with local Moran’s I and Lee’s L to understand local contributions to these metrics and potentially other local spatial methods such as geographically weighted PCA^62^. In addition, the LMM might not accurately model dependency of Moran’s I or Lee’s L between bin sizes as the binned data were created from the same original transcript spots. Hence the LMM results should be considered exploratory. To ensure that inferred stage-specific slopes reflect stable biological signal rather than artifacts of spatial resolution or preprocessing, additional robustness and consistency checks are needed. However, formulating an appropriate null hypothesis remains challenging for structured spatiotemporal data^63^. Future studies could formulate a conditional null hypothesis using knockoff-based significance testing^64^, by comparing observed stage-specific slopes to those obtained under spatially constrained null transformations that preserve sampling geometry and multiscale structure while breaking biological associations. Finally, a multi-condition, representation learning framework such as Patches^65^ may also be adapted to multi-scale spatial data to find gene programs common and unique to cancer stages.

## Methods

### Data acquisition

Processed paired Visium and Xenium data from ^41,42^ were downloaded from https://kero.hgc.jp/Ad-SpatialAnalysis_2024.html, as Space Ranger output. For IA, only data from fresh frozen samples were used, in order not to introduce technical differences between IA samples and samples from other stages for which all samples are fresh frozen. For AIS type A, TSU-23 was not used for spatial analyses as it appears to be multiple needle biopsies that would complicate downstream spatial analysis.

### Visium analysis

All analyses were performed with R 4.5.0 and Bioconductor 3.22 running on Ubuntu. All plots were made with ggplot2 (v3.5.2.9001)^66^ and multi-panel plots were assembled with patchwork (v1.3.1)^67^ unless otherwise indicated. Visium data was only analyzed for TSU-21, from the stage AIS Noguchi type B. Processed filtered Visium data was read into R as a SpatialFeatureExperiment (SFE, v1.11.1) object and a spatial neighborhood graph of adjacent spots was created in the read process ^68^. Then the SpotSweeper (v1.5.0) package was used to remove Visium spots that consistently have low library size across datasets due to their spot barcode sequences ^69^. Spots with no spatial neighbors were also removed. Genes detected in fewer than 5 spots were removed.

Visium data was normalized with logNormCounts() from the scran package (v1.37.0) ^70^. Highly variable genes (HVGs) were identified with scran’s getTopHVGs(), with default parameters. Non-spatial PCA was performed with runPCA() of scater (v1.37.0) ^71^, with scale = TRUE and using the highly variable genes. Moran’s I, Lee’s L, and MULTISPATI PCA were run with the Voyager package (v1.11.0)^72^, on log normalized data. To compare Moran’s I with published SVG methods, we ran nnSVG (v1.13.1) and SpatialDE (via the Bioconductor package spatialDE v1.15.1) with log normalized HVG data.

For Figure 2A, the list of “significant” SVGs are the genes with adjusted p-values < 0.05 from SpatialDE and nnSVG output. For Moran’s I, spdep::moran.test() was used to obtain the mean and variance of Moran’s I under randomization null hypothesis given TSU-21 Visium’s spatial neighborhood graph. The mean and variance were used to compute z-scores for observed Moran’s I values for each gene. Because negative spatial autocorrelation has not been observed here, a one tailed test for positive spatial autocorrelation was performed on Moran’s I based on the z-score, and the p-values were adjusted for multiple testing with the Benjamini-Hochberg method. Genes with adjusted p < 0.05 were included in the list of SVGs according to Moran’s I. The eulerr R package (v7.0.2)^73^ was used to make the Euler diagram.

To profile memory usage as a function of number of bins, Moran’s I (Voyager and spdep), nnSVG, and SpatialDE were run on binned TSU-21 data, until SpatialDE ran out of memory because a dense covariance matrix among the bins was made. The peakRAM R package was used to profile peak RAM usage. As SpatialDE was run through reticulate, Python’s resource.getrusage(resource.RUSAGE_SELF).ru_maxrss was used to get peak RAM usage.

After quality control (QC) and basic analyses, the SFE object that holds analysis results was saved to disk with the alabster.sfe package (v1.1.0)^68^. Lee’s L results as a correlation matrix were saved as a separate RDS file.

### Xenium analysis

Xenium QC and basic ESDA were performed on all downloaded Xenium datasets. Processed Xenium data was read into R with SpatialFeatureExperiment; transcript spots were not read into memory. Because of the small gene panel with 302 curated genes that does not cover markers of cell types not of interest, many cells with clearly visible nuclei staining and good segmentation have low transcript count. Therefore only cells with 0 detected transcripts were removed. Cells with over 60% spot counts from negative control probes were also removed. Negative control probes were removed before downstream analysis.

Cells and small clumps of cells far from tissue were considered debris and also removed; such clumps have at most 20 cells and are at least 50 micrometers from the main piece of tissue, although the number of cells has been adjusted to remove obvious debris that appear larger. The clumps were detected by constructing a distance-based graph where cells whose centroids are within a distance cutoff are considered neighbors. The distance cutoff is the minimum distance from the main piece of tissue (here 50 microns). This was done with BiocNeighbors (v2.3.1)^74^, and implemented as a new function findDebrisCells() in SpatialFeatureExperiment. Then n.comp.nb() in spdep (v1.3.13)^75^ was used to identify which cells are in which connected component in the graph. The number of cells in each component is the number of cells in each piece of tissue, and in each “piece”, all cells are further from other pieces than the distance cutoff. If the number of cells is lower than a threshold, then this piece is considered debris.

Tissue boundaries were found with concave hulls of cell segmentation polygons of cells that passed QC, with st_concave_hull() in the sf package (v1.0.21) ^76^, and holes are allowed. This seems to work better than segmenting the tissue from DAPI staining as this dataset is from Xenium v1 for which only DAPI nuclei staining is available, without cell membrane stains.

Xenium data was normalized with logNormCounts(), but with cell area instead of total counts as size factors, as suggested by ^77^. PCA was run as in Visium, but with all genes. The cells were clustered using the first 20 PCs with Leiden as implemented in bluster (v1.19.0)^78^, with objective function “modularity” and resolution 0.8. A k-nearest neighbor graph with k=5 was used for spatial analyses for Xenium. Moran’s I, Lee’s L, and MULTISPATI were performed with all genes, using log normalized data. The updated SFE object after QC and holding ESDA results was saved to disk with alabaster.sfe.

### Comparison of Visium and Xenium

The alignment and comparison between Visium and Xenium was only performed for TSU-21, as this is sufficient to demonstrate the effects of spatial binning and sparsity on spatial analyses. Furthermore, for many other samples, the Visium and Xenium sections appear very different despite still appearing to come from nearby sections, which makes alignment difficult.

The EBImage package (v4.51.0)^79^ was used to find tissue boundaries from images. Tissue boundary was segmented from the “hi-res” image from Visium Space Ranger output and from Xenium DAPI image (morphology_focus, resolution 6). For Xenium, contrast is enhanced with CLAHE. Then Otsu thresholding was used to find a mask indicating the tissue, and the mask was converted to polygon with as.polygon() in terra (v1.8.60)^80^, which is then converted to sf and simplified with st_simplify(). For Xenium, morphological closing was applied to remove debris and holes before the mask was converted into a polygon. This was added to SpatialFeatureExperiment as a new function getTissueBoundaryImg().

After the polygon is simplified, regions with more complicated shapes have more vertices, so the complicated regions are used to align the Visium and Xenium polygons. Vertices of the Visium and Xenium polygons are aligned with affine transformation. Point cloud registration as implemented in the LOMAR package (v0.5.0)^81^ was used to find a transformation matrix and translation vector, which are then applied to the Visium vertices to align them to the Xenium ones. The same transformation was applied to Visium spot polygon vertices to align them with Xenium.

To make pseudo-Visium, the aggregateTxTech() function in SpatialFeatureExperiment was used to aggregate Xenium transcript spots with the aligned Visium spots. For each gene, the number of transcript spots that intersect with each Visium spot is counted to make the gene count matrix. Then basic ESDA was performed on pseudo-Visium in the same way as Visium.

In order to give a fair comparison between Visium and Xenium, the Xenium data was cropped to the area included in Visium, because regions not included in Visium might have different morphology and cell type compositions that can change global spatial analysis results.

Cell type labels were transferred to the full TSU-21 Xenium data with SingleR (v2.11.3)^82^. The reference is the LUAD cells in the Human Lung Cell Atlas (HLCA) ^83^, which have pre-existing annotations (ann_finest_level). The H5AD file of the full HLCA was downloaded and read into R as a SingleCellExperiment (SCE) ^84^ object with zellkonverter (v1.19.2)^85^, which was subsetted to keep cells from LUAD samples. Because many cells were annotated “Unknown”, the SCE object was Leiden clustered with the batch corrected scANVI embeddings in order to annotate the “Unknown” cells. Differential expression (DE) was run with findMarkers() in scater (test.type = "wilcox", pval.type = "all"), and the marker genes were used to annotate Leiden clusters whose cells are predominantly “Unknown”. As other Leiden clusters map quite cleanly to pre-annotated cell types, the cell type labels (not “Unknown”) are transferred to the Leiden clusters. Then this SCE object, with cleaned cell type labels, was used as the reference in SingleR to transfer labels to Xenium. The transferred labels are used in Figure 2H to help interpret genes on the plot. The gene is deemed a marker of the cell type if in the DE results from the SCE object, the gene has the lowest p-value for that cell type, although it may have adjusted p < 0.05 for multiple cell types.

### Spatial binning

The transcript spots csv file from Xenium output was converted to parquet format with the arrow R package (v21.0.0.1), partitioned by gene, so they become much more efficient to be read repeatedly, one gene at a time. The whole transcript spots file was never read into memory. A square spatial grid was made with st_make_grid() in sf, with side lengths 8, 12, 16, 24, 32, 48, 64, 96, 128, 192, 256, and 384 μm. Larger side lengths were explored but not used because edge effect becomes too strong, making spatial analysis results too noisy. The aggregateTx() function in SpatialFeatureExperiment was used to aggregate the transcript spots, counting the number of spots intersecting each bin. Empty bins and bins outside the tissue boundary (found during Xenium QC with concave hull) are removed. The resulting gene count matrix and associated bin polygon geometries are saved as SFE objects to disk with alabaster.sfe.

To mitigate edge effect, the proportion of each bin occupied by cell polygons was computed, and bins with cell proportion lower than a threshold were removed. When log normalizing the data, The area of each bin occupied by cells was used as the size factor because bins with larger area not in cells are expected to have lower gene counts because of the area not in cells. The threshold is 0.9 for smallest bins, and around 0.2 for the largest bins. The threshold needs to be manually adjusted to avoid outliers in PCA caused by edge effects. ESDA was performed as in Xenium, for each sample and each bin size separately. The updated SFE objects after QC and with ESDA results are saved to disk with alabaster.sfe.

### Comparing Moran’s I and Lee’s L across scales

Moran’s I for each gene and Lee’s L from each gene pair were read into R from saved results for each bin size so they can be plotted as curves (Figure 2I). For Lee’s L, due to the large number of gene pairs, only pairs with absolute value of Lee’s L greater than 0.2 were kept for further analysis. These curves can be clustered to find patterns. In Figure 2I, Leiden clustering was performed on the difference between Moran’s I of successive scales for each gene, to cluster how Moran’s I changes through scales. Clustering performed on Moran’s I values themselves would differentiate between curves by Moran’s I values themselves rather than changes through scales. Lee’s L curves in Supplementary Figure 8 were clustered with hierarchical clustering with correlation (using 1 - cor(x) as distance) to cluster patterns of change through scales.

To demonstrate why Lee’s L changes through scale, the plots showing expression values of two genes with a bivariate palette in Figure 4 were made with the biscale R package (v1.0.0)^86^, with the “BlueGold” palette.

### Comparing Moran’s I and Lee’s L across stages

The above-mentioned curves were plotted for all samples and stages and considerable variability among different patients in the same stage has been observed for many genes and gene pairs. LMMs were used to determine whether the curves indeed differ among stages. The lme4 package (v1.1.37)^87^ was used to fit a LMM for each gene and gene pair. The Moran’s I or Lee’s L value is the response variable, and B-splines with degree 2 were used to model the curves. The full model has random effects for the intercept and slope for the B-spline term in the LUAD stage, i.e. each stage has its own intercept and slope. The ranova() function in lmerTest (v3.1.3)^88^ to test if the random effect terms are significant. The random slope but not the random intercept was dropped to see if the way Moran’s I or Lee’s L change through scales differ among stages, and both the random slope and random intercept are dropped to see if the curves differ among stages, even if the slopes do not differ significantly. Likelihood ratio test was used to compare the reduced and full models. The p-values were corrected for multiple hypothesis testing with the Benjamini-Hochberg method.

As most gene pairs have significant random effects, to narrow down to potentially biologically relevant gene pairs, we used CellChat’s (v2.1.2) ligand-receptor database^55^ to find gene pairs with significant slopes or random effects that are known to interact (Supplementary Figures 15-16).

### Comparing MULTISPATI gene loadings

Gene loadings are the eigenvectors from non-spatial and MULTISPATI PCA, which mean each gene’s contribution to each PC. The loading matrices from all bin sizes were read into R as part of the saved SFE objects. Since the eigenvectors can have flipped signs, all PC1’s are aligned to PC1 of the 8 μm data, and the same was done to other PC’s. To align the PC’s, for example, if PC1 at 12 μm is more similar to that at 8 μm with signs flipped, then the 12 μm PC1 will have its signs flipped. Then the loading of each gene is plotted as a curve to visualize the change in loadings through scales.

### Univariate synthetic data

To better understand Moran’s I curves through scales, we created synthetic data to see which synthetic patterns can recapitulate patterns in the curves. Although these patterns are very unrealistic, we have recapitulated some Moran’s I curve patterns. The following patterns were created on 8 μm bins from TSU-21 (Supplementary Figures 3, 4.4, 4.6, 4.7,also see processed data on Figshare):

● Binary, with selected regions set to 1 and the rest to 0

○ Large regions with straight, circular, and diamond shapes
○ Round spots, large (500-1000 μm in radius), medium large (200-500), medium small (100-250), small (50-150), and smaller (20-50)
○ Random lines with various thickness of the lines
○ Mixture of the large regions and the spots and lines
● Gradients

○ Large smooth polynomial gradients
○ Periodic patterns (large, medium, small)
○ Stripes (periodic in one direction)
○ Combination of large gradients and periodic patterns or stripes
○ The values in the gradient patterns are then scaled, centered, and exponentiated.
● No spatial pattern

○ Randomly sampling 0’s and 1’s
○ Sampling from a uniform distribution

Sparsity was introduced in the following ways: For binary patterns, a proportion of non-zero values are randomly set to 0. For all patterns, the values are used as means for Poisson distributions and noisier values are sampled from the Poisson distributions. To introduce sparsity, the means were divided by 2 or 5 before sampling. The sparsify score is the proportion set to 0, or when the Poisson mean is divided, e.g. when dividing by 5, the sparsify score is 1-⅕= 0.8.

After the patterns were created at the 8 μm resolution, they were aggregated into larger bins. To aggregate into 12 μm which would partially overlap with many 8 μm bins, the values at overlapping 8 μm bins were multiplied by the proportion of their overlap with the 12 μm bin. However, since the overlap creates artificially high Moran’s I that is not observed in real data, Moran’s I and Lee’s L values at 12 μm for synthetic are omitted when plotting and clustering. The 8 μm bins are nested in all of the larger bins from 16 to 384 μm, so the value at a larger bin would be the sum of values at all 8 μm bins covered by the larger bin. Then Moran’s I was computed for all patterns at all bin sizes.

### Multivariate synthetic data

To better understand Lee’s L and MULTISPATI curves through scales and to create more synthetic patterns for Moran’s I, we created multivariate synthetic data with correlations among genes. There are two cell types corresponding to two gene programs. Spatial patterns were created by different spatial distributions of two synthetic cell types. This synthetic dataset is based on TSU-20 at 8 μm resolution. The intention is not to make realistic simulations, but to recapitulate patterns in Lee’s L curves to better understand them.

Consensus NMF (cNMF) ^89^ was run on the real TSU-21 data, although we reimplemented and modified cNMF for much faster runtime as this was run for all samples and resolutions, although only TSU-21 was used in the context of making synthetic data. It was originally intended to find gene programs common and distinct to LUAD stages, and to find whether some gene programs at one stage appear in another stage at a different length scale. However, such an analysis with cNMF was not successful. The log normalized data was used for cNMF. First, the RcppML package (v0.5.6)^90^ was used to run cross validation to find the number of factors to use (k), holding out 5% of the data. Cross validation was run 10 times for k between 3 and 20, and the k with the smallest mean cross validation error was selected, which was 5 for TSU-21 at 8 μm resolution. Then 100 replicates were run with k=5. This is much faster than the original implementation of cNMF where 100 replicas are run for all values of k of interest in order to find k. As described in ^89^, 30 nearest neighbors were found for all factors from the 100 replicas, and outliers were removed. Here outliers were factors whose mean distance to their neighbors are greater than the 99th percentile. The remaining factors were k-means clustered with k=5. The centroids of the k-means clusters are the consensus NMF factors. Then non-negative least squares was used to find cell usage of each factor. Both the factors and usage were scaled to sum up to 1 as described in ^89^.

The 5 cell usage vectors were used to cluster the bins, to find bins that use each of the 5 gene programs. Two of the factors/gene programs roughly match PC2 and PC3 in non-spatial PCA. First, the gene with the highest loading in PC2 and PC3 (CD74 and FCMR respectively) were selected. Varying strengths of correlation were introduced. The strongest correlation was chosen by selecting the gene with the second highest loading (HLA-DPB1 and CD19 respectively in PC2 and PC3). Then among all genes whose loadings are the same sign as CD74 and FCMR, genes were selected at the 0.9, 0.5, 0.2 quantiles of their loadings, signifying different strengths of correlation. So each “gene program” has 5 genes, with varying strengths of correlation among them. The two corresponding cNMF factors were then subsetted to only include these genes to use as bases.

In NMF, the data matrix is decomposed into WH where W is the factors and H is the usage of each factor by each cell. The bases described above compose the W matrix, so there are 2 factors with 10 genes. For the H matrix, cell type labels are first assigned to each bin to make the spatial pattern. Each cell type corresponds to one factor, so entries in the usage vector corresponding to one cell type have usage values sampled from the fitted gamma distribution while entries corresponding to the other cell type are 0. The W matrix is multiplied by the H matrix to produce the simulated data matrix, which was then scaled so simulated mean expression of the 10 genes would match that in the real data. These values were used as means of Poisson distributions. Sparser versions were also made by dividing the means by 2 or 4. After the synthetic spatial count data were made, they were spatially aggregated in the same way as in univariate synthetic data. Moran’s I was run on all features and Lee’s L on all pairs of features, at all levels of aggregation.

The following spatial patterns were made (Supplementary Figure 11, also see processed data in Figshare):

● No spatial pattern – cell type labels were randomly assigned with varying proportions: 50/50, 70/30, and 90/10
● Large (50 to 150 μm in radius) and small spots (20 to 50 μm in radius) randomly distributed across the section; bins that intersect with the spots have one cell type, while the rest of the bins have the other cell type.
● Checkerboards with side lengths 8, 32, 96, and 128 μm to simulate negative spatial autocorrelation at different scales
● Checkerboard at 8 μm but one cell type is much more prevalent than the other. The checkerboard was made to assign two probability types bins; one type has probability 0.9 to select cell type A, while the other type has probability 0.5 to select cell type A, so cell type A is much more prevalent than cell type B. This makes a weaker negative spatial autocorrelation.
● Mixture of positive and negative spatial autocorrelation: two large spots patterns were made. In one of the patterns, bins that intersect with the spots get checkerboards. In the other spots pattern, bins intersecting the spots but not the checkerboard regions are cell type A, and the rest of the bins are cell type B.
● Mixture of negative spatial autocorrelation and no pattern: large random spots were made and bins that intersect the spots get checkerboards. The other bins get random cell type assignments.
● Mixture of positive spatial autocorrelation and no pattern: two sets of large random spots were made. Bins that intersect one set were assigned cell type A, and bins that intersect the other set were assigned cell type B. The rest get random assignments.

## Data availability

Visium and Xenium data Space Ranger outputs were downloaded from https://kero.hgc.jp/Ad-SpatialAnalysis_2024.html

Visium and Xenium data with QC and ESDA results from this study in ArtifactDB format, all spatial aggregations, and synthetic data are deposited on Figshare: https://doi.org/10.6084/m9.figshare.30876110

## Code availability

Code to reproduce analyses and figures: https://github.com/Computational-Morphogenomics-Group/scale_comparison

The wayfarer R package: https://github.com/Computational-Morphogenomics-Group/wayfarer

## Author Contributions

L.M. and B.D. conceived the study and developed the methodology. L.M. performed the investigation and data visualization, with input from A.H., L.C. and B.D. A.H. and L.C. contributed to biological insights when interpreting results. B.D. acquired funding and provided overall supervision and project administration. L.M. drafted the manuscript. L.M., B.D., A.H. and L.C. reviewed, edited and approved the final manuscript.

## Acknowledgements

We thank Jean Fan for discussions and suggestions regarding this project and Lior Pachter for discussing initial ideas regarding this project. BD acknowledges the support of the CIFAR MacMillan Multiscale Human Project.

## Notes

https://doi.org/10.6084/m9.figshare.30876110

### Competing Interest Statement

The authors have declared no competing interest.

## References

1. Han, M. et al. Programmable control of spatial transcriptome in live cells and neurons. Nature 643, 241–251 (2025).

2. Middleton, S. A., Eberwine, J. & Kim, J. Comprehensive catalog of dendritically localized mRNA isoforms from sub-cellular sequencing of single mouse neurons. BMC Biol. 17, 5 (2019).

3. Zhao, J., Andreev, I. & Silva, H. M. Resident tissue macrophages: Key coordinators of tissue homeostasis beyond immunity. Sci. Immunol. 9, eadd1967 (2024).

4. Kietzmann, T. Metabolic zonation of the liver: The oxygen gradient revisited. Redox Biol. 11, 622–630 (2017).

5. Hawrylycz, M. J. et al. An anatomically comprehensive atlas of the adult human brain transcriptome. Nature 489, 391–399 (2012).

6. Jia, E. et al. Spatial transcriptome profiling of mouse hippocampal single cell microzone in Parkinson’s disease. Int. J. Mol. Sci. 24, 1810 (2023).

7. Oliveira, M. F., et al. Characterization of immune cell populations in the tumor microenvironment of colorectal cancer using high definition spatial profiling. bioRxiv (2024) doi:10.1101/2024.06.04.597233.

8. Chen, A. et al. Spatiotemporal transcriptomic atlas of mouse organogenesis using DNA nanoball-patterned arrays. Cell 185, 1777–1792.e21 (2022).

9. Nucera, M. R., et al. Application of spatial transcriptomics across organoids: a high-resolution spatial whole-transcriptome benchmarking dataset. bioRxiv (2025) doi:10.1101/2025.05.04.651803.

10. Li, L. et al. An organ-wide spatiotemporal transcriptomic and cellular atlas of the regenerating zebrafish heart. Nat. Commun. 16, 3716 (2025).

11. Cervilla, S., et al. Benchmarking of spatial transcriptomics platforms across six cancer types. bioRxiv (2024) doi:10.1101/2024.05.21.593407.

12. Ozirmak Lermi, N., et al. Comparison of imaging-based single-cell resolution spatial transcriptomics profiling platforms using formalin-fixed, paraffin-embedded tumor samples. bioRxiv (2024) doi:10.1101/2024.12.13.628390.

13. You, Y. et al. Systematic comparison of sequencing-based spatial transcriptomic methods. Nat. Methods 21, 1743–1754 (2024).

14. Zemp, F. J. et al. Development and first-in-human CAR T therapy against the pathognomonic MiT fusion driven protein GPNMB. medRxiv (2025) doi:10.1101/2025.02.26.24319604.

15. Liu, C. et al. Integrative single-cell and spatial transcriptome analysis reveals heterogeneity of human liver progenitor cells. Hepatol. Commun. 9, (2025).

16. Li, Z. et al. Presence of onco-fetal neighborhoods in hepatocellular carcinoma is associated with relapse and response to immunotherapy. *Nat*. Cancer 5, 167–186 (2024).

17. Golfinos-Owens, A. E., et al. Integrated single-cell and spatial analysis identifies context-dependent myeloid-T cell interactions in head and neck cancer immune checkpoint blockade response. bioRxivorg (2025) doi:10.1101/2025.03.24.644582.

18. Grande, E. et al. Spatial biomarkers of response to neoadjuvant therapy in muscle-invasive bladder cancer: the DUTRENEO trial. medRxiv (2025) doi:10.1101/2025.02.07.25321742.

19. Wang, H. et al. Type I interferon drives T cell cytotoxicity by upregulation of interferon regulatory factor 7 in autoimmune kidney diseases in mice. Nat. Commun. 16, 4686 (2025).

20. Zhang, M. et al. Spatially resolved cell atlas of the mouse primary motor cortex by MERFISH. Nature 598, 137–143 (2021).

21. Joulia, R. et al. A single-cell spatial chart of the airway wall reveals proinflammatory cellular ecosystems and their interactions in health and asthma. Nat. Immunol. 26, 920–933 (2025).

22. Zeng, H. What is a cell type and how to define it? Cell 185, 2739–2755 (2022).

23. Cliff, A. D. & Ord, J. K. Spatial Processes: Models & Applications. (Pion, London, England, 1981).

24. Benninghoff, W. S. Quantitative Plant Ecology . P. Grieg-Smith. Academic Press, New York; Butterworths, London, 1957. x + 198 pp. Illus. $6. Science 128, 1620–1621 (1958).

25. Ludwig, J. A. A test of different quadrat variance methods for the analysis of spatial pattern. in Spatial and temporal analysis in ecology (eds. Cormack, R. M. & Ord, J. K.) 289–304 (Fairland, Md. : International Co-operative Pub. House, 1979).

26. Griffith, D. A., Chun, Y. & Li, B. *Spatial Regression Analysis Using Eigenvector Spatial Filtering*. (Academic Press, San Diego, CA, 2019). doi:10.1016/c2017-0-01015-7.

27. Baddeley, A., Rubak, E. & Turner, R. Spatial Point Patterns. (Apple Academic Press, Oakville, MO, 2015).

28. Jombart, T., Dray, S. & Dufour, A.-B. Finding essential scales of spatial variation in ecological data: a multivariate approach. Ecography (Cop.) 32, 161–168 (2009).

29. Manley, D. Scale, aggregation, and the modifiable areal unit problem. in *Handbook of Regional Science* 1711–1725 (Springer Berlin Heidelberg, Berlin, Heidelberg, 2021). doi:10.1007/978-3-662-60723-7_69.

30. Tanevski, J., Flores, R. O. R., Gabor, A., Schapiro, D. & Saez-Rodriguez, J. Explainable multiview framework for dissecting spatial relationships from highly multiplexed data. Genome Biol. 23, 97 (2022).

31. Palla, G. et al. Squidpy: a scalable framework for spatial omics analysis. Nat. Methods 19, 171–178 (2022).

32. Dos Santos Peixoto, R., et al. Characterizing cell-type spatial relationships across length scales in spatially resolved omics data. bioRxivorg (2024) doi:10.1101/2023.10.05.560733.

33. Ameen, F., Robertson, N., Lin, D. M., Ghazanfar, S. & Patrick, E. Kontextual: Reframing analysis of spatial omics data reveals consistent cell relationships across images. bioRxiv (2024) doi:10.1101/2024.09.03.611109.

34. Jerison, E. R., Romeo, N. & Quake, S. R. Spatially-structured inflammatory response in the presence of a uniform stimulus. bioRxiv (2025) doi:10.1101/2025.01.28.635318.

35. Venkat, A. et al. Mapping the gene space at single-cell resolution with gene signal pattern analysis. Nat. Comput. Sci. 4, 955–977 (2024).

36. Maher, K. & Wang, X. Harmonic representations of regions and interactions in spatial transcriptomics. bioRxiv (2024) doi:10.1101/2024.08.14.607982.

37. Svensson, V., Teichmann, S. A. & Stegle, O. SpatialDE: identification of spatially variable genes. Nat. Methods 15, 343–346 (2018).

38. Sun, S., Zhu, J. & Zhou, X. Statistical analysis of spatial expression patterns for spatially resolved transcriptomic studies. Nat. Methods 17, 193–200 (2020).

39. Weber, L. M., Saha, A., Datta, A., Hansen, K. D. & Hicks, S. C. nnSVG for the scalable identification of spatially variable genes using nearest-neighbor Gaussian processes. Nat. Commun. 14, 4059 (2023).

40. Aihara, G., et al. SEraster: a rasterization preprocessing framework for scalable spatial omics data analysis. Preprint at (2024).

41. Haga, Y. et al. Whole-genome sequencing reveals the molecular implications of the stepwise progression of lung adenocarcinoma. Nat. Commun. 14, 8375 (2023).

42. Takano, Y. et al. Spatially resolved gene expression profiling of tumor microenvironment reveals key steps of lung adenocarcinoma development. Nat. Commun. 15, 10637 (2024).

43. Moran, P. A. P. The interpretation of statistical maps. J. R. Stat. Soc. Series B Stat. Methodol. 10, 243–251 (1948).

44. Li, Z., et al. Benchmarking computational methods to identify spatially variable genes and peaks. bioRxivorg (2023) doi:10.1101/2023.12.02.569717.

45. Chen, X. et al. Benchmarking algorithms for spatially variable gene identification in spatial transcriptomics. Bioinformatics 41, (2025).

46. Lee, S.-I. Developing a bivariate spatial association measure: An integration of Pearson’s r and Moran’s I. J. Geogr. Syst. 3, 369–385 (2001).

47. Li, K.-J. et al. NAT10 promotes prostate cancer growth and metastasis by acetylating mRNAs of HMGA1 and KRT8. Adv. Sci. (Weinh.) 11, e2310131 (2024).

48. Zhao, J. & Lan, G. TFAP2A activates HMGA1 to promote glycolysis and lung adenocarcinoma progression. Pathol. Res. Pract. 249, 154759 (2023).

49. Aifantis, I. et al. A critical role for the cytoplasmic tail of pTalpha in T lymphocyte development. Nat. Immunol. 3, 483–488 (2002).

50. Materna, M. et al. The immunopathological landscape of human pre-TCRα deficiency: From rare to common variants. Science 383, eadh4059 (2024).

51. Dray, S., Saïd, S. & Débias, F. Spatial ordination of vegetation data using a generalization of Wartenberg’s multivariate spatial correlation. J. Veg. Sci. 19, 45–56 (2008).

52. Wartenberg, D. Multivariate spatial correlation: A method for exploratory geographical analysis. Geogr. Anal. 17, 263–283 (1985).

53. Yin, L., Zhang, H., Shang, Y., Wu, S. & Jin, T. ErbB/HER family in cancer immunology: therapeutic advances and mechanisms. Drug Discov. Today 30, 104436 (2025).

54. Ding, Q. et al. CXCL9: evidence and contradictions for its role in tumor progression. Cancer Med. 5, 3246–3259 (2016).

55. Jin, S. et al. Inference and analysis of cell-cell communication using CellChat. Nat. Commun. 12, 1088 (2021).

56. Zhang, L. et al. PRG4 represses the genesis and metastasis of osteosarcoma by inhibiting PDL1 expression. Tissue Cell 88, 102409 (2024).

57. Dituri, F. et al. Proteoglycan-4 is correlated with longer survival in HCC patients and enhances sorafenib and regorafenib effectiveness via CD44 in vitro. Cell Death Dis. 11, 984 (2020).

58. Ge, Q. et al. Spatially segregated APOE+ macrophages restrict immunotherapy efficacy in clear cell renal cell carcinoma. Theranostics 15, 5312–5336 (2025).

59. Khan, S. M., et al. Mapping the spatial architecture of glioblastoma from core to edge delineates niche-specific tumor cell states and intercellular interactions. bioRxivorg (2025) doi:10.1101/2025.04.04.647096.

60. Pei, G. et al. Spatial mapping of transcriptomic plasticity in metastatic pancreatic cancer. Nature 642, 212–221 (2025).

61. Jelinski, D. E. & Wu, J. The modifiable areal unit problem and implications for landscape ecology. Landsc. Ecol. 11, 129–140 (1996).

62. Harris, P., Brunsdon, C. & Charlton, M. Geographically weighted principal components analysis. *Geogr*. Inf. Syst. 25, 1717–1736 (2011).

63. Elsayed, G. F. & Cunningham, J. P. Structure in neural population recordings: an expected byproduct of simpler phenomena? Nat. Neurosci. 20, 1310–1318 (2017).

64. Barber, R. F. & Candès, E. J. A knockoff filter for high-dimensional selective inference. Ann. Stat. 47, 2504–2537 (2019).

65. Beker, O. et al. Patches: A representation learning framework for decoding shared and condition-specific transcriptional programs in wound healing. bioRxiv (2024) doi:10.1101/2024.12.23.630186.

66. Wickham, H. *Ggplot2: Elegant Graphics for Data Analysis*. (Springer, New York, NY, 2009).

67. Pedersen, T. L. patchwork: The Composer of Plots. CRAN: Contributed Packages The R Foundation 10.32614/cran.package.patchwork (2019).

68. Moses, L., Huseynov, A., Rich, J. M. & Pachter, L. Geospatially informed representation of spatial genomics data with SpatialFeatureExperiment. bioRxivorg (2025) doi:10.1101/2025.02.24.640007.

69. Totty, M., Hicks, S. C. & Guo, B. SpotSweeper: spatially aware quality control for spatial transcriptomics. Nat. Methods 22, 1520–1530 (2025).

70. Lun, A. T. L., McCarthy, D. J. & Marioni, J. C. A step-by-step workflow for low-level analysis of single-cell RNA-seq data with Bioconductor. F1000Res. 5, 2122 (2016).

71. McCarthy, D. J., Campbell, K. R., Lun, A. T. L. & Wills, Q. F. Scater: pre-processing, quality control, normalization and visualization of single-cell RNA-seq data in R. Bioinformatics 33, 1179–1186 (2017).

72. Moses, L., et al. Voyager: exploratory single-cell genomics data analysis with geospatial statistics. *bioRxivorg* (2023) doi:10.1101/2023.07.20.549945.

73. A Case Study in Fitting Area-Proportional Euler Diagrams with Ellipses Using Eulerr.

74. Lun, A. *BiocNeighbors*. (Bioconductor, 2018). doi:10.18129/B9.BIOC.BIOCNEIGHBORS.

75. Pebesma, E. & Bivand, R. Spatial Data Science: With Applications in R. (CRC Press, Boca Raton, FL, 2023).

76. Pebesma, E. Simple features for R: Standardized support for spatial vector data. R J. 10, 439 (2018).

77. Atta, L., Clifton, K., Anant, M., Aihara, G. & Fan, J. Gene count normalization in single-cell imaging-based spatially resolved transcriptomics. Genome Biol. 25, 153 (2024).

78. Lun, A. Bluster. (Bioconductor, 2020). doi:10.18129/B9.BIOC.BLUSTER.

79. Pau, G., Fuchs, F., Sklyar, O., Boutros, M. & Huber, W. EBImage--an R package for image processing with applications to cellular phenotypes. Bioinformatics 26, 979–981 (2010).

80. Hijmans, R. J. terra: Spatial Data Analysis. CRAN: Contributed Packages The R Foundation 10.32614/cran.package.terra (2020).

81. Heriche, J.-K. LOMAR: Localization microscopy data analysis. CRAN: Contributed Packages The R Foundation 10.32614/cran.package.lomar (2021).

82. Aran, D. et al. Reference-based analysis of lung single-cell sequencing reveals a transitional profibrotic macrophage. Nat. Immunol. 20, 163–172 (2019).

83. Sikkema, L. et al. An integrated cell atlas of the lung in health and disease. Nat. Med. 29, 1563–1577 (2023).

84. Amezquita, R. A. et al. Orchestrating single-cell analysis with Bioconductor. Nat. Methods 17, 137–145 (2020).

85. Luke Zappia, A. L. *Zellkonverter*. (Bioconductor, 2020). doi:10.18129/B9.BIOC.ZELLKONVERTER.

86. Prener, C., Grossenbacher, T. & Zehr, A. biscale: Tools and Palettes for Bivariate Thematic Mapping. CRAN: Contributed Packages The R Foundation 10.32614/cran.package.biscale (2019).

87. Bates, D., Mächler, M., Bolker, B. & Walker, S. Fitting linear mixed-effects models Usinglme4. J. Stat. Softw. 67, (2015).

88. Kuznetsova, A., Brockhoff, P. B. & Christensen, R. H. B. LmerTest package: Tests in linear mixed effects models. J. Stat. Softw. 82, (2017).

89. Kotliar, D. et al. Identifying gene expression programs of cell-type identity and cellular activity with single-cell RNA-Seq. Elife 8, (2019).

90. DeBruine, Z. J., Pospisilik, J. A. & Triche, T. J., Jr. Fast and interpretable non-negative matrix factorization for atlas-scale single cell data. bioRxiv (2021) doi:10.1101/2021.09.01.458620.

