## Supplementary figures for "Wayfarer: A multiscale framework for spatial analysis of tumor progression"

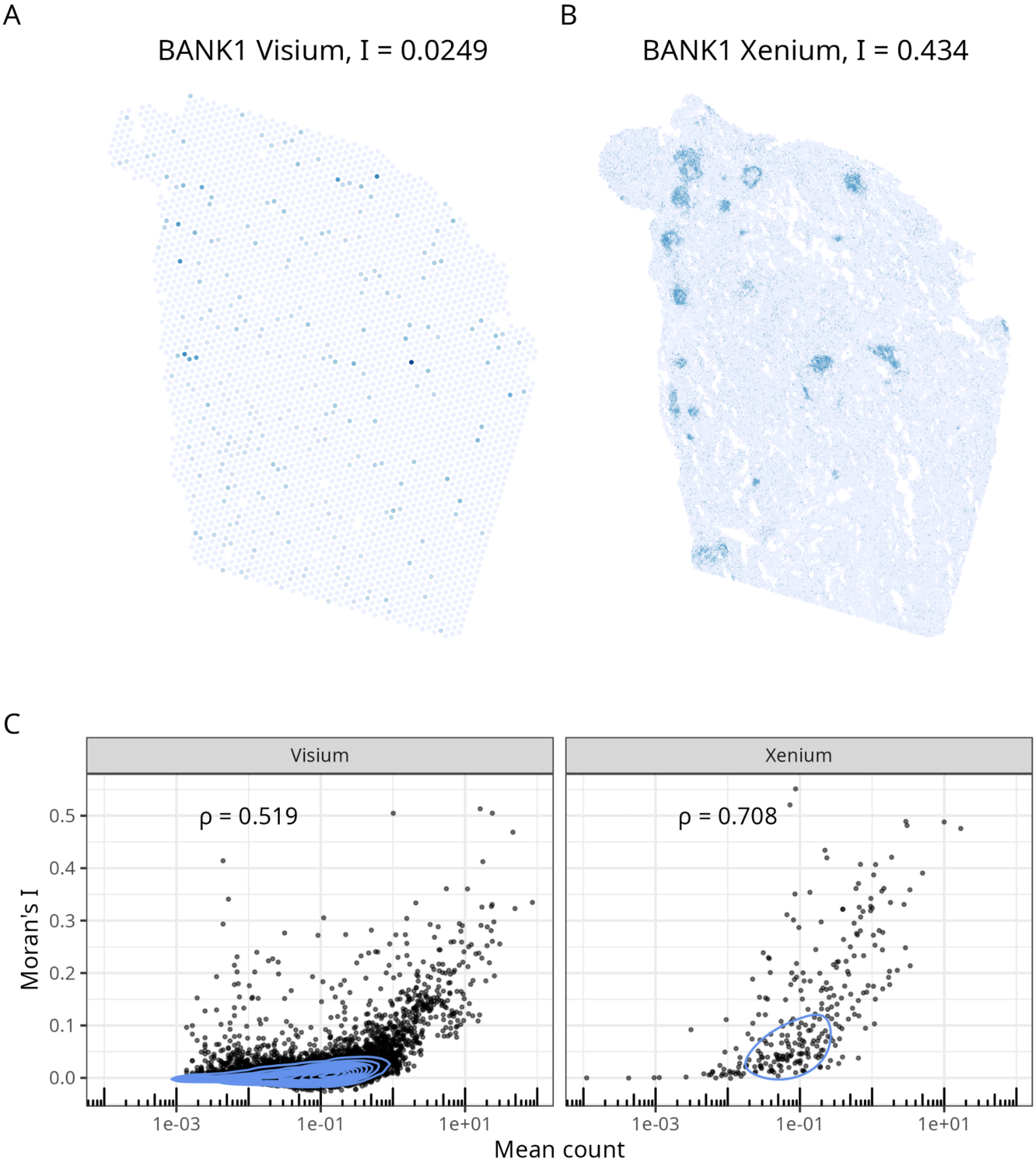


Supplementary Figure 1: A) Log normalized expression of BANK1 in real Visium in TSU-21, with Moran's I = 0.0249. B) Log normalized expression of BANK1 in Xenium plotted on single cells, with Moran's I = 0.434. C) Scatter plot of Moran's I of each gene vs. its mean counts in each Visium spot or each cell in Xenium. Spearman correlation is 0.519 for Visium and 0.708 for Xenium. The light blue curves are point density contours.

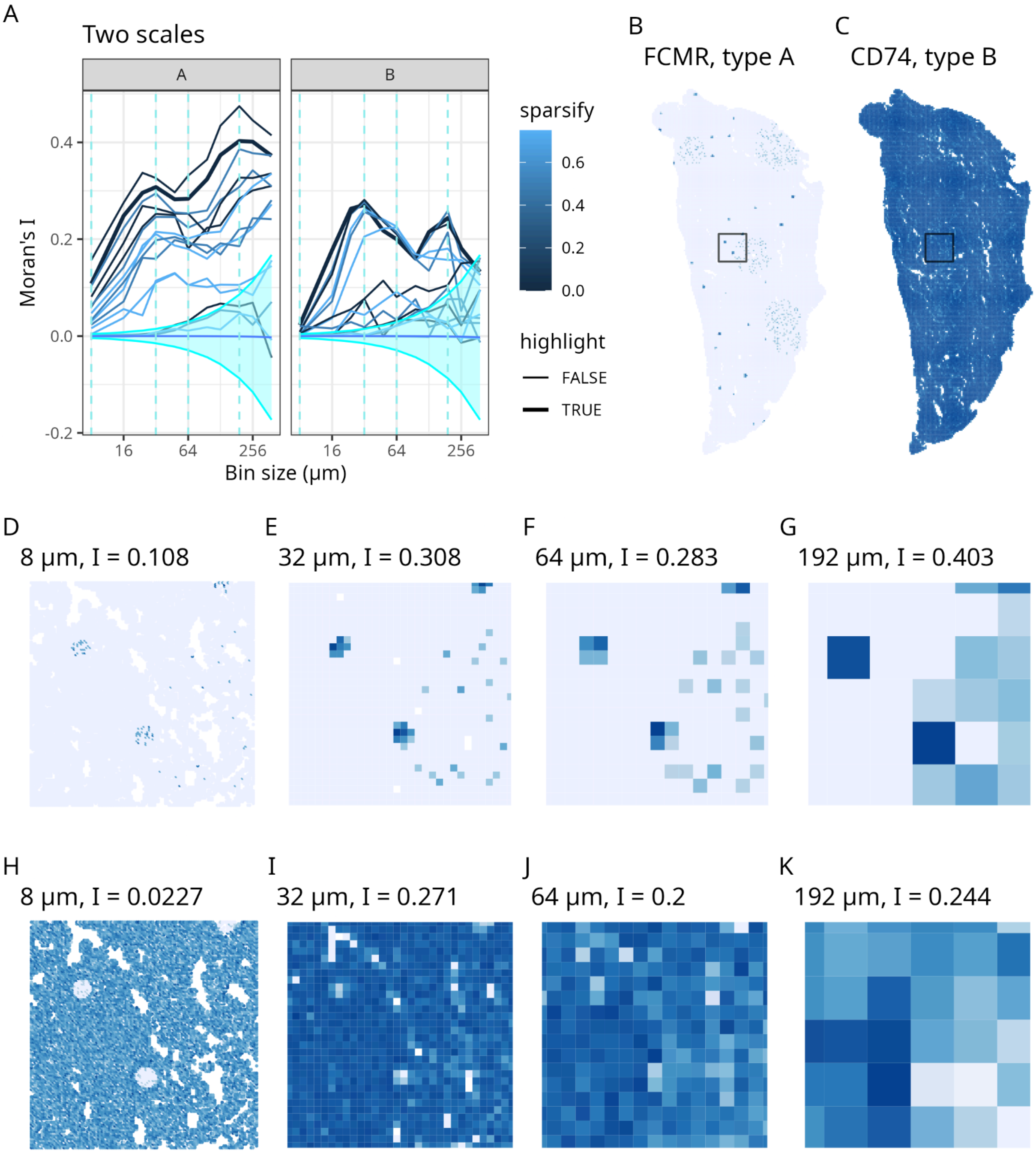


Supplementary Figure 2: Synthetic pattern with spatial structures at two different length scales. A) Moran's I vs. bin size for each synthetic feature. The curves are colored by how much the pattern was sparsified in construction, where a higher sparsify value means sparser. The two features plotted in other panels are shown in bold curves. Types A and B are two different gene programs (Methods). B) Example of Type A pattern, showing small and large spots. C) Example of Type B pattern, which is here the complement of Type A. D-G) Zooming into the box in B, at bin sizes of 8, 32, 64, and 192 µm. H-K) Zooming into the box in C.

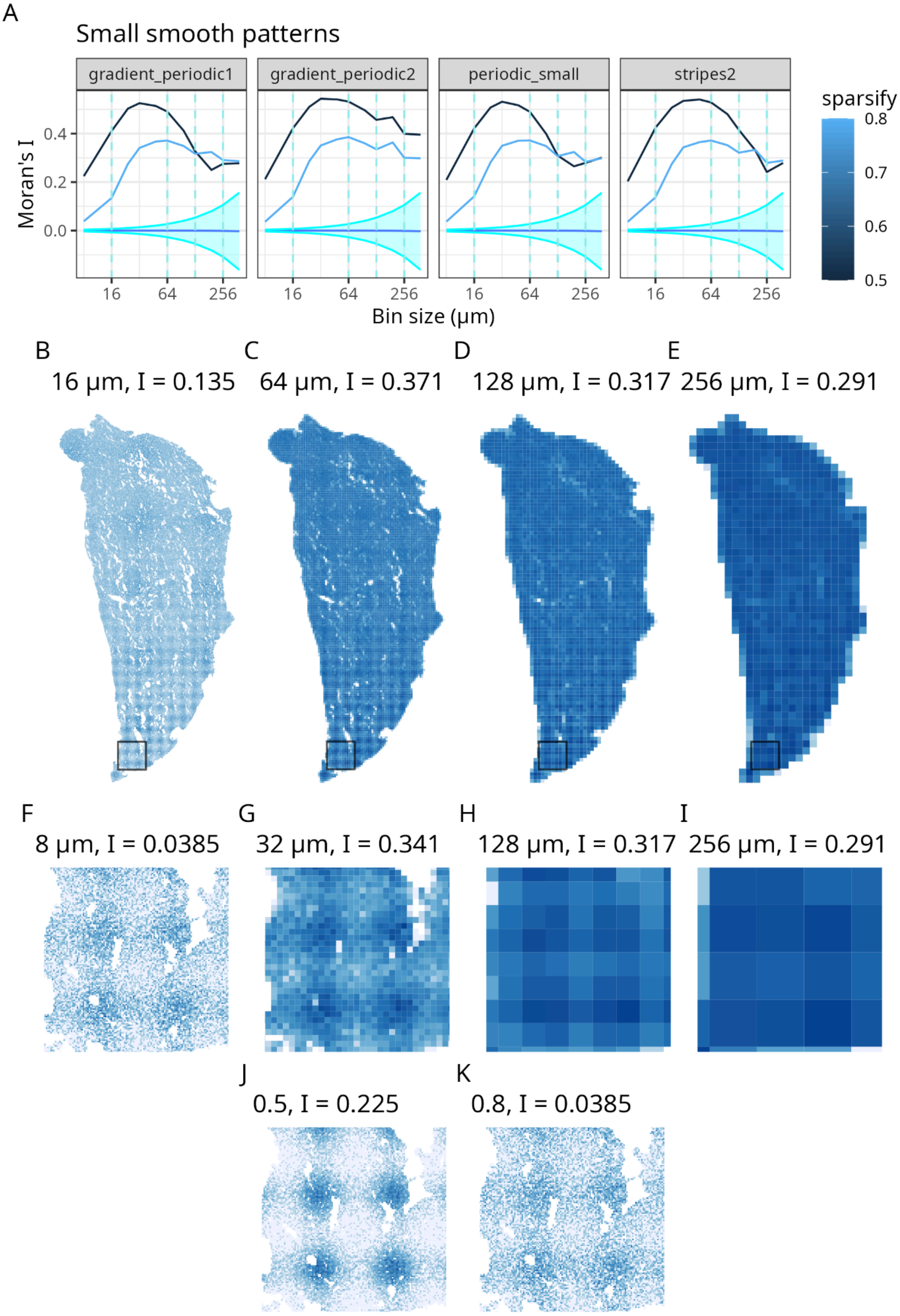
A
Small smooth patterns
gradient_periodic1
gradient_periodic2
periodic_small
stripes2
Moran's I
Bin size (μm)
sparsify
B
16 μm, I = 0.135
C
64 μm, I = 0.371
D
128 μm, I = 0.317
E
256 μm, I = 0.291
F
8 μm, I = 0.0385
G
32 μm, I = 0.341
H
128 μm, I = 0.317
I
256 μm, I = 0.291
J
0.5, I = 0.225
K
0.8, I = 0.0385

Supplementary Figure 3: Synthetic periodic pattern with shorter period. A) Moran's I of various synthetic patterns with short period, colored with sparsity. B-E) The specific pattern plotted here is the sparser version of gradient_periodic1. The pattern on the entire section is plotted at bin sizes 16, 64, 128, and 256 μm. F-I) Zooming into the box shown in B-E to show the local pattern. J) Same as F, but a less sparse version. K) Same as F, but showing the degree of sparsity (see Methods for details).

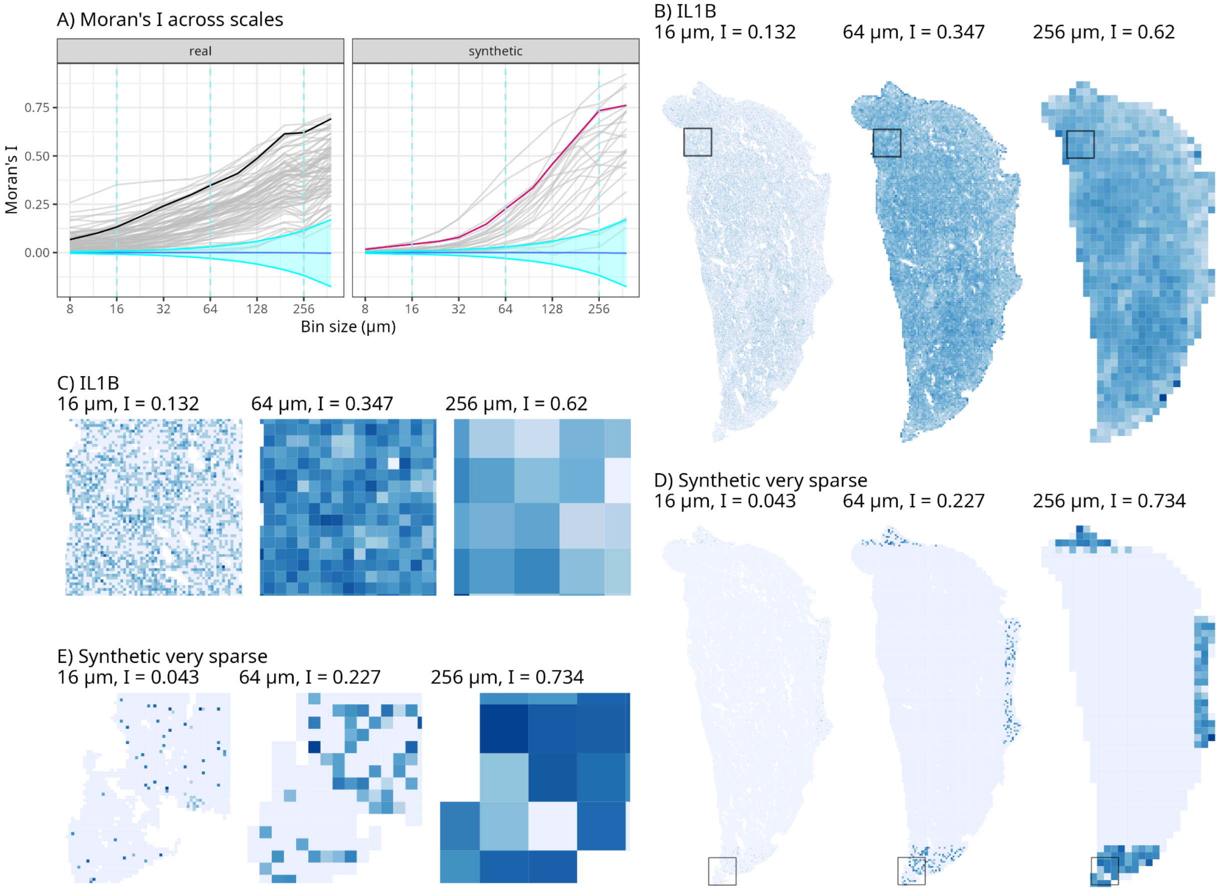


Supplementary Figure 4.1: Examples from cluster 1 of Moran's I curves in Figure 2I. A) Moran's I vs. bin size for each gene; curves from real data and synthetic data are plotted in separate facets. The curves not in gray are from the features plotted spatially in panels B-E. B) Global pattern of a gene from real data, plotted at different bin sizes. C) Zooming into the box shown in B. D) Global pattern of a synthetic feature. E) Zooming into the box shown in D.

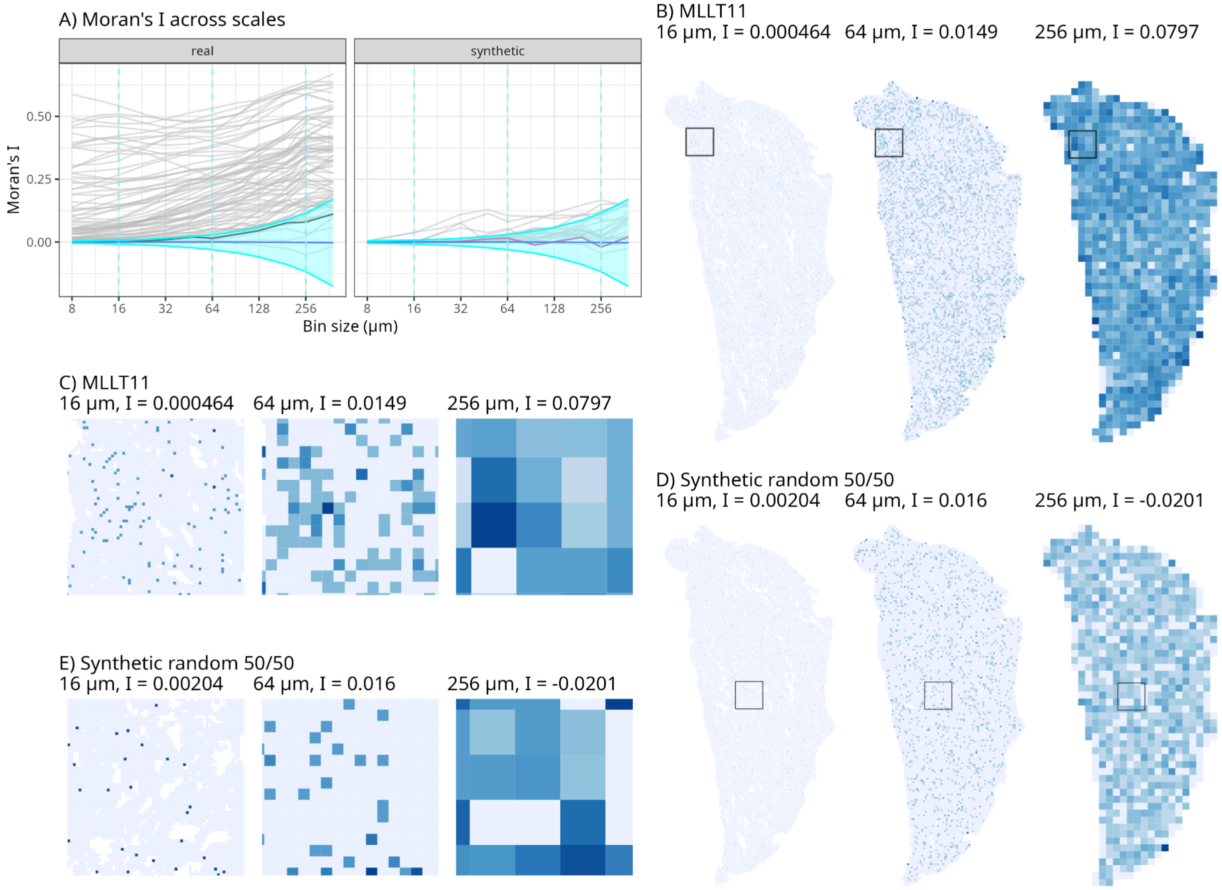


Supplementary Figure 4.2: Same as Supplementary Figure 4.1, but for cluster 3 in Figure 2I

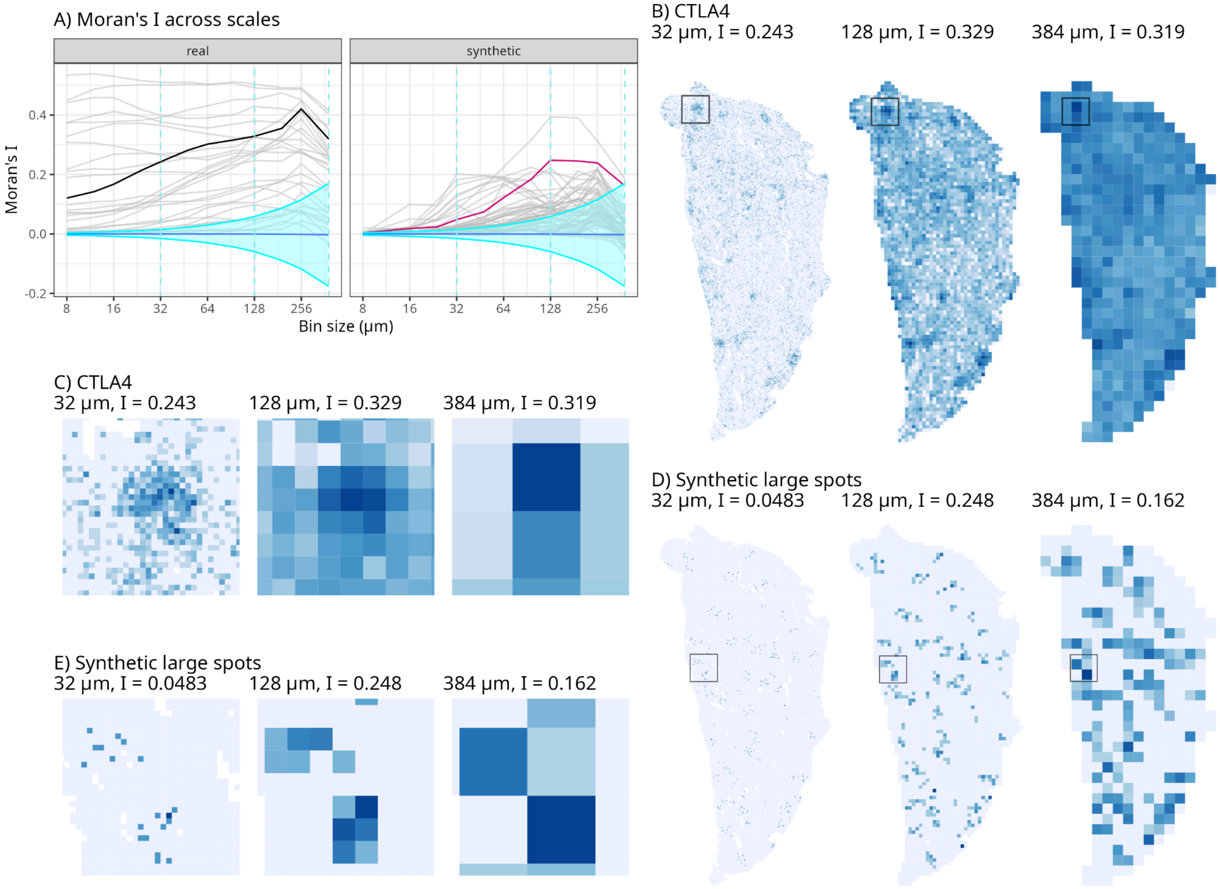


Supplementary Figure 4.3: Same as Supplementary Figure 4.1, but for cluster 4 in Figure 2I

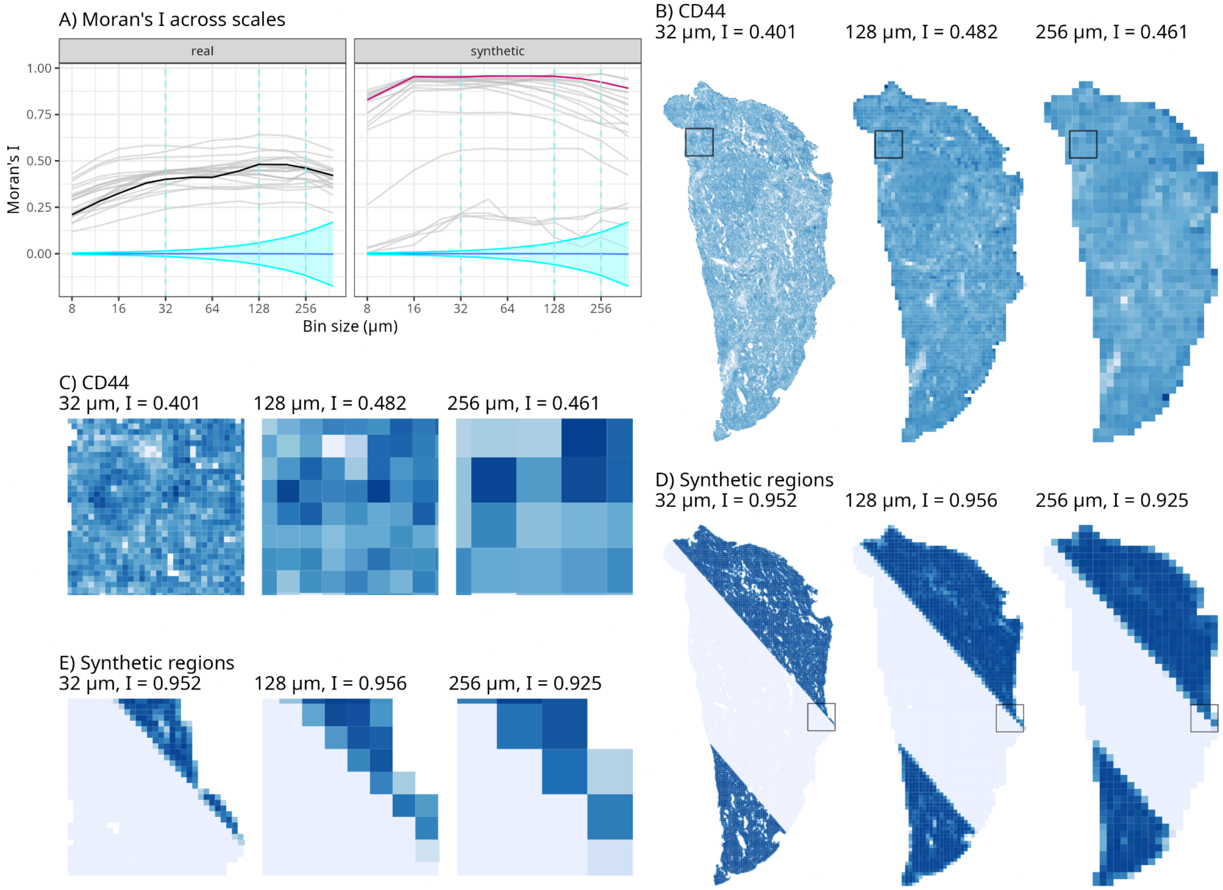


Supplementary Figure 4.4: Same as Supplementary Figure 4.1, but for cluster 5 in Figure 2I

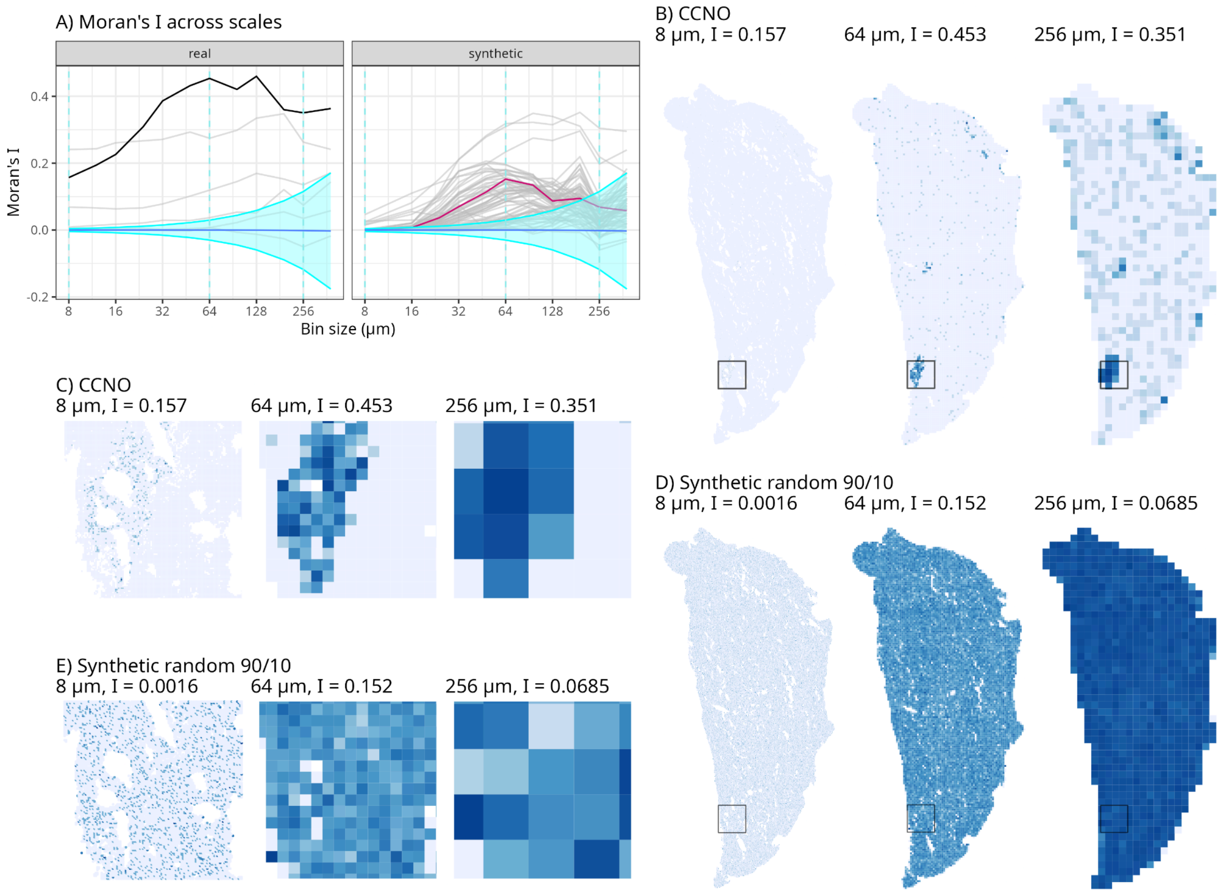


Supplementary Figure 4.5: Same as Supplementary Figure 4.1, but for cluster 6 in Figure 2I

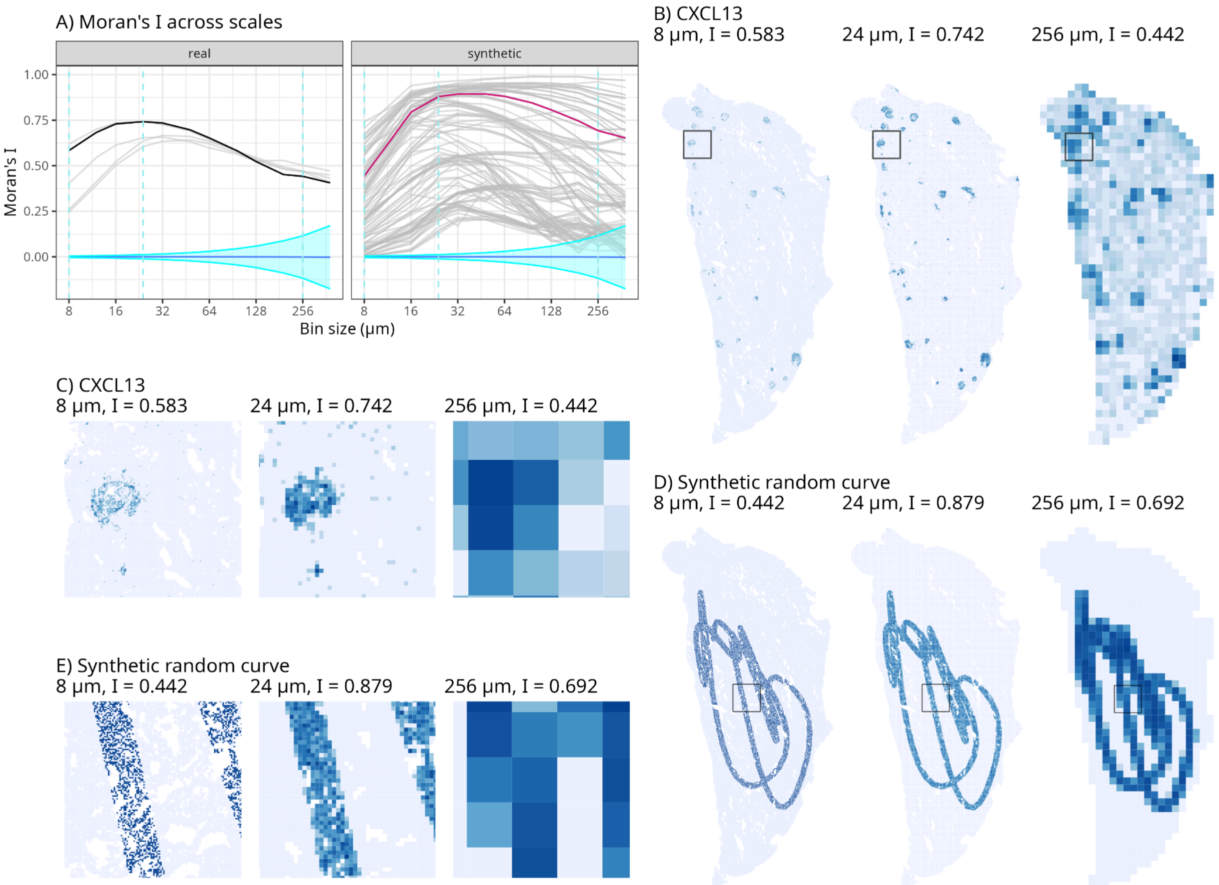


Supplementary Figure 4.6: Same as Supplementary Figure 4.1, but for cluster 7 in Figure 2I

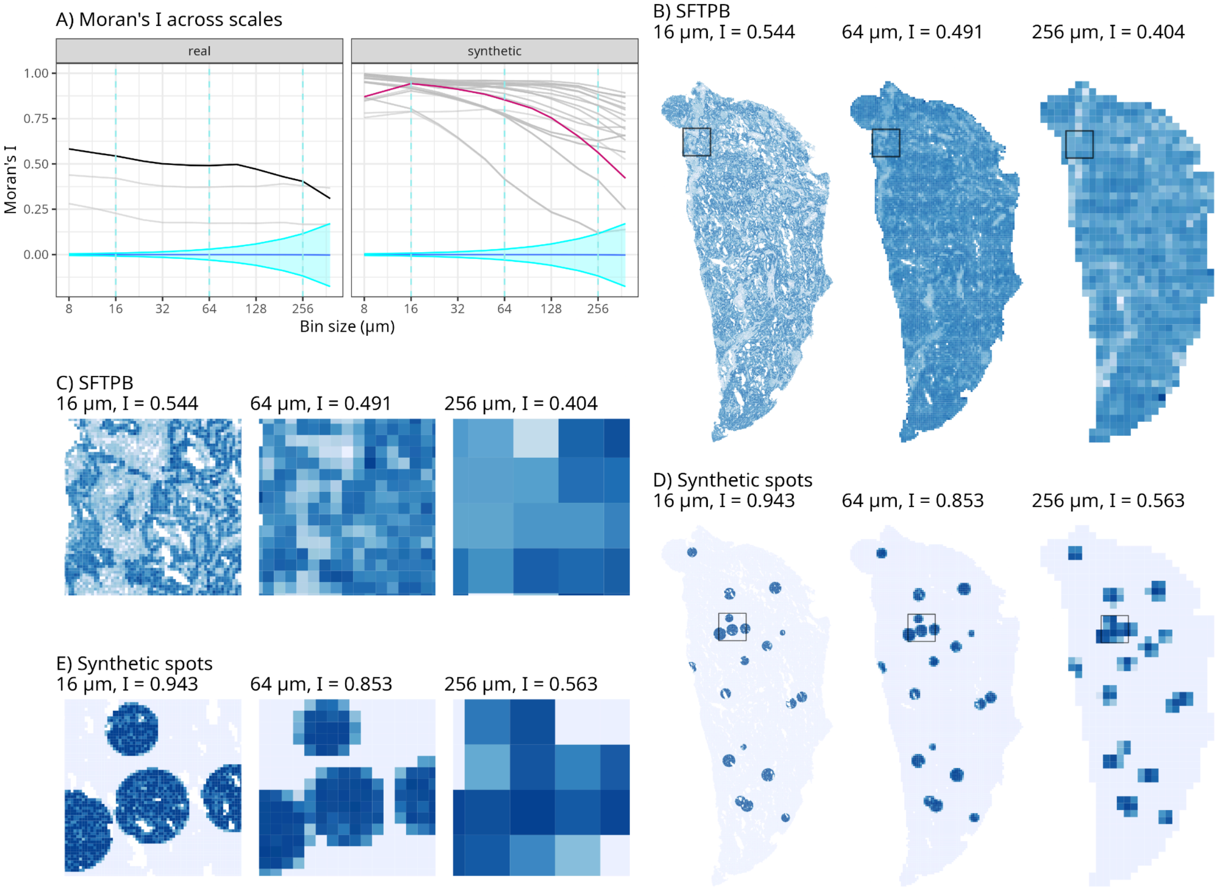


Supplementary Figure 4.7: Same as Supplementary Figure 4.1, but for cluster 8 in Figure 2I

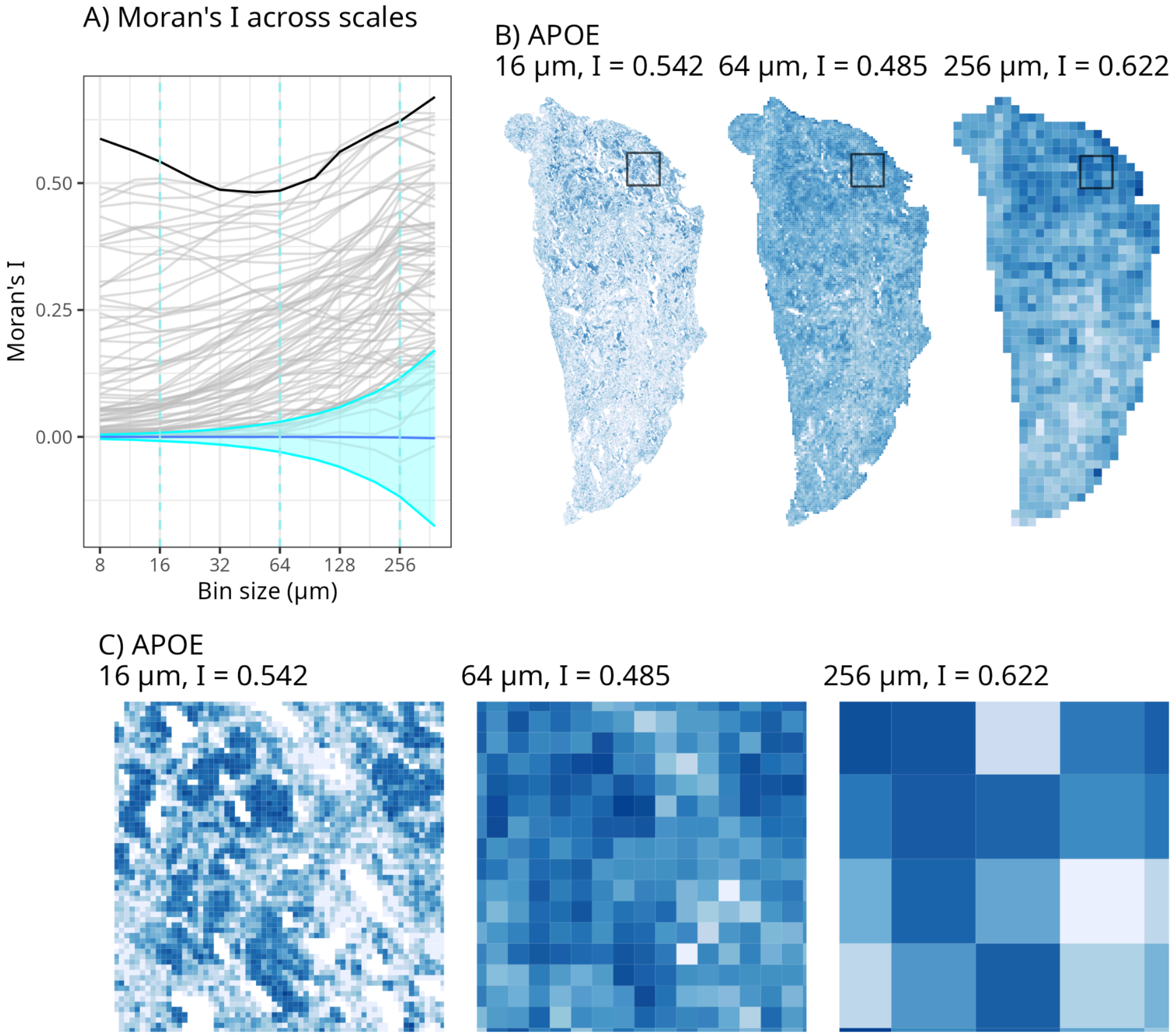


Supplementary Figure 5: Example of bimodal Moran's I curve. A) Moran's I curves from cluster 3 in Figure 2I, with the curve for APOE highlighted. B) APOE expression in the whole section of TSU-21 at bin sizes of 16, 64, and 256 μm. C) Zooming into the box marked in B to show local patterns.

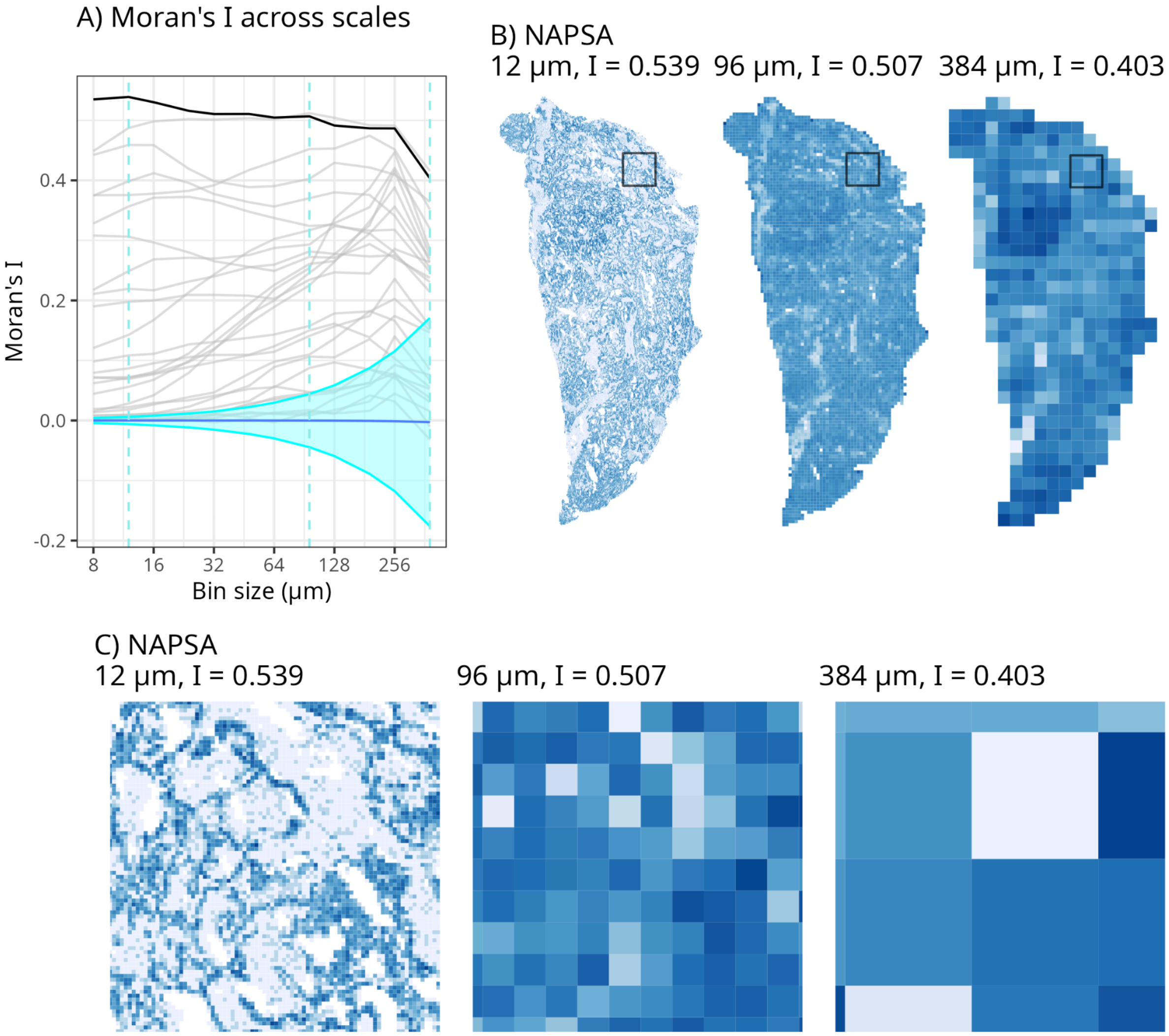


Supplementary Figure 6: Example of a somewhat flat Moran's I curve. A) Moran's I curves from cluster 4 in Figure 2I, with the curve for NAPSA highlighted. B) NAPSA expression in the whole section of TSU-21 at bin sizes of 12, 96, and 384 μm. C) Zooming into the box marked in B to show local patterns.

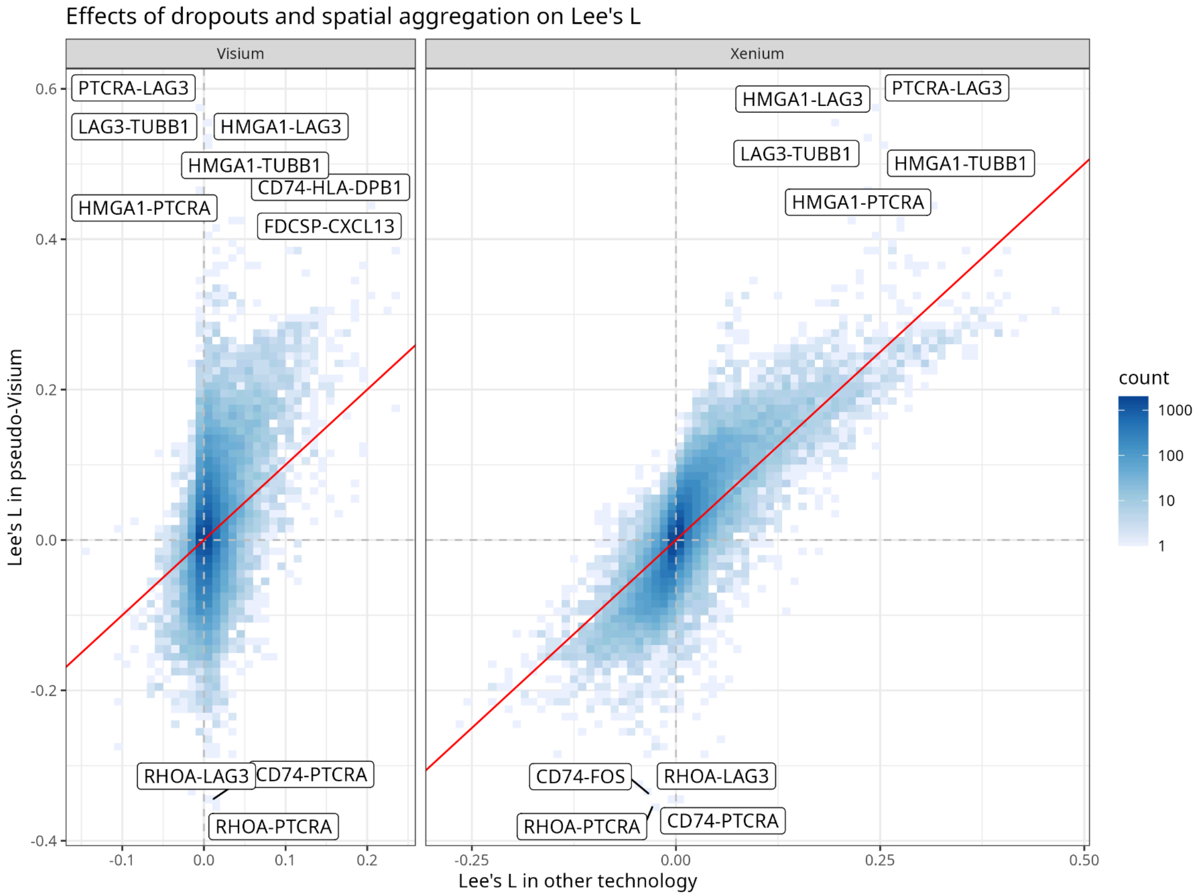


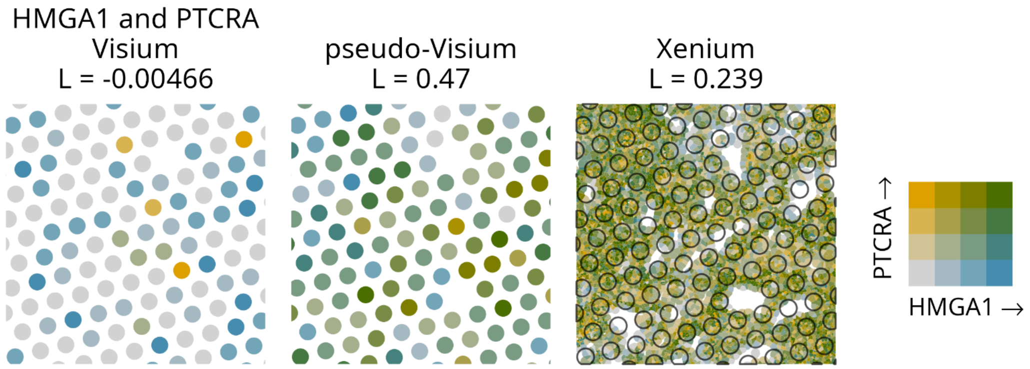


Supplementary Figure 7: Comparison of Lee's L for gene pairs in Visium, Xenium, and pseudo-Visium. Values for pseudo-Visium are plotted on the y-axis, while those for Visium and Xenium are on the x-axis. Due to the large number of gene pairs, a 2D histogram is shown, and darker color means more points (i.e. gene pairs). The red line is y = x, and gene pairs the furthest from the line are labeled. Dotted lines mark y = 0 and x = 0. HMGA1 and PTCRA deviate far from y = x so they are plotted spatially as an example. In the bivariate palette, yellow means high in PTCRA and low in HMGA1, blue means high in HMGA1 and low in PTCRA, green means high in both, and gray means low in both. Outlines of the aligned Visium spots are superimposed on the Xenium single cell plot.

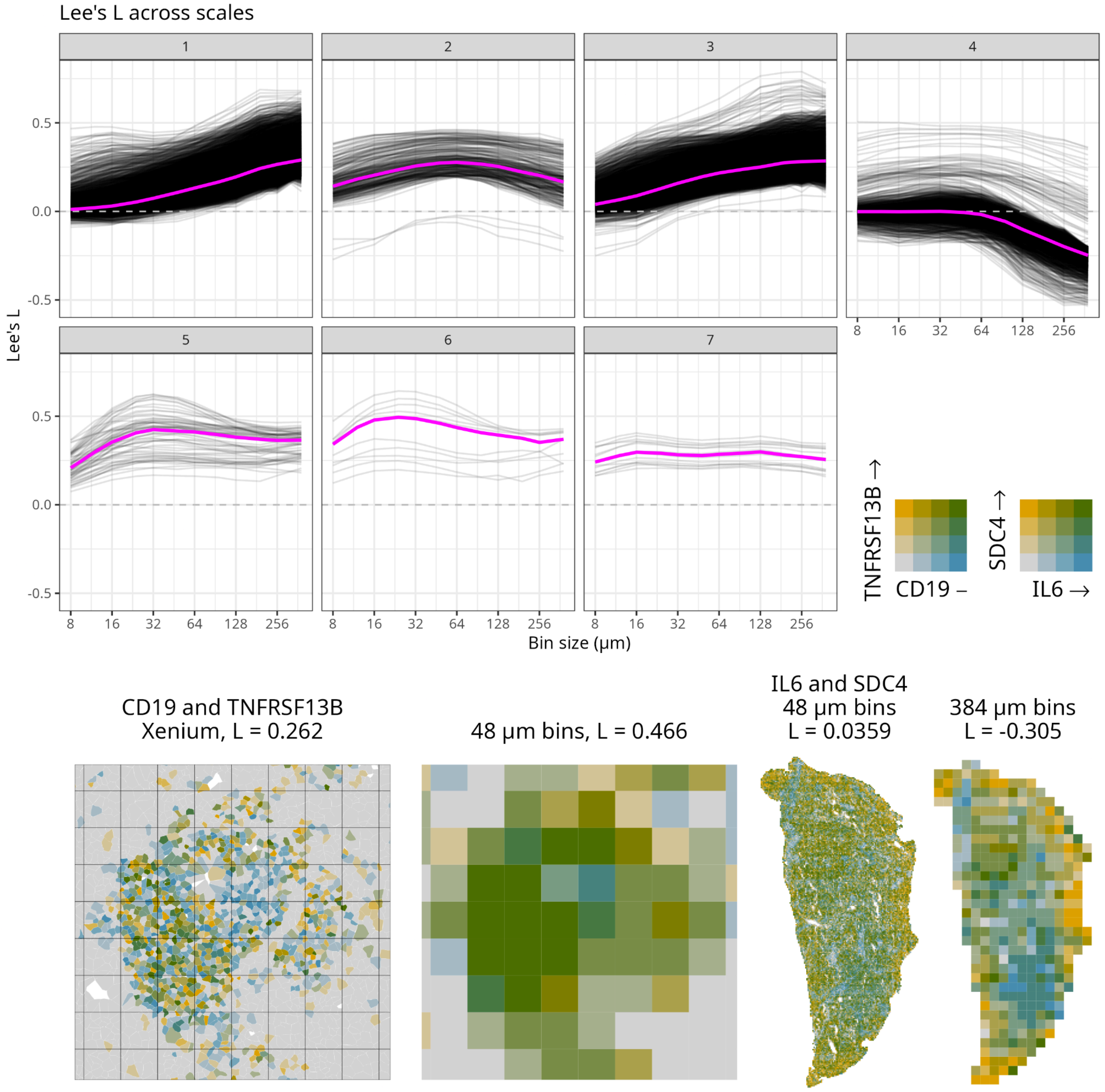


Supplementary Figure 8: Clusters of Lee's L curve patterns, along with one example of Lee's L becoming more positive with spatial aggregation and one example of Lee's L becoming more negative with spatial aggregation.

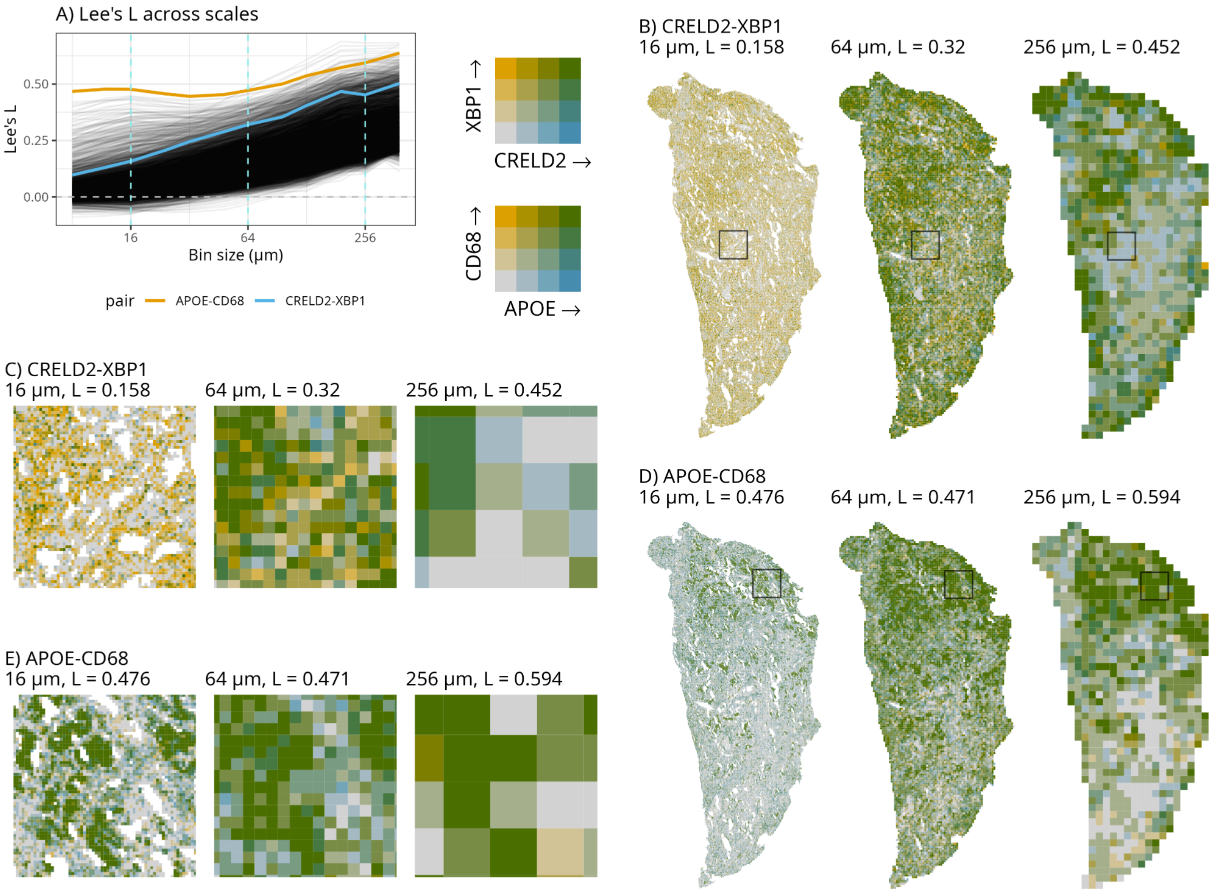


Supplementary Figure 9.1: Same as Figure 4 but for cluster 1 in Supplementary Figure 8; the example gene pair examples are CRELD2-XBP1 and APOE-CD68

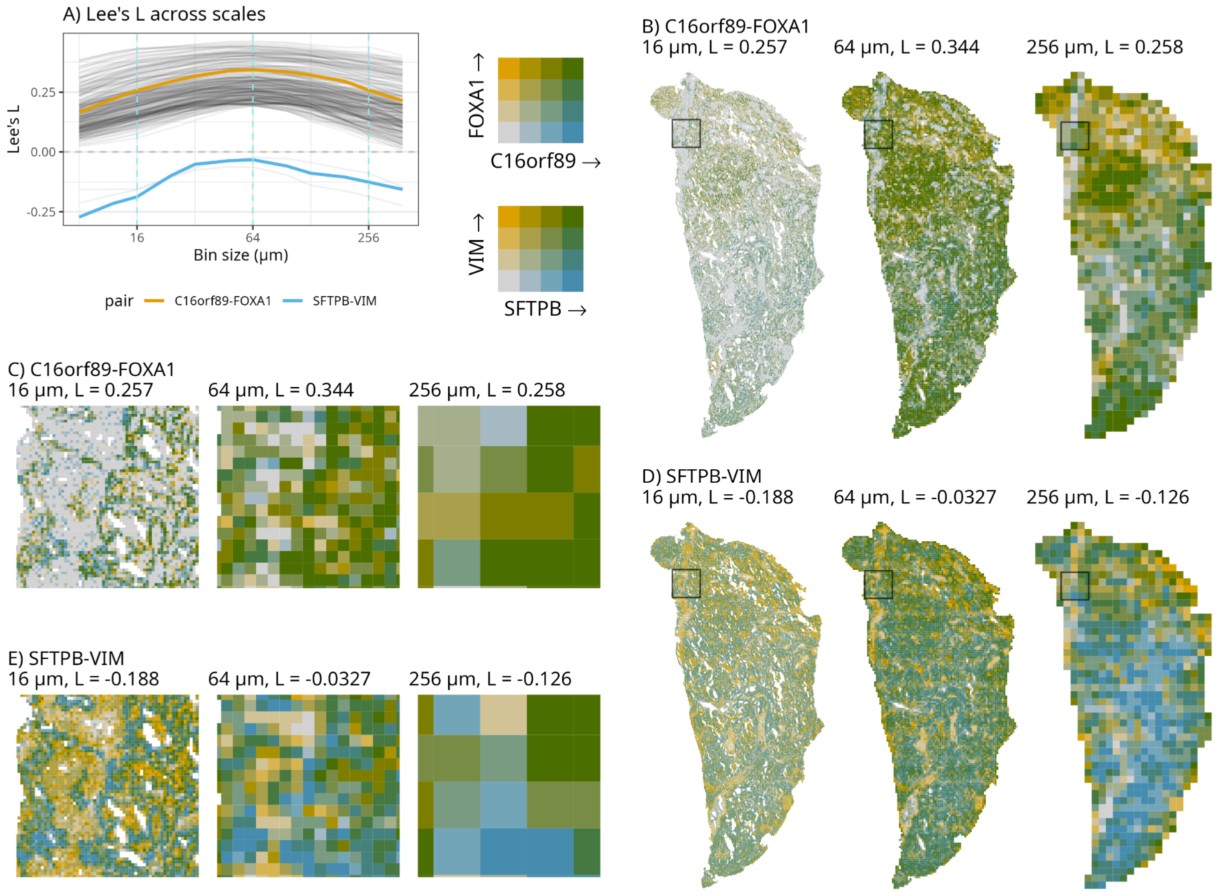


Supplementary Figure 9.2: Same as Figure 4 but for cluster 2 in Supplementary Figure 8, where Lee's L shows a peak at an intermediate scale. The two examples show two very different scenarios with this trend.

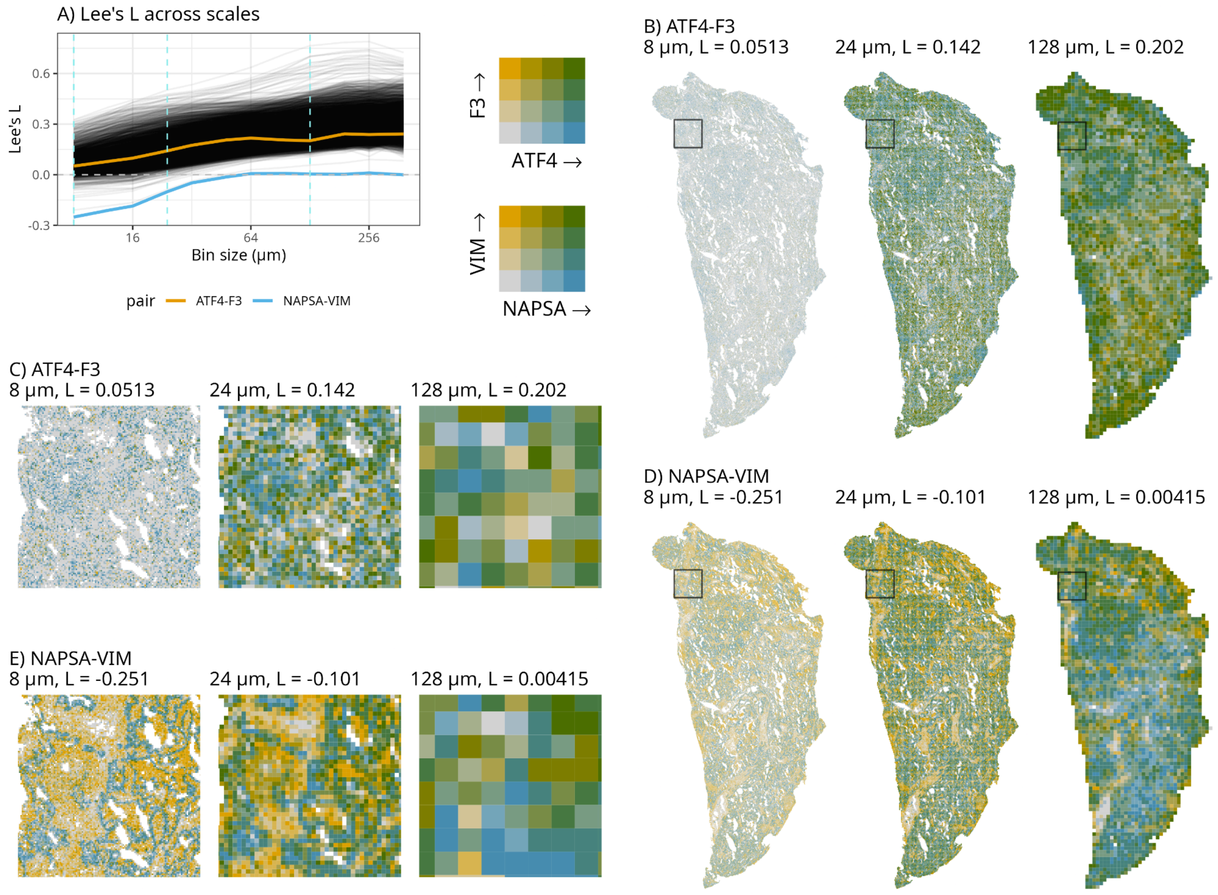


Supplementary Figure 9.3: Same as Figure 4 but for cluster 3 in Supplementary Figure 8

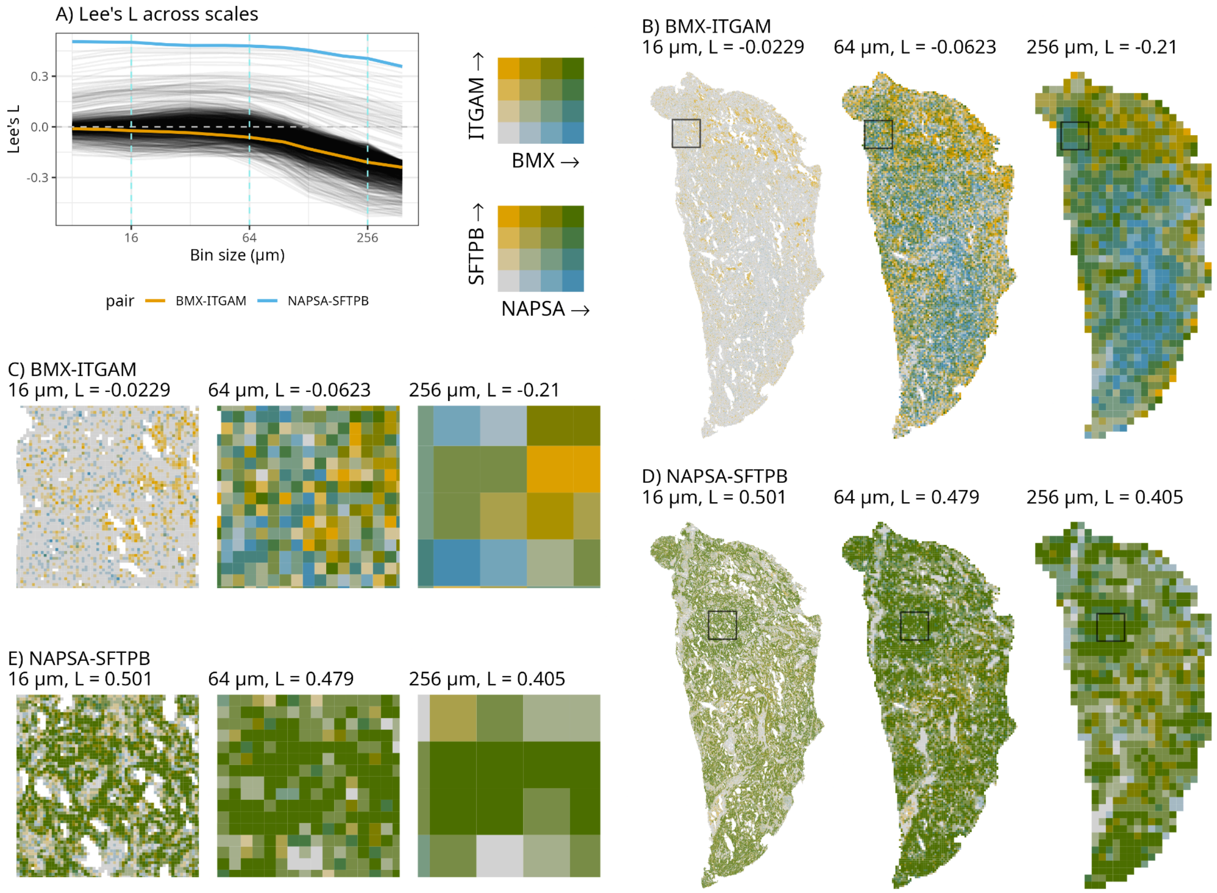


Supplementary 9.4: Same as Figure 4 but for cluster 4 in Supplementary Figure 8

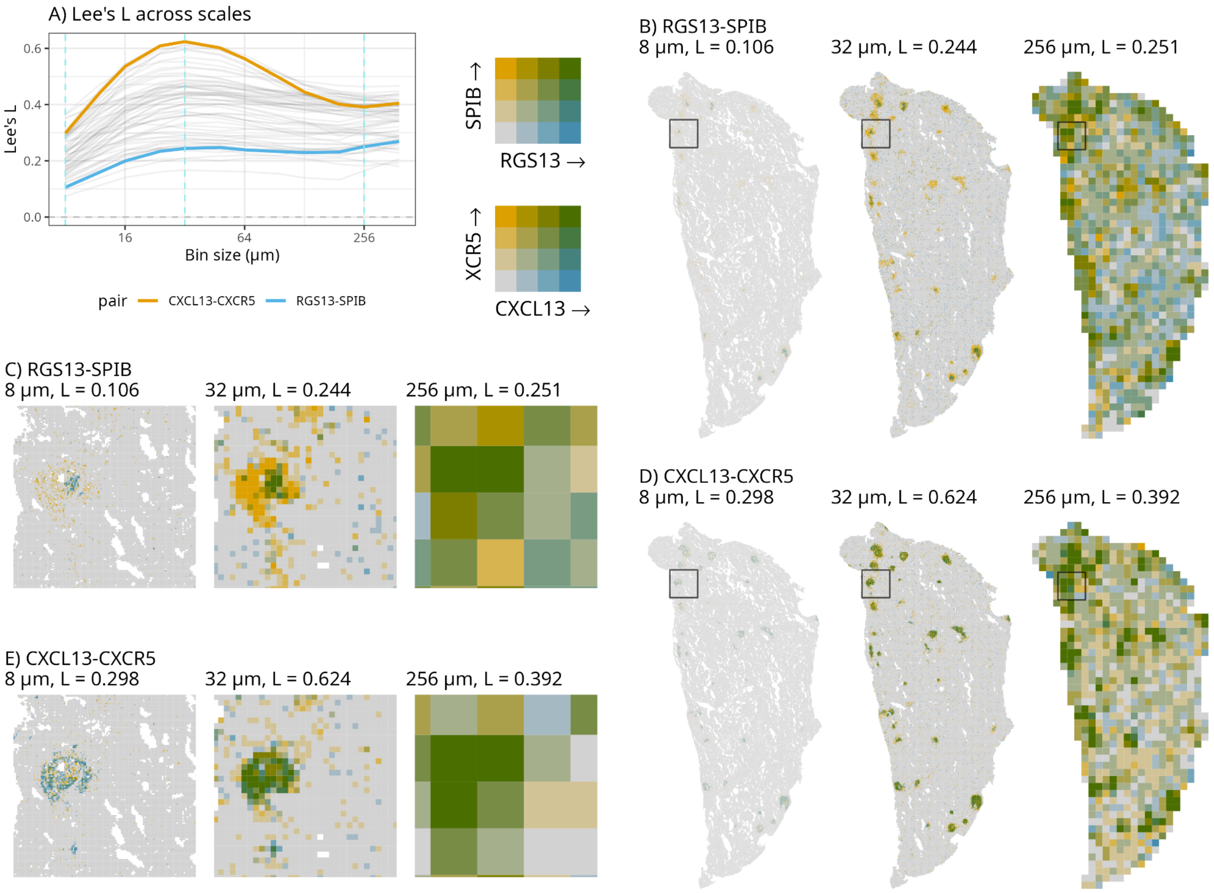


Supplementary Figure 9.5: Same as Figure 4 but for cluster 5 in Supplementary Figure 8

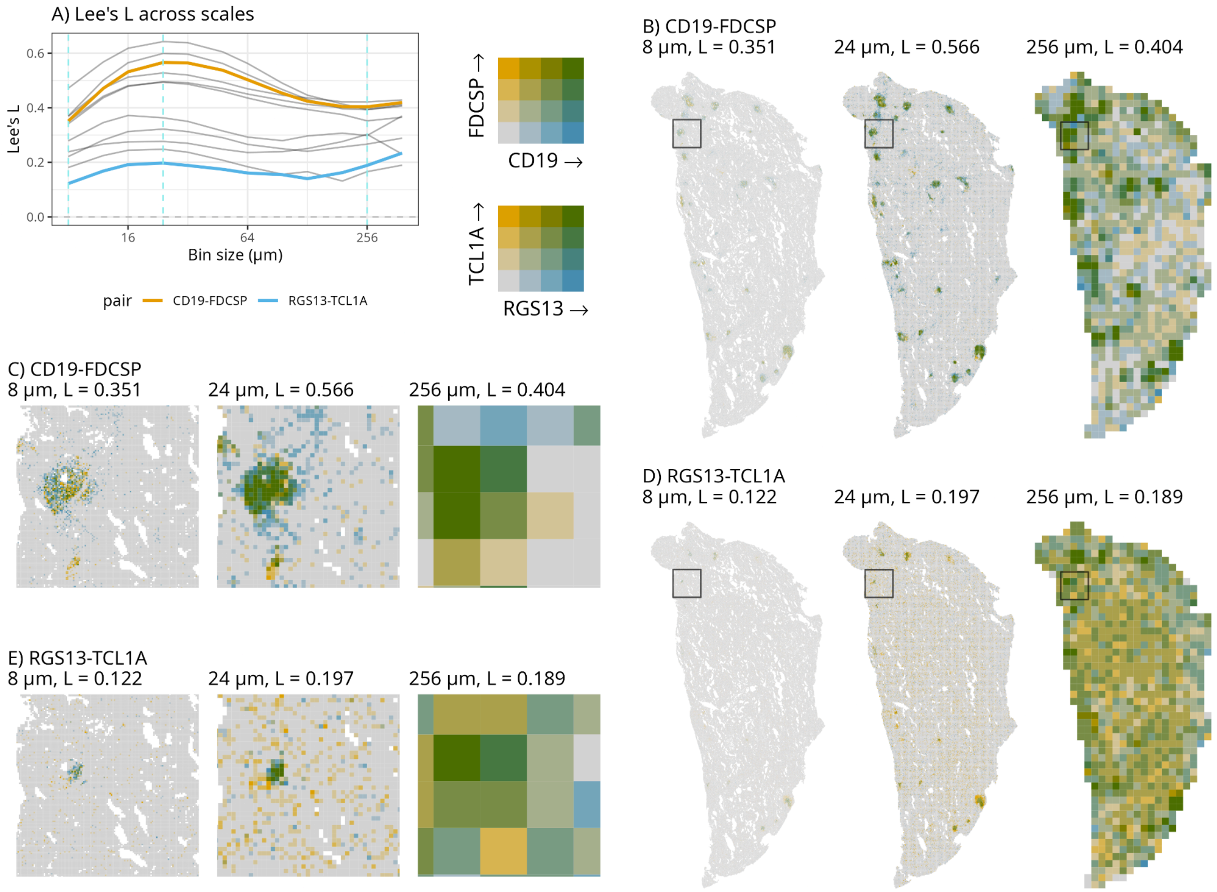


Supplementary Figure 9.6: Same as Figure 4 but for cluster 6 in Supplementary Figure 8

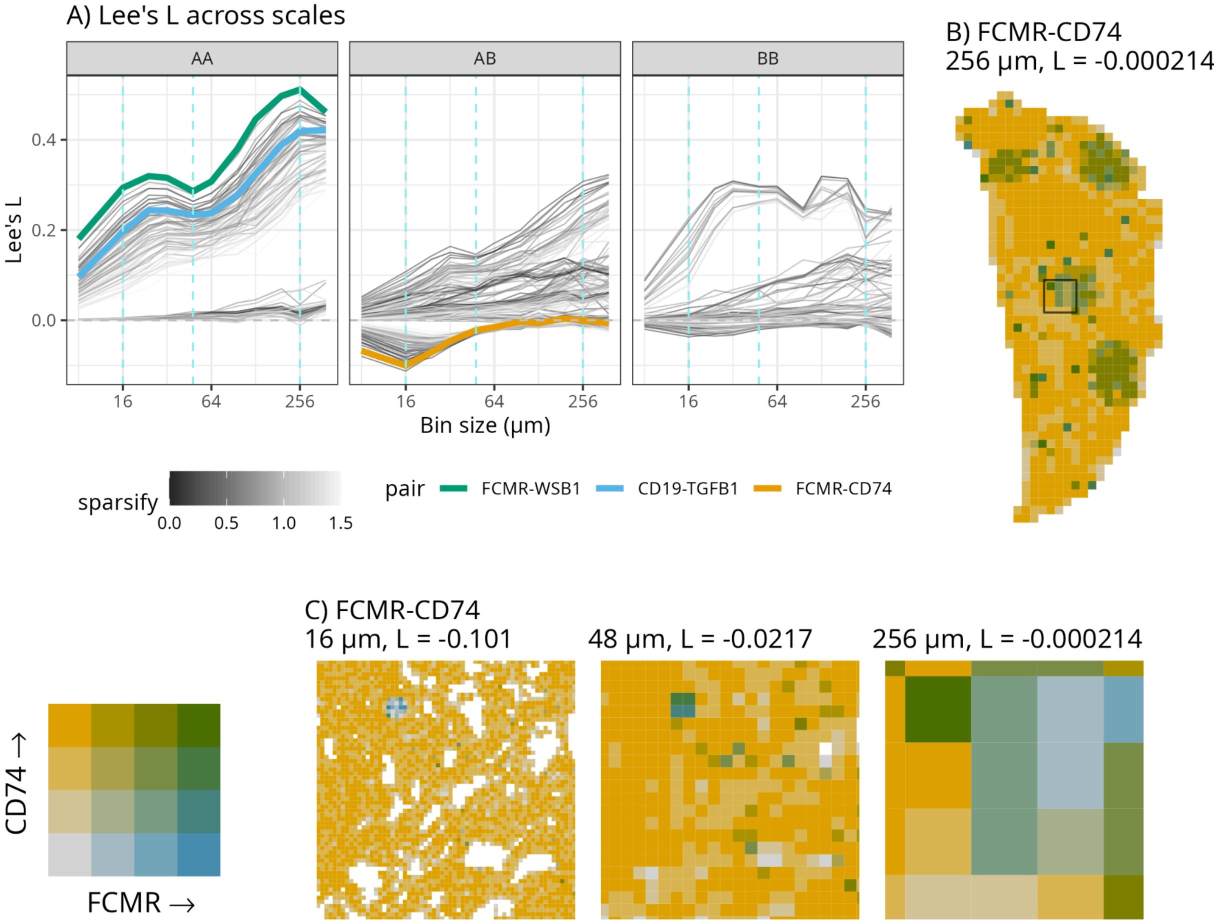


Supplementary Figure 10.1: Synthetic multivariate pattern example for Lee's L (two scales, bimodal). A) Lee's L vs. bin size, where each curve is for one pair of features. AA means correlation between features both designated to "gene program" A. AB means one feature is for "gene program" A and the other for B. BB means both are for B. These features have various strengths of correlations. In AA, the upper thick line is for a feature pair with strong correlation, and the lower thick line is for a feature pair with moderate correlation. Those with no correlation are around 0 at all scales. In AB, the thick line is for the top "marker" for A and top "marker" for B. The thin lines are not highlighted and are colored by the sum of the sparsification score (how much is it sparsified, see Methods) in both features, so fainter lines are for sparser gene pairs. B) Global plot of the feature pair highlighted in AB with bin size 256 µm, using a bivariate palette. C) Zooming into the box marked in B, showing 16, 48, and 256 µm bins.

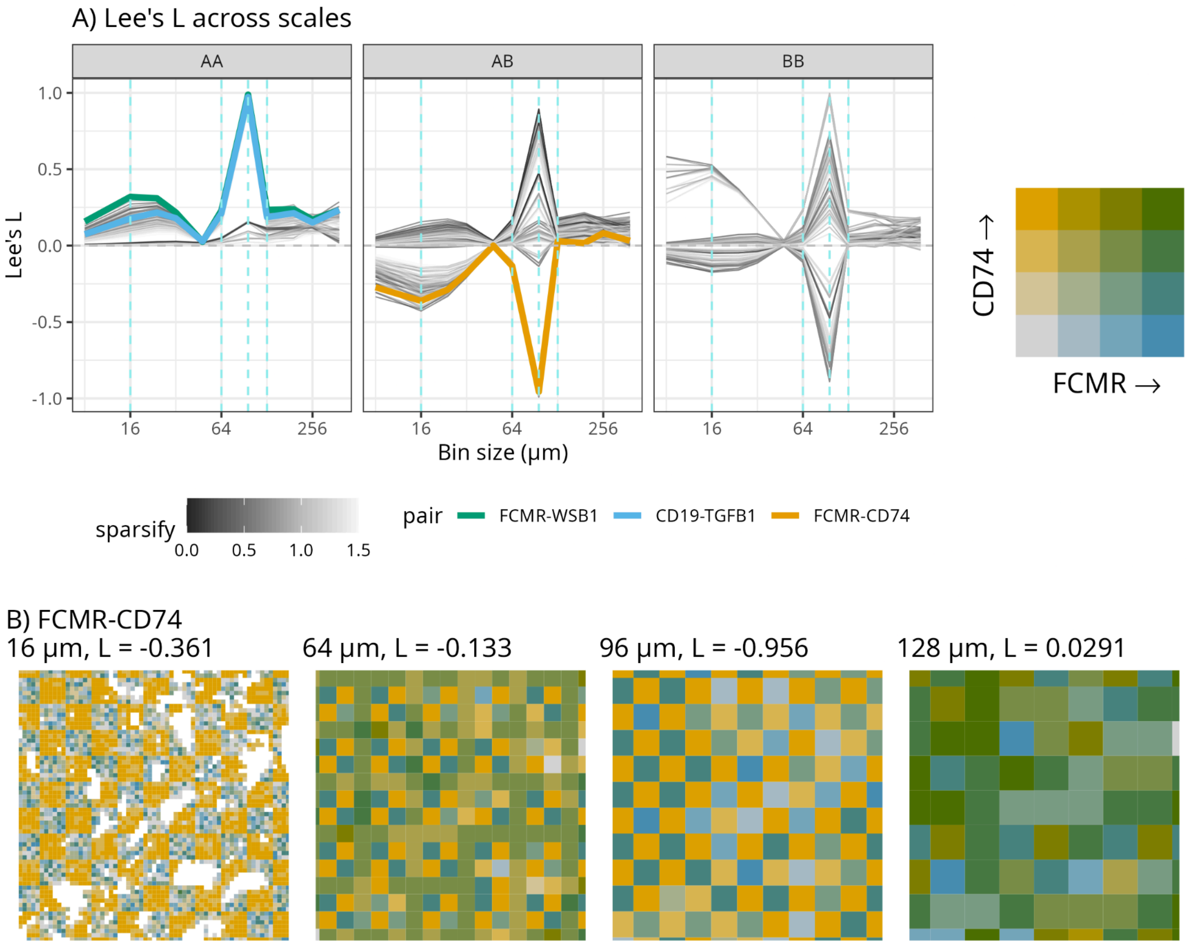


Supplementary Figure 10.2: Same as Supplementary Figure 11.1 but for a pattern where “cell types” A and B are arranged in a checkerboard pattern with side length 96. Only the local plots in a 1000 μm x 1000 μm box is shown.

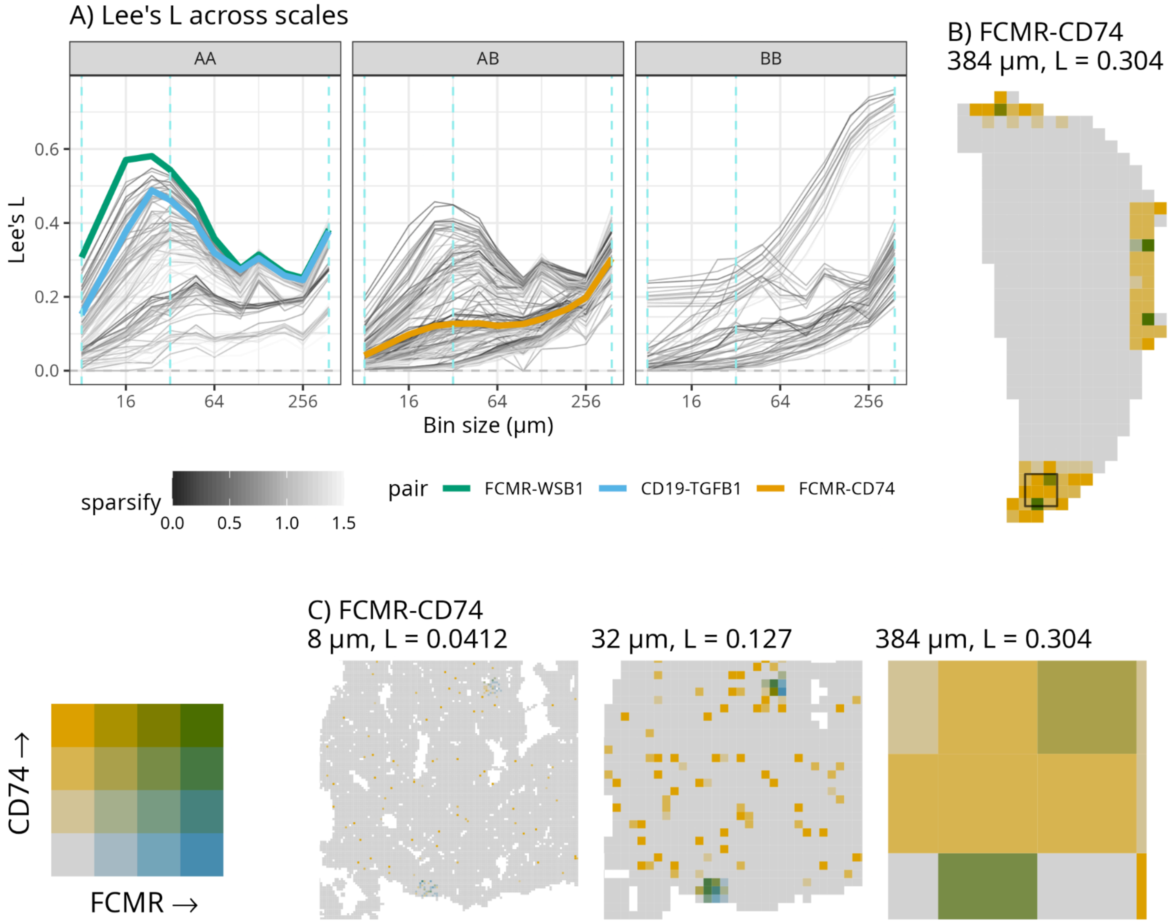


Supplementary Figure 10.3: Same as Supplementary Figure 11.1 but for a pattern in which both features are sparse and confined to small regions of the section. “Cell type” A is in small spots located in those regions, while “cell type” B is sparsely and randomly distributed in those regions.

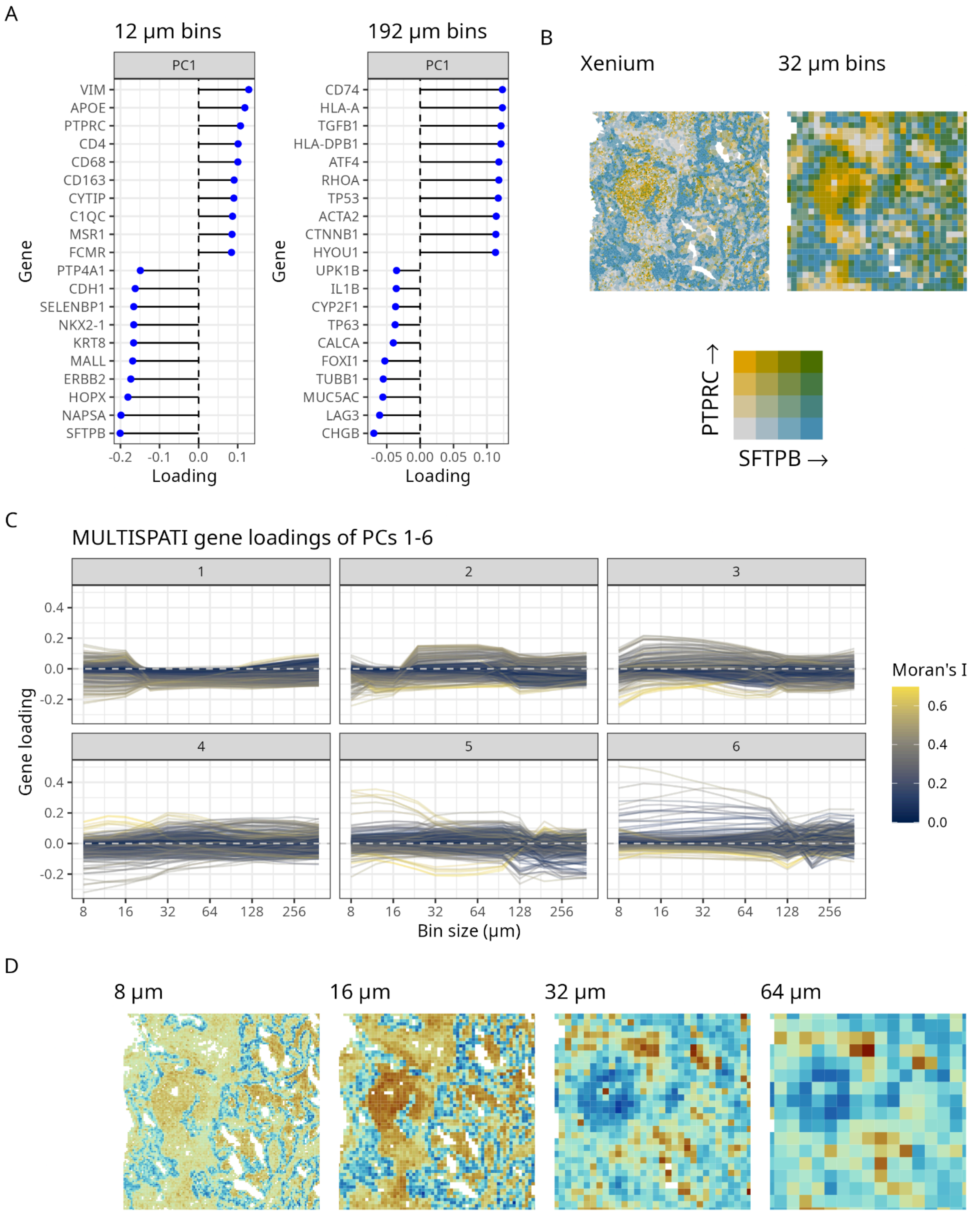


Supplementary Figure 11: Changes of MULTISPATI PCA results in TSU-21 with bin sizes. A) PC1 loadings of genes with the largest loadings for 12 μm and 192 μm bins. B) Bivariate plot of SFTPB and PTPRC in a 1000 μm x 1000 μm box in Xenium (single cell) and 32 μm bins. C) Loadings of each gene across bin sizes are plotted as a curve, for PC1-6. The curves are colored by Moran's I of the gene in 48 µm bins. D) PC1 loadings shown in the same box as in B; warm colors are positive values and cold colors are negative values.

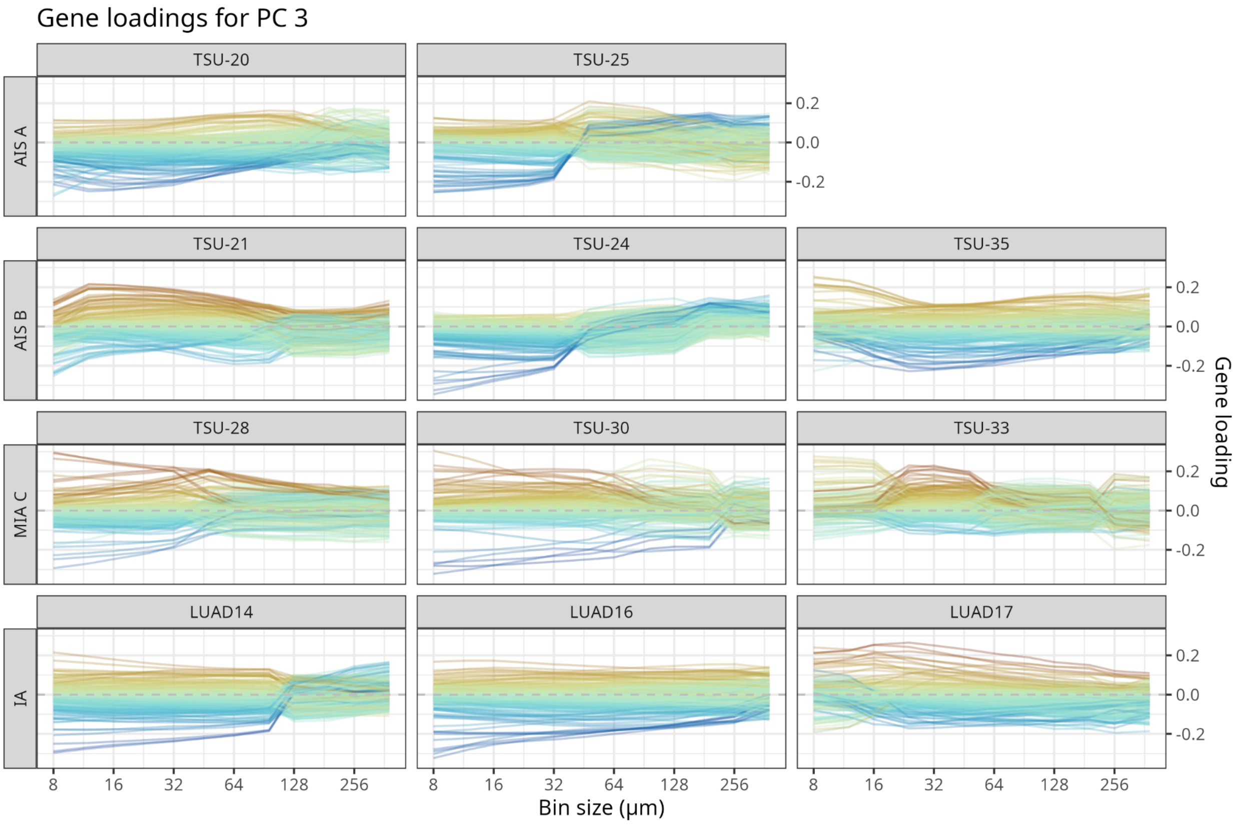


Supplementary Figure 12: MULTISPATI PC3 loadings vs. bin size in all samples across stages. Each gene has a curve, and the curves are colored by loadings in 32 µm bins in each sample. While "phase shifts" can be observed, they seem inconsistent between samples across samples within the same stage.

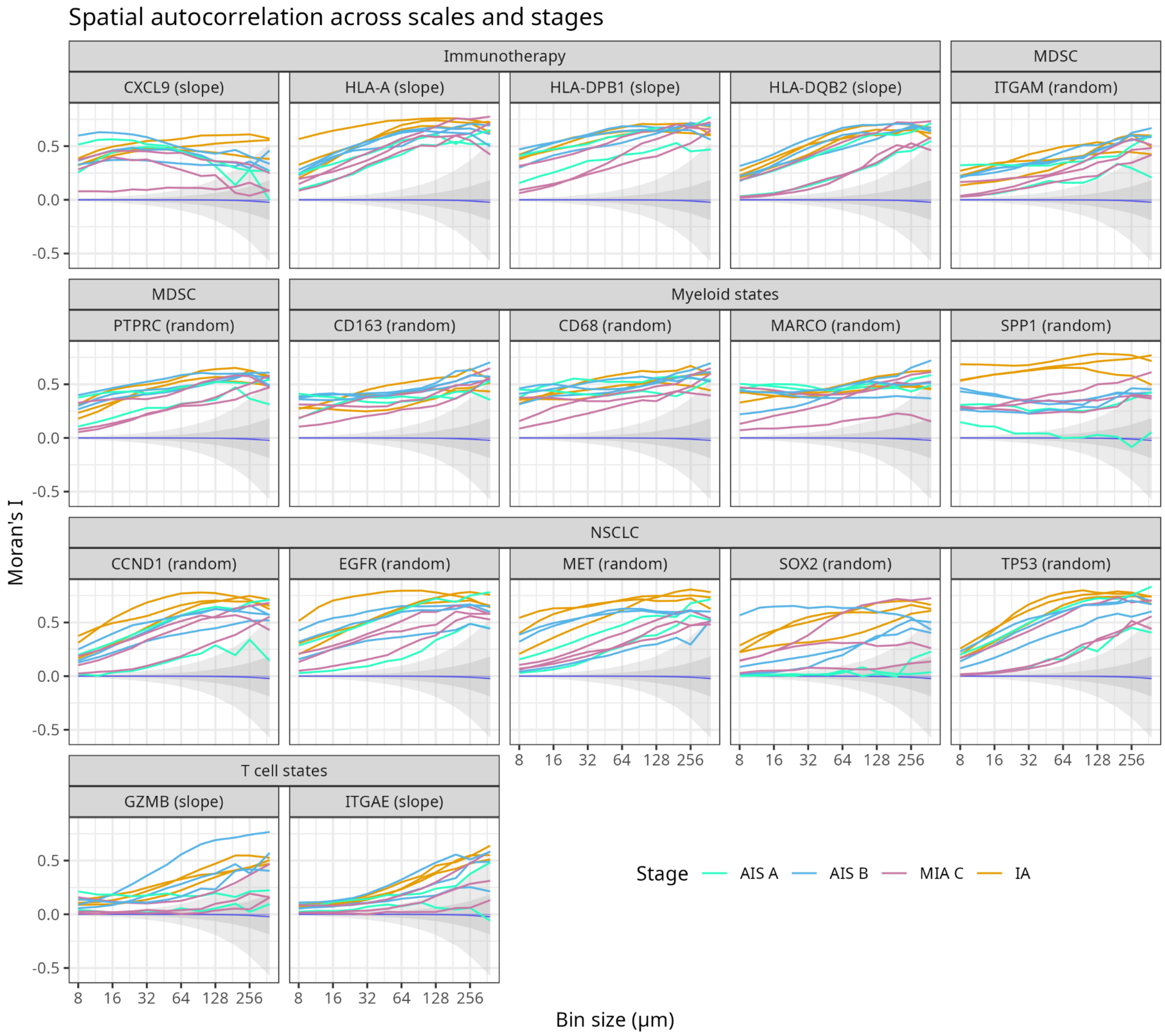


Supplementary Figure 13: Moran's I curves across stages for genes of interest. The upper strip shows the biological function the gene is relevant to. The gray ribbons are the same as in Figure 5B. In the parentheses in facet labels, "random" means the gene has significant random effect after correcting for multiple testing but not significant random slope, and "slope" means the gene has significant random slope after correcting for multiple testing.

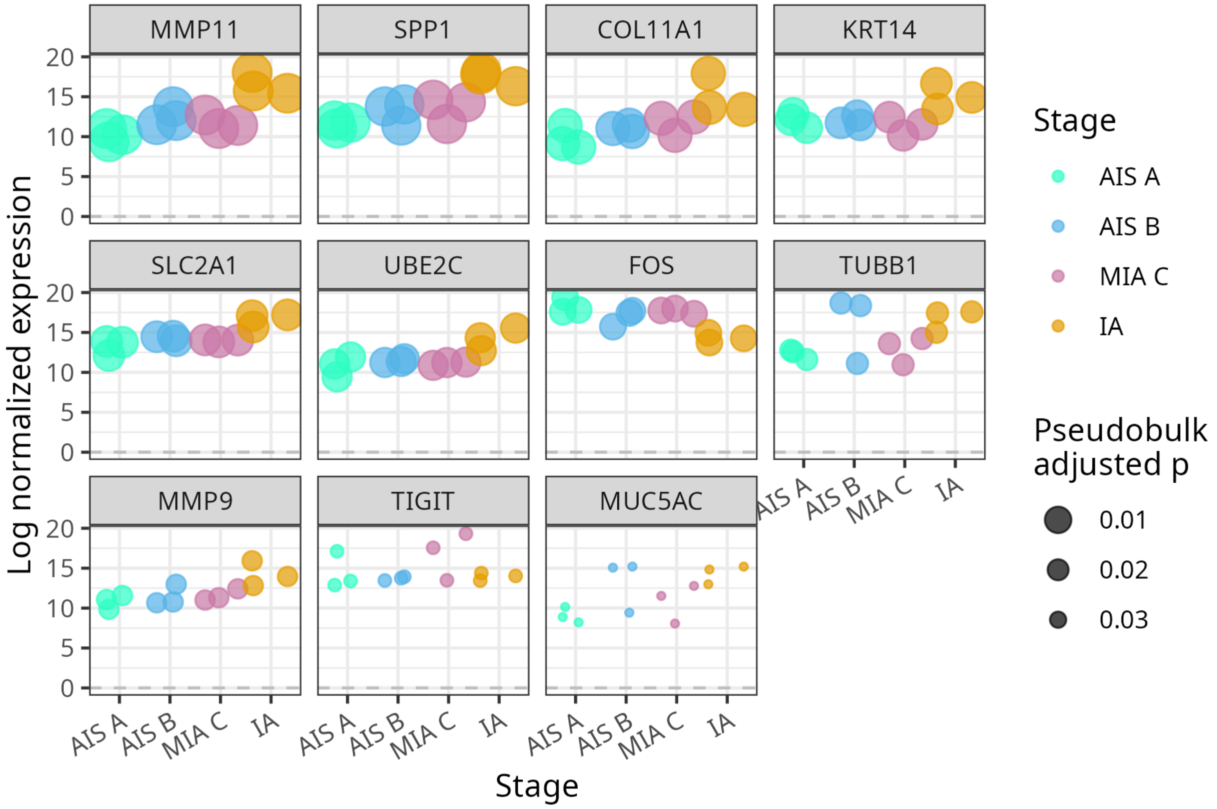


Supplementary Figure 14: Log normalized expression of each pseudobulk DE gene. Each point represents a sample, and point size corresponds to adjusted p-value of the gene in pseudobulk DE.

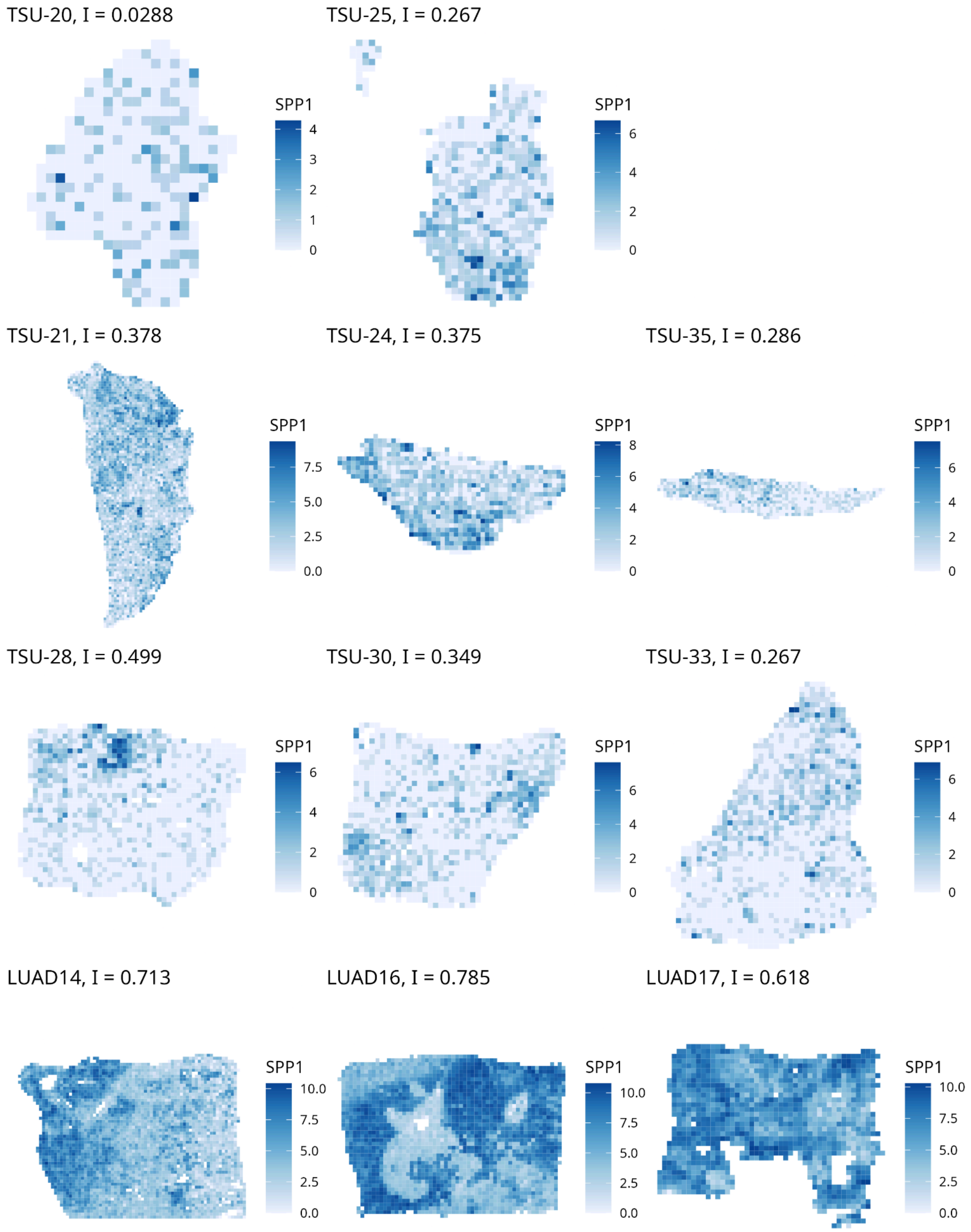

TSU-20, I = 0.0288
SPP1
4
3
2
1
0
TSU-25, I = 0.267
SPP1
6
4
2
0
TSU-21, I = 0.378
SPP1
7.5
5.0
2.5
0.0
TSU-24, I = 0.375
SPP1
8
6
4
2
0
TSU-35, I = 0.286
SPP1
6
4
2
0
TSU-28, I = 0.499
SPP1
6
4
2
0
TSU-30, I = 0.349
SPP1
6
4
2
0
TSU-33, I = 0.267
SPP1
6
4
2
0
LUAD14, I = 0.713
SPP1
10.0
7.5
5.0
2.5
0.0
LUAD16, I = 0.785
SPP1
10.0
7.5
5.0
2.5
0.0
LUAD17, I = 0.618
SPP1
10.0
7.5
5.0
2.5
0.0

Supplementary Figure 15.1: Spatial patterns of log normalized SPP1 expression in all 11 samples, with 128 μm bins.

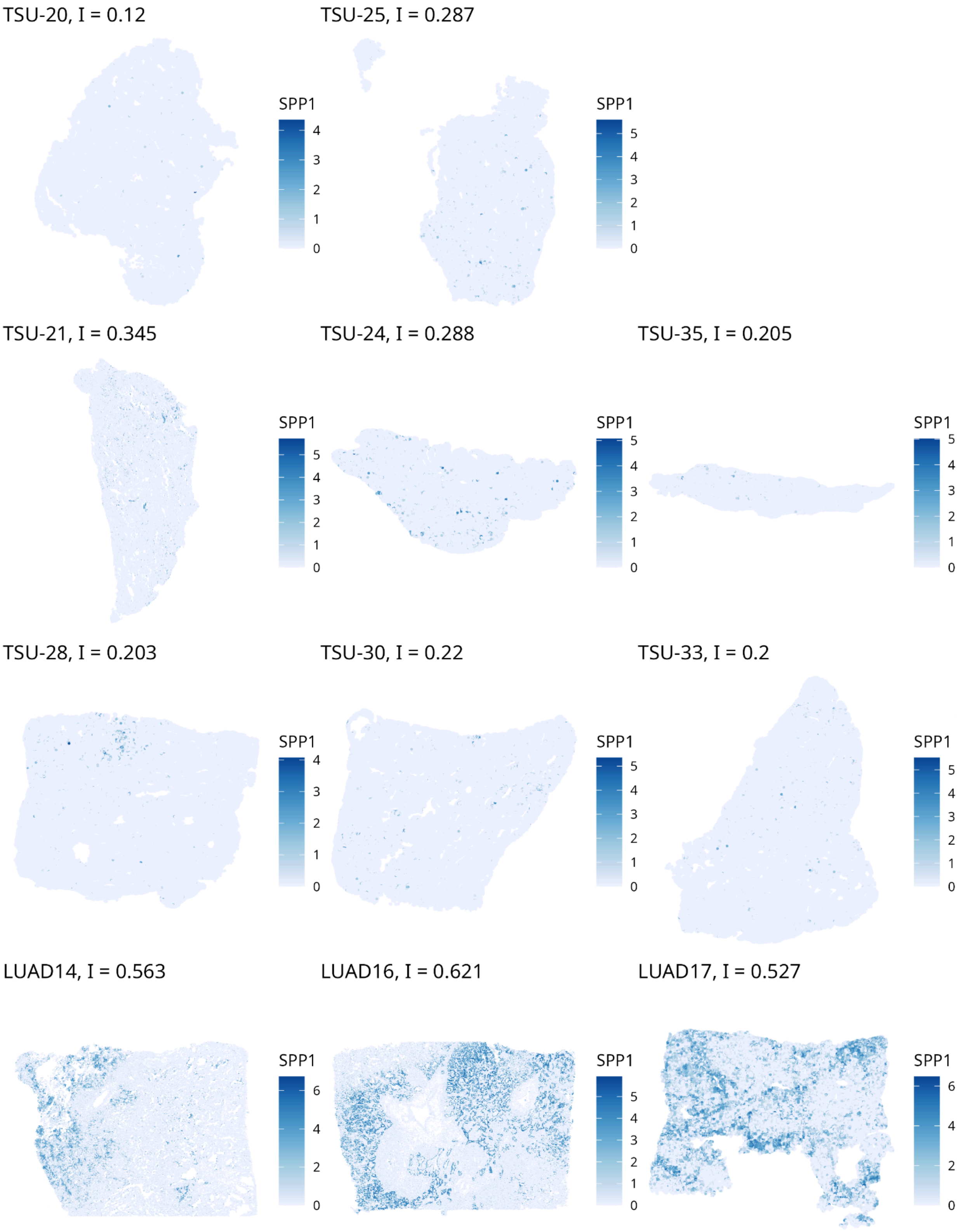


Supplementary Figure 15.2: Spatial pattern of SPP1 in all samples plotted on Xenium cells.

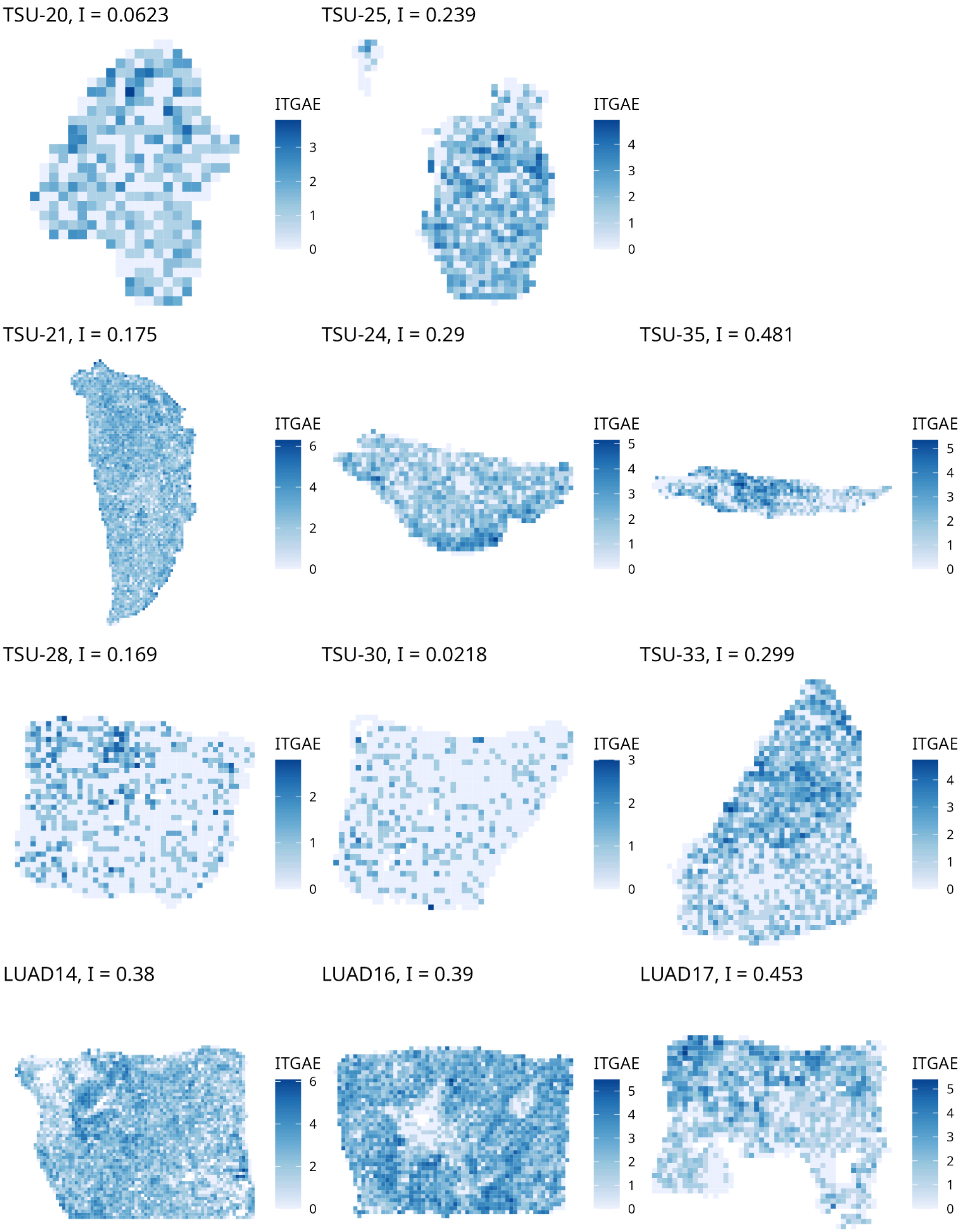


Supplementary Figure 15.3: Spatial pattern of ITGAE in all samples on 128 µm bins.

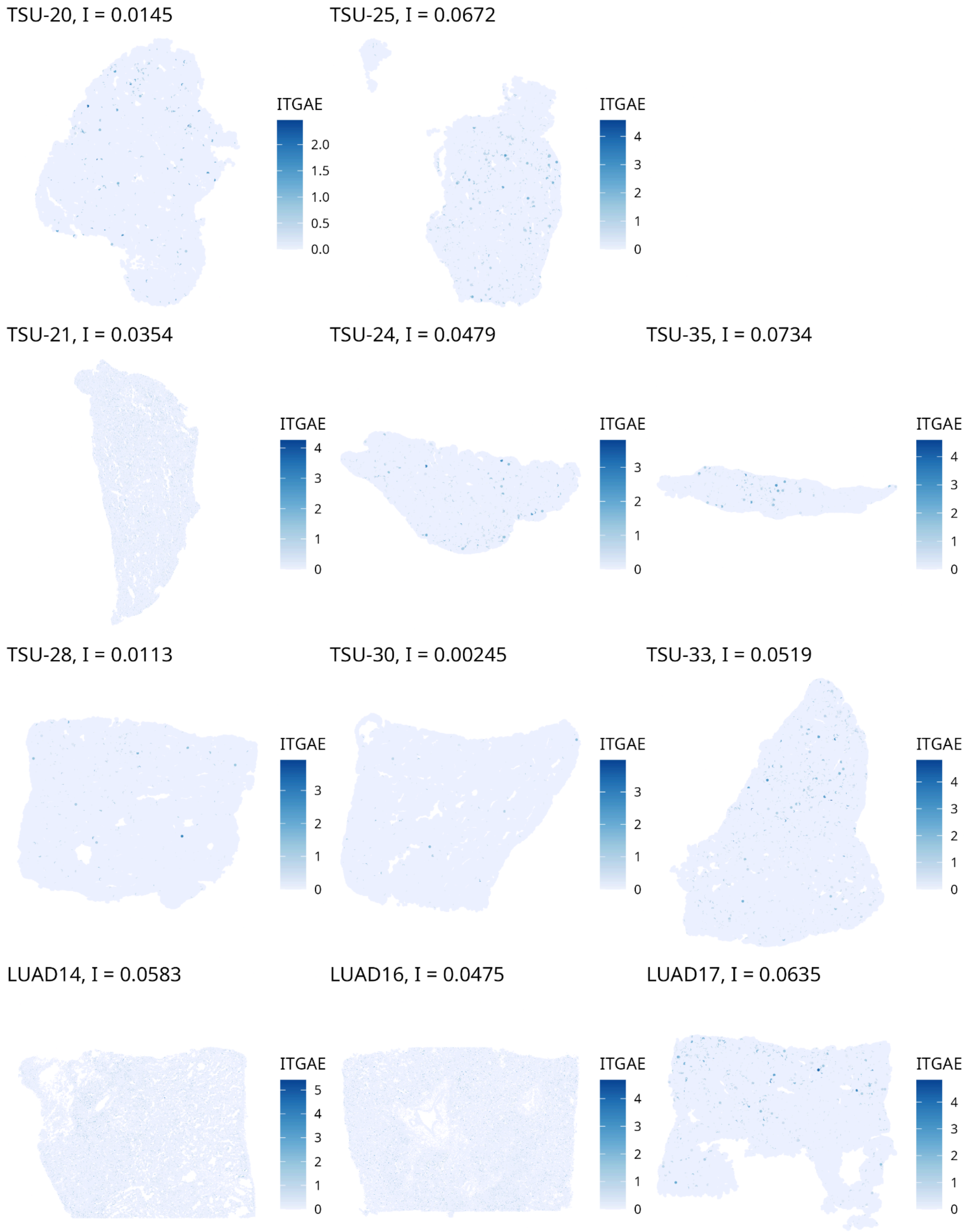


Supplementary Figure 15.4: Spatial pattern of ITGAE in all samples on Xenium cells.

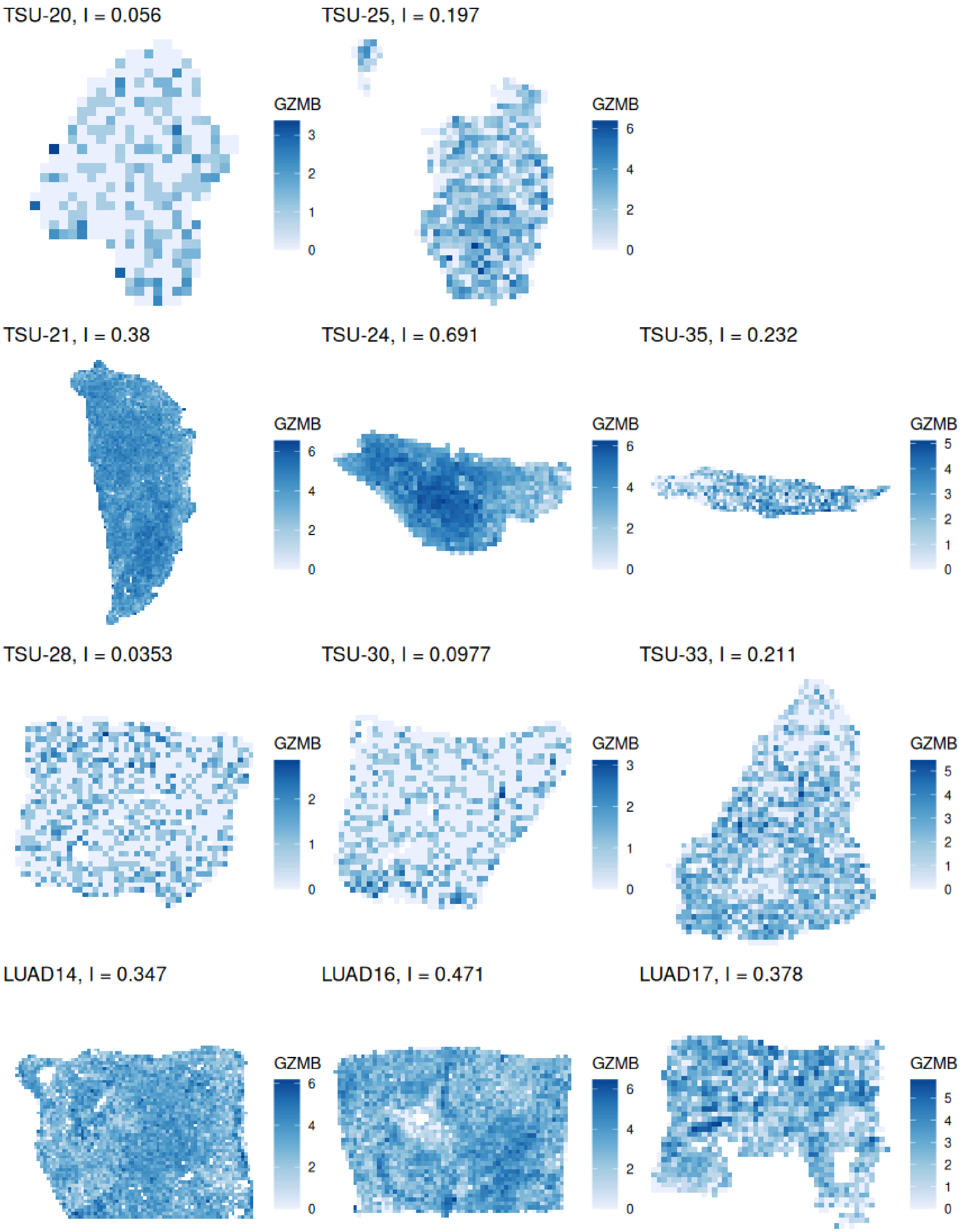


Supplementary Figure 15.5: Spatial pattern of GZMB in all samples on 128 μm bins.

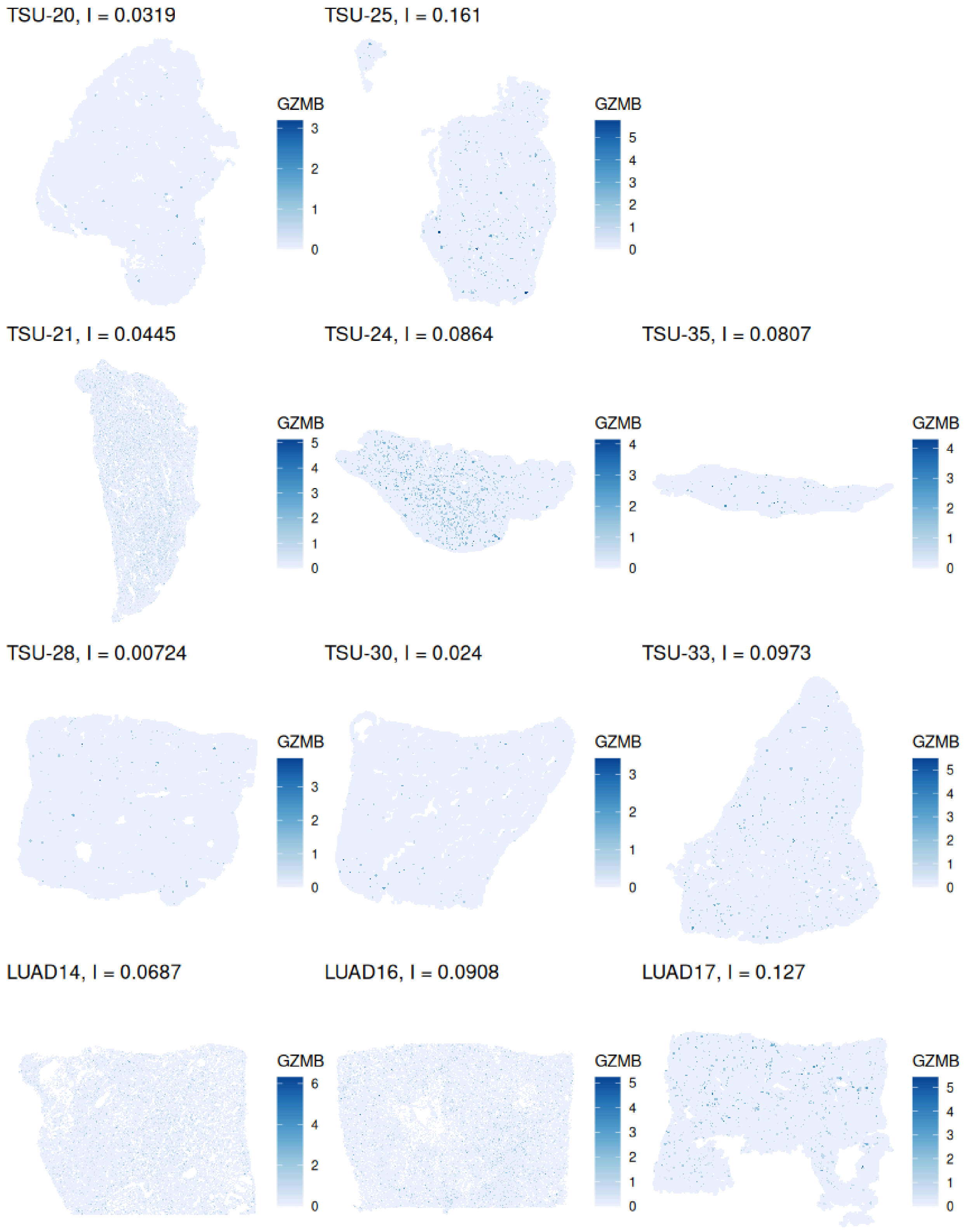


Supplementary Figure 15.6: Spatial pattern of GZMB in all samples on Xenium cells.

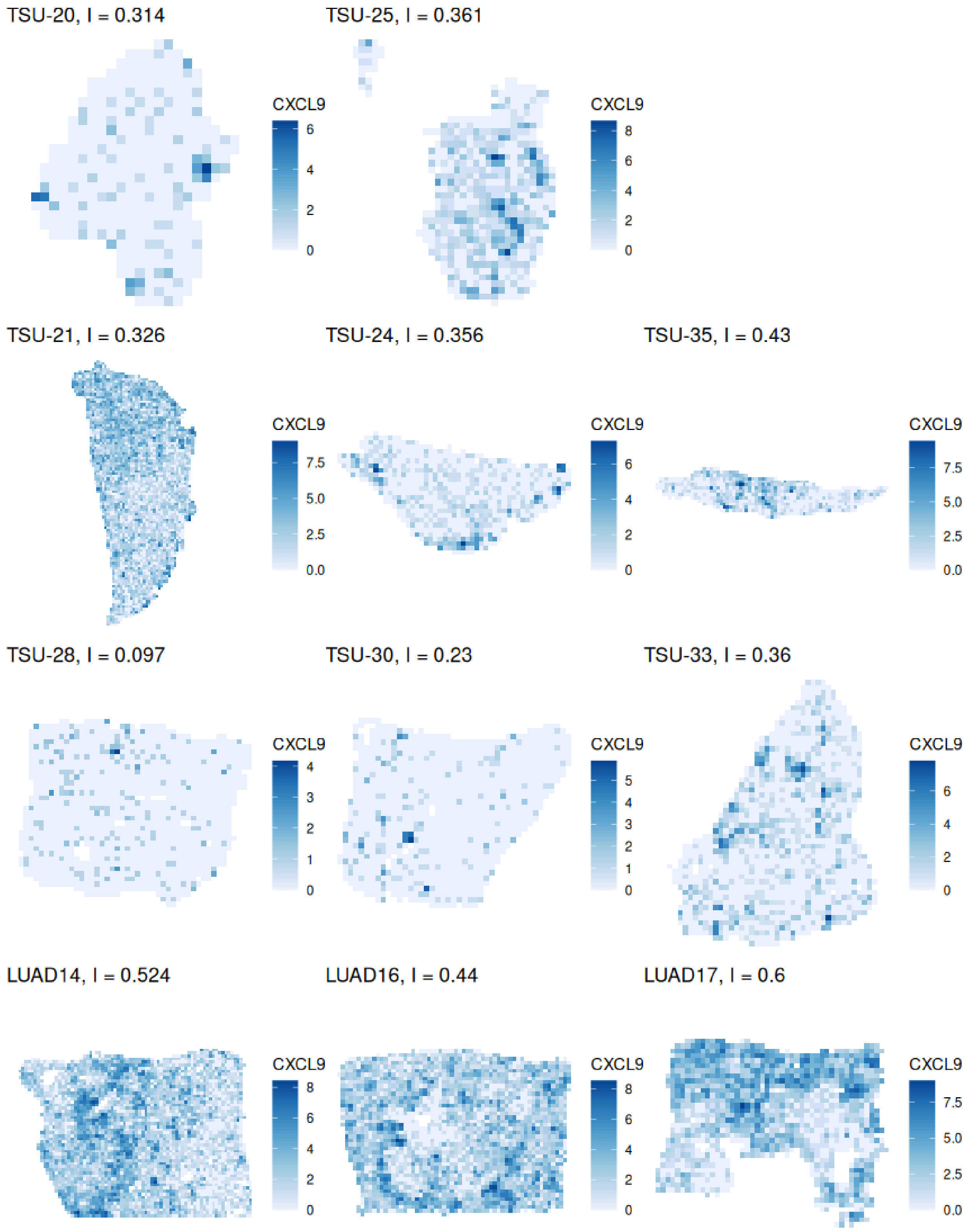


Supplementary Figure 15.7: Spatial pattern of CXCL9 in all samples plotted on 128 μm bins.

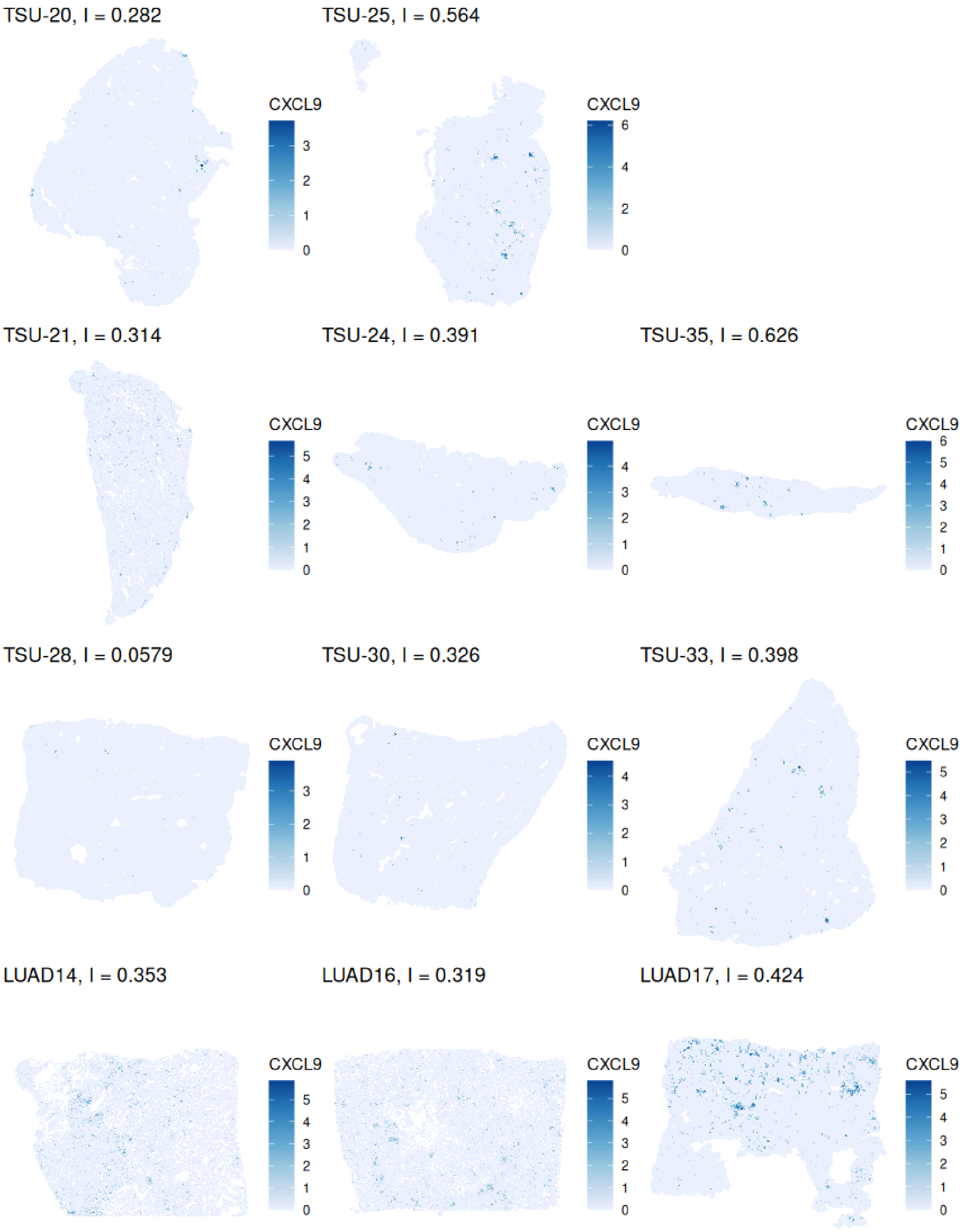


Supplementary Figure 15.8: Spatial pattern of CXCL9 in all samples plotted on Xenium cells.

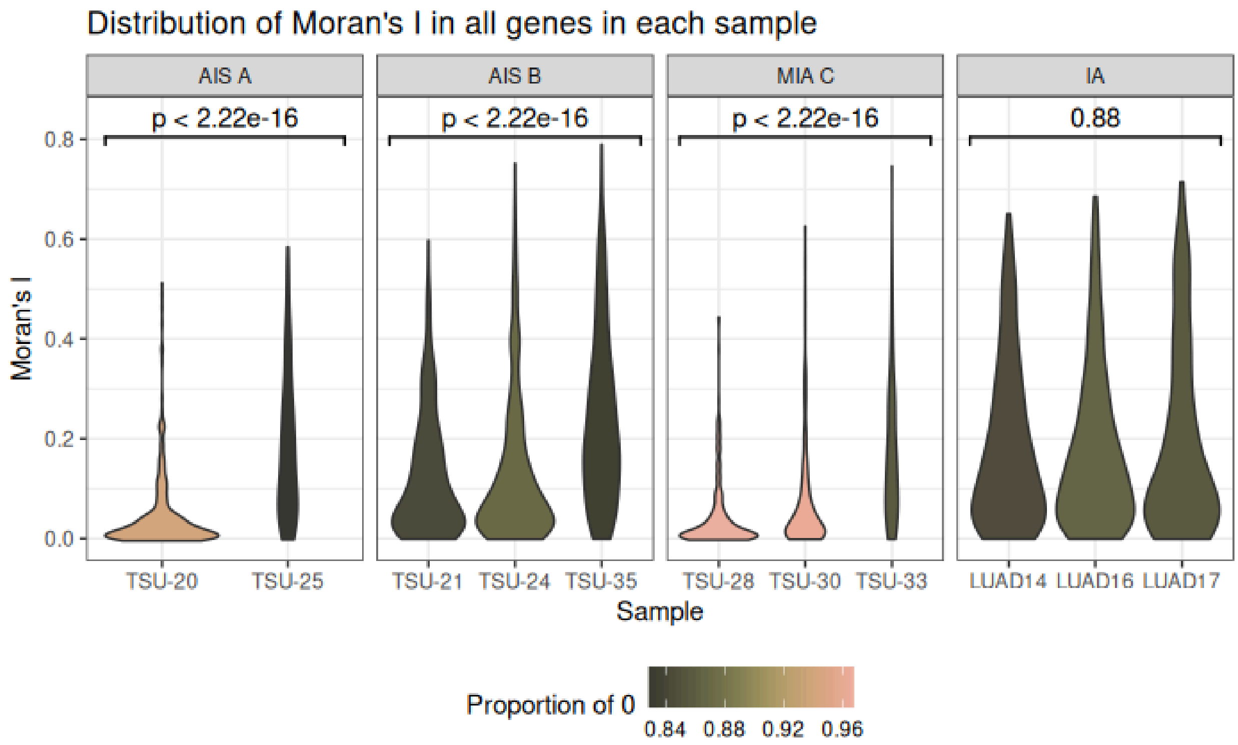


Supplementary Figure 16: Violin plot of distribution of Moran's I across all 302 genes in each sample, grouped by LUAD stage. The violins are colored by the proportion of 0's in the raw count matrix of the sample. The p-values are from the Kruskal-Wallis test; in AIS A, AIS B, and MIA C, $p < 2.22e-16$, indicating that different samples have different distributions, which seems to be associated with proportion of 0's; this is not significant in IA.

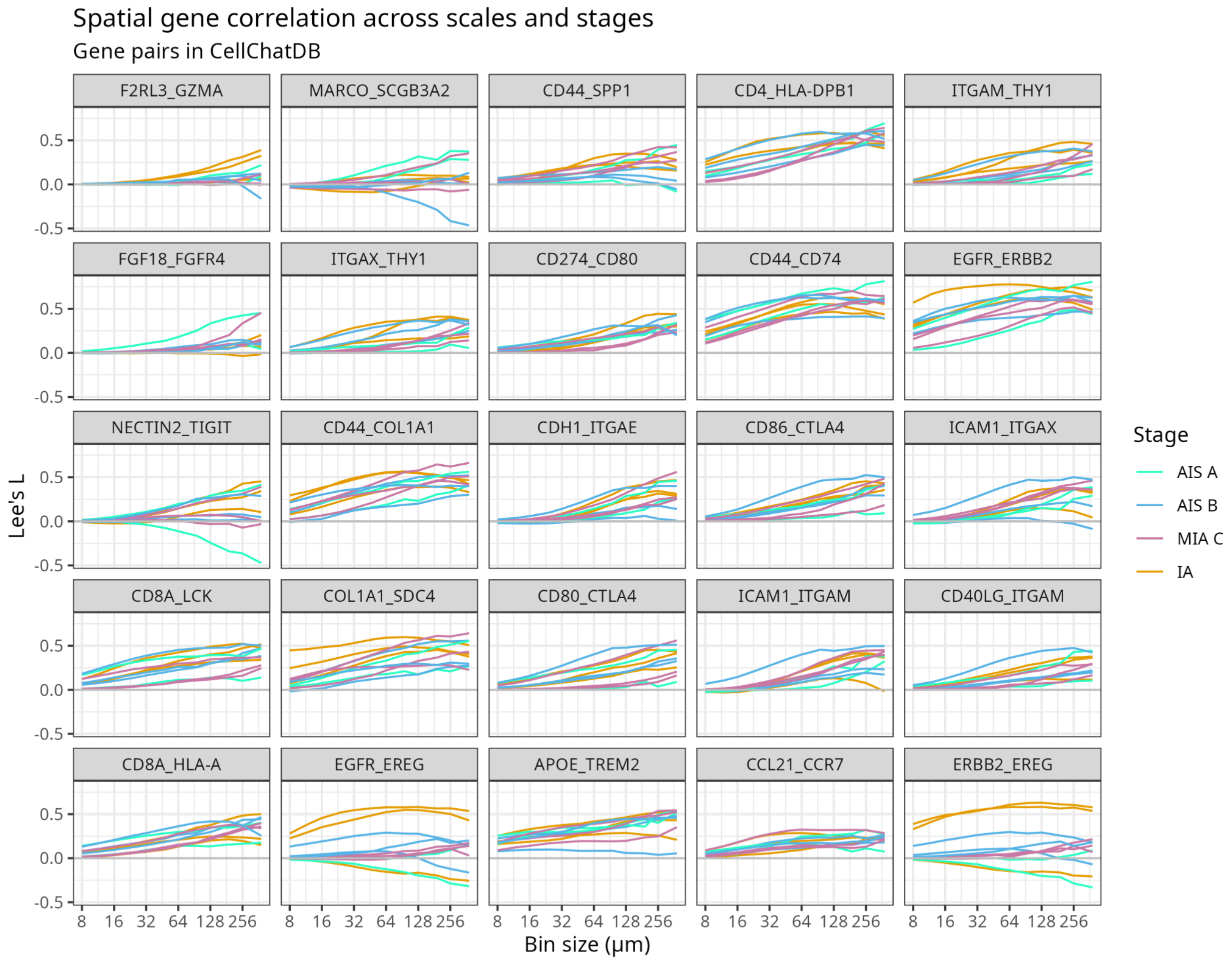


Supplementary Figure 17: Lee's L curves for gene pairs in Cell Chat's database

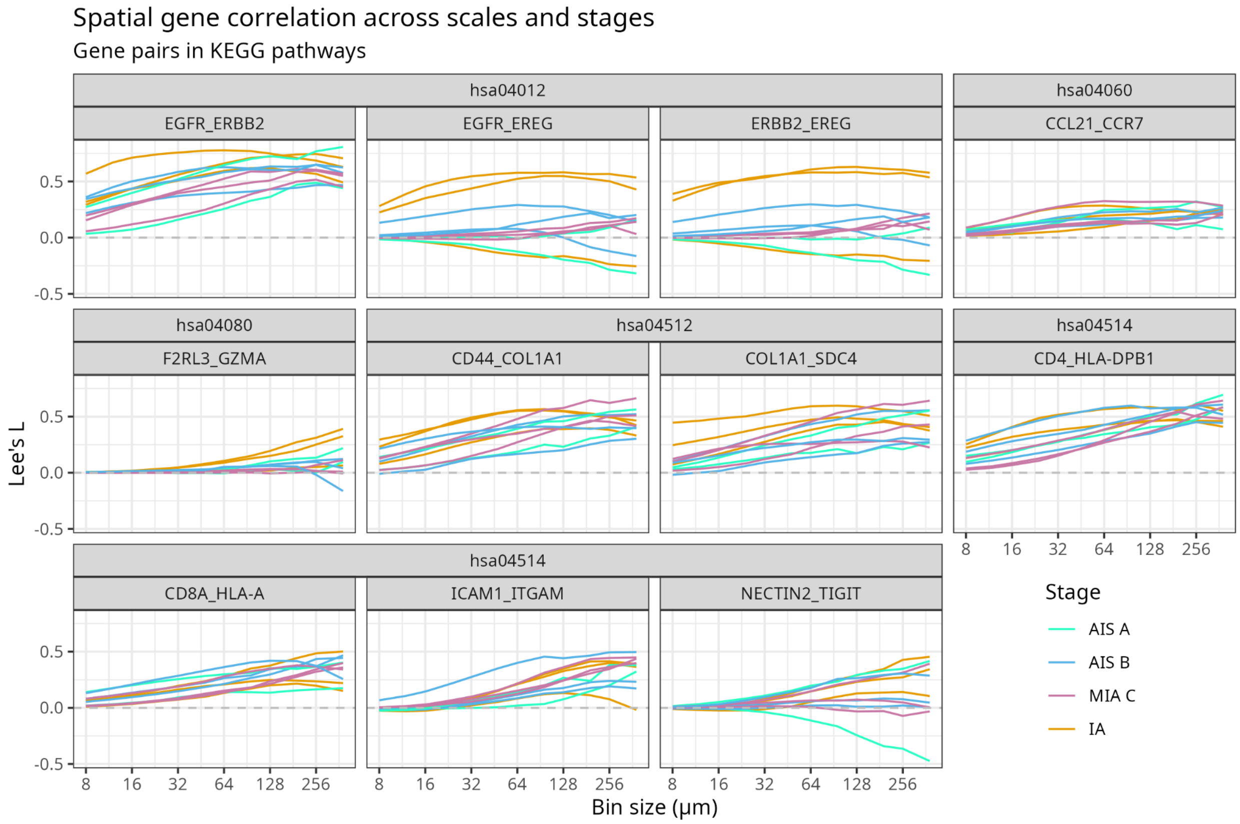


Supplementary Figure 18: Lee's L curves for gene pairs in KEGG pathways. hsa04012 is ERBB signaling; hsa04060 is Cytokine-cytokine receptor interaction; hsa04080 is Neuroactive ligand-receptor interaction; hsa04512 is ECM-receptor interaction; hsa04514 is cell adhesion molecule interactions.

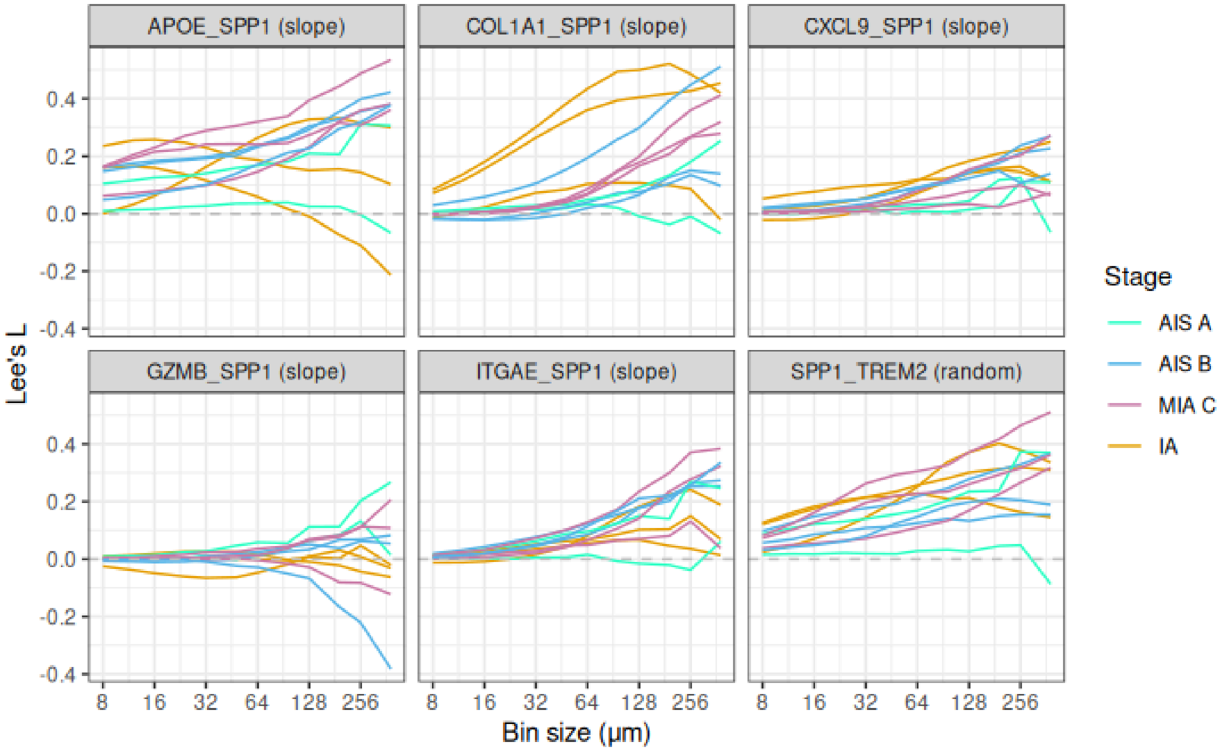


Supplementary Figure 19: Lee’s L curves across stages for gene pairs of interest involving SPP1 and markers of myeloid cells, T cells, and cancer progression.

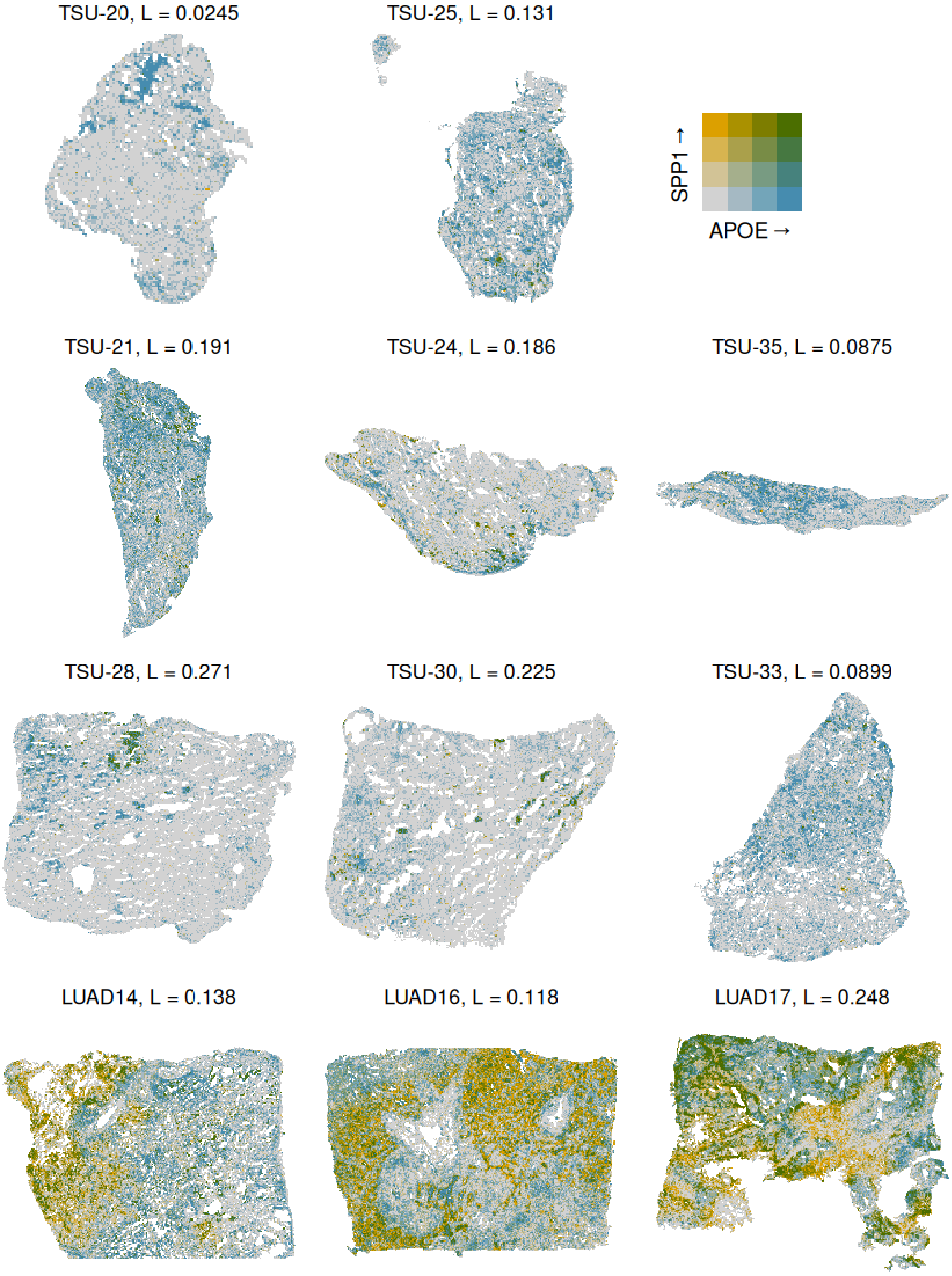

TSU-20, L = 0.0245
TSU-25, L = 0.131
SPP1 →
APOE →
TSU-21, L = 0.191
TSU-24, L = 0.186
TSU-35, L = 0.0875
TSU-28, L = 0.271
TSU-30, L = 0.225
TSU-33, L = 0.0899
LUAD14, L = 0.138
LUAD16, L = 0.118
LUAD17, L = 0.248

Supplementary Figure 20.1: Bivariate plot showing spatial co-expression of SPP1 and APOE in all tissues with 24 µm bins; yellow means high in SPP1 and low in APOE. Blue means high in APOE and low in SPP1. Green means high in both. Gray means low in both.

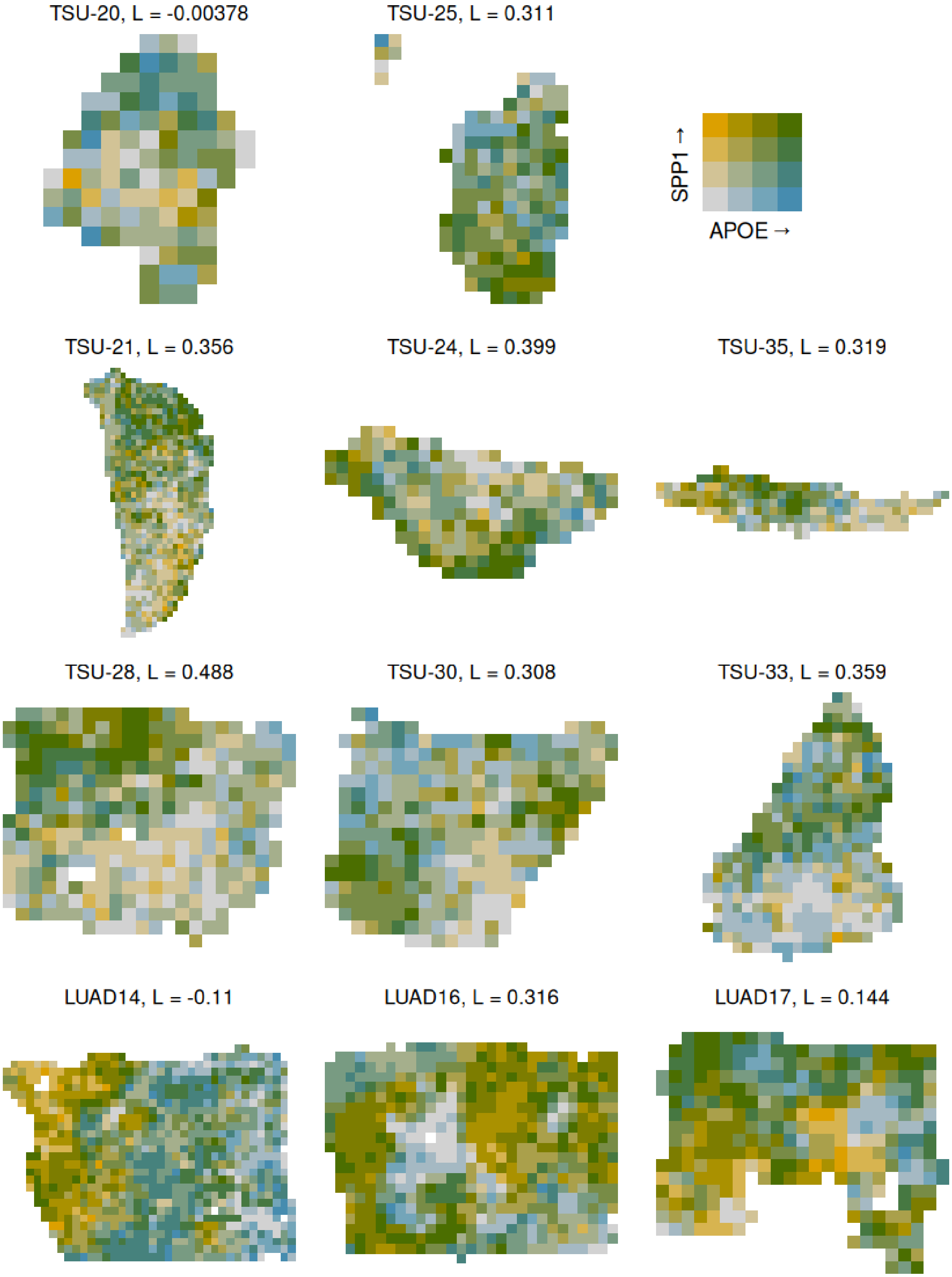
TSU-20, L = -0.00378
TSU-25, L = 0.311
SPP1 →
APOE →
TSU-21, L = 0.356
TSU-24, L = 0.399
TSU-35, L = 0.319
TSU-28, L = 0.488
TSU-30, L = 0.308
TSU-33, L = 0.359
LUAD14, L = -0.11
LUAD16, L = 0.316
LUAD17, L = 0.144

Supplementary Figure 20.2: Bivariate plot showing spatial co-expression of SPP1 and APOE in all tissues with 256 μm bins.

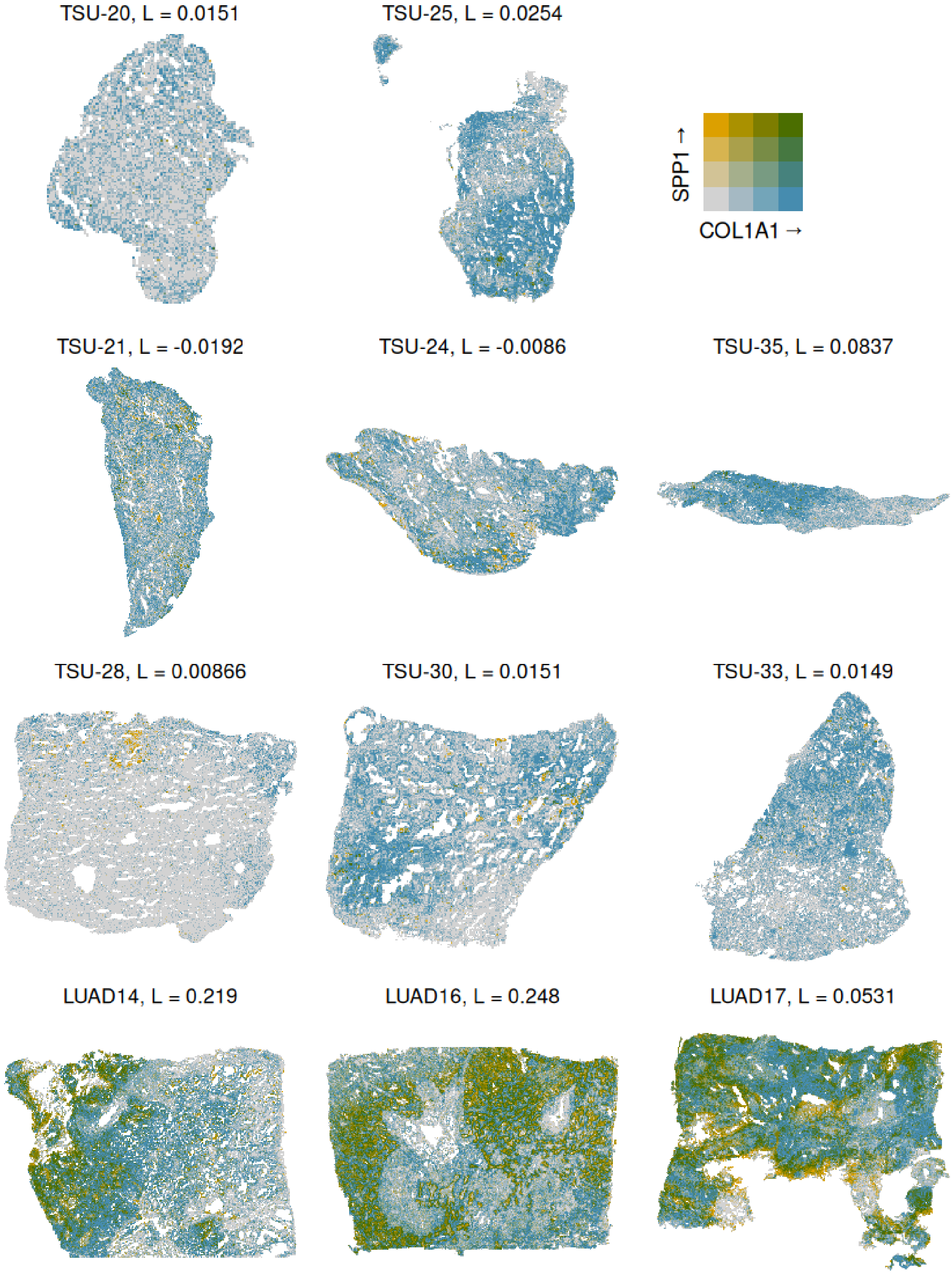
TSU-20, L = 0.0151
TSU-25, L = 0.0254
SPP1 →
COL1A1 →
TSU-21, L = -0.0192
TSU-24, L = -0.0086
TSU-35, L = 0.0837
TSU-28, L = 0.00866
TSU-30, L = 0.0151
TSU-33, L = 0.0149
LUAD14, L = 0.219
LUAD16, L = 0.248
LUAD17, L = 0.0531

Supplementary Figure 20.3: Bivariate plot showing spatial co-expression of SPP1 and COL1A1 in all tissues with 24 μm bins

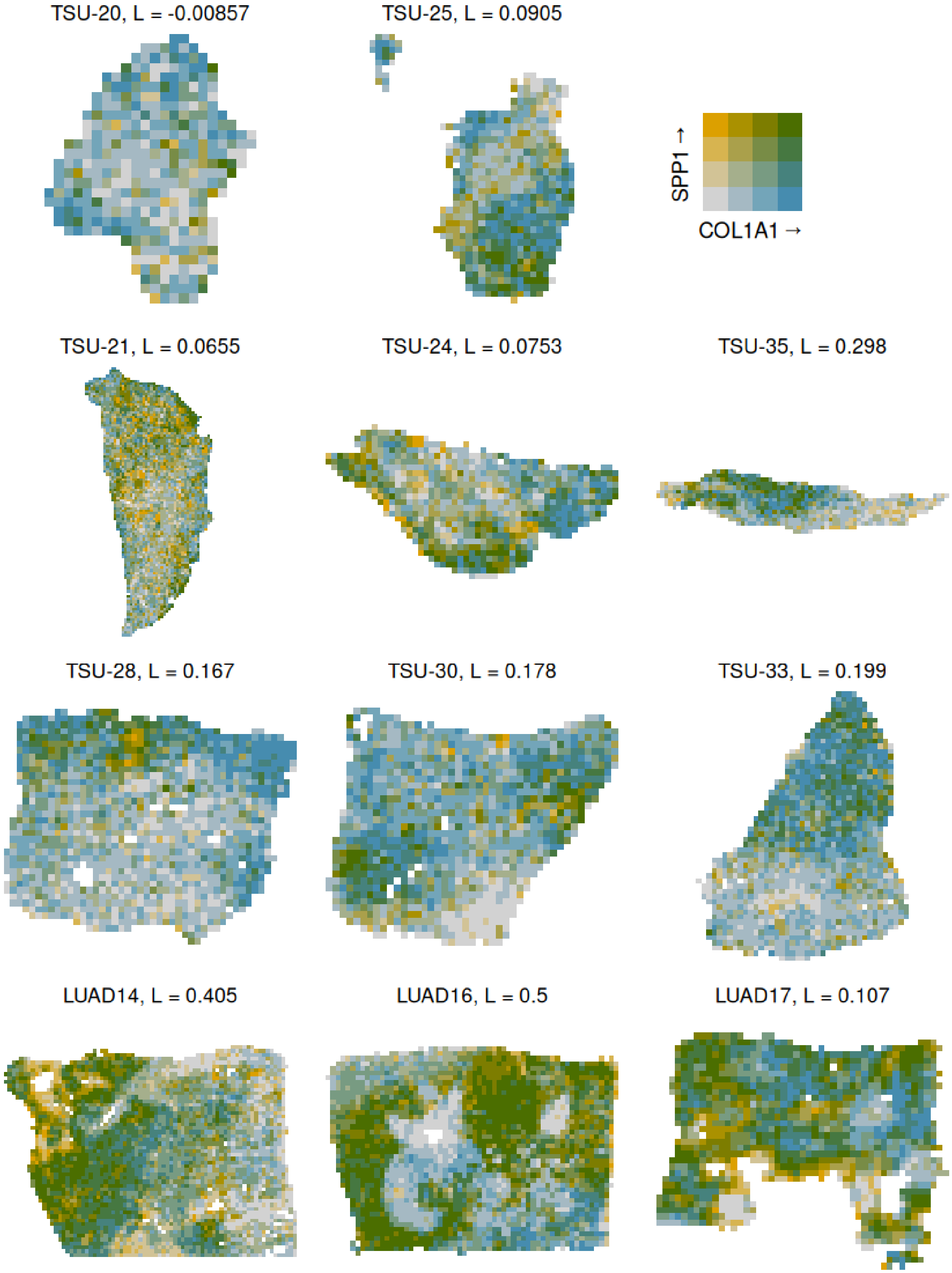

TSU-20, L = -0.00857
TSU-25, L = 0.0905
SPP1 →
COL1A1 →
TSU-21, L = 0.0655
TSU-24, L = 0.0753
TSU-35, L = 0.298
TSU-28, L = 0.167
TSU-30, L = 0.178
TSU-33, L = 0.199
LUAD14, L = 0.405
LUAD16, L = 0.5
LUAD17, L = 0.107

Supplementary Figure 20.4: Bivariate plot showing spatial co-expression of SPP1 and COL1A1 in all tissues with 128 μm bins

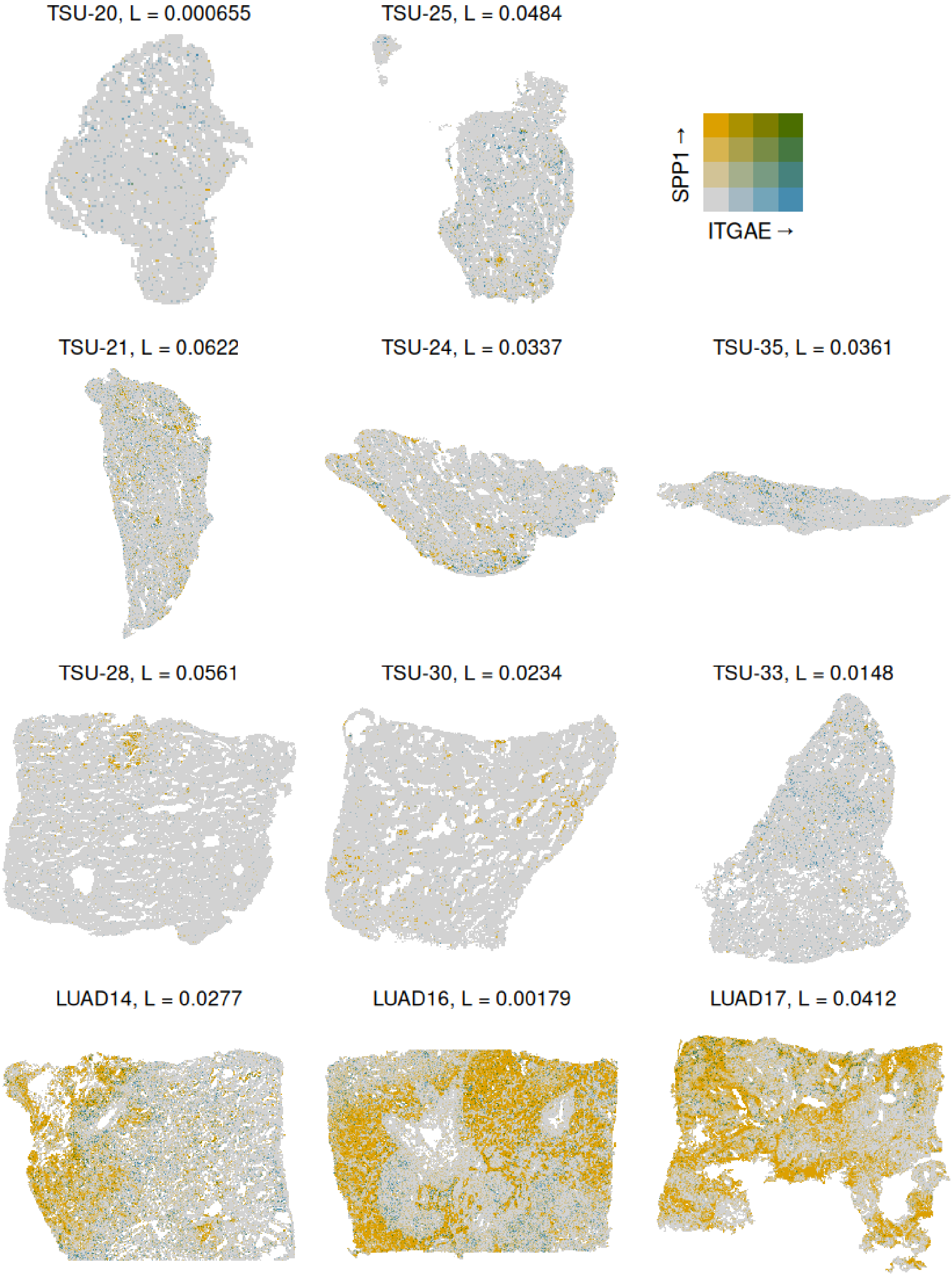

TSU-20, L = 0.000655
TSU-25, L = 0.0484
SPP1 →
ITGAE →
TSU-21, L = 0.0622
TSU-24, L = 0.0337
TSU-35, L = 0.0361
TSU-28, L = 0.0561
TSU-30, L = 0.0234
TSU-33, L = 0.0148
LUAD14, L = 0.0277
LUAD16, L = 0.00179
LUAD17, L = 0.0412

Supplementary Figure 20.5: Bivariate plot showing spatial co-expression of SPP1 and ITGAE in all tissues with 24 μm bins

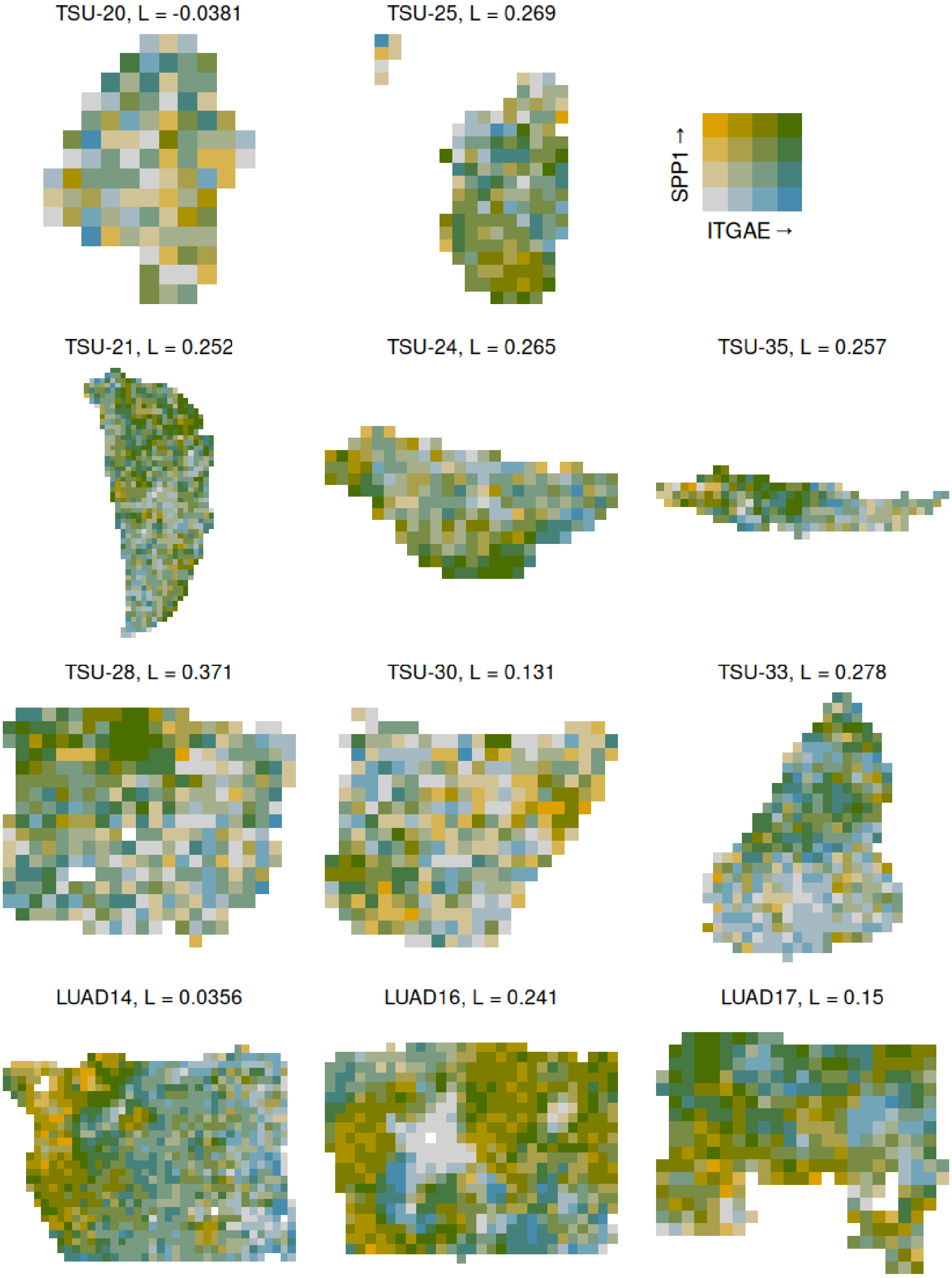

TSU-20, L = -0.0381
TSU-25, L = 0.269
SPP1 →
ITGAE →
TSU-21, L = 0.252
TSU-24, L = 0.265
TSU-35, L = 0.257
TSU-28, L = 0.371
TSU-30, L = 0.131
TSU-33, L = 0.278
LUAD14, L = 0.0356
LUAD16, L = 0.241
LUAD17, L = 0.15

Supplementary Figure 20.6: Bivariate plot showing spatial co-expression of SPP1 and ITGAE in all tissues with 256μm bins

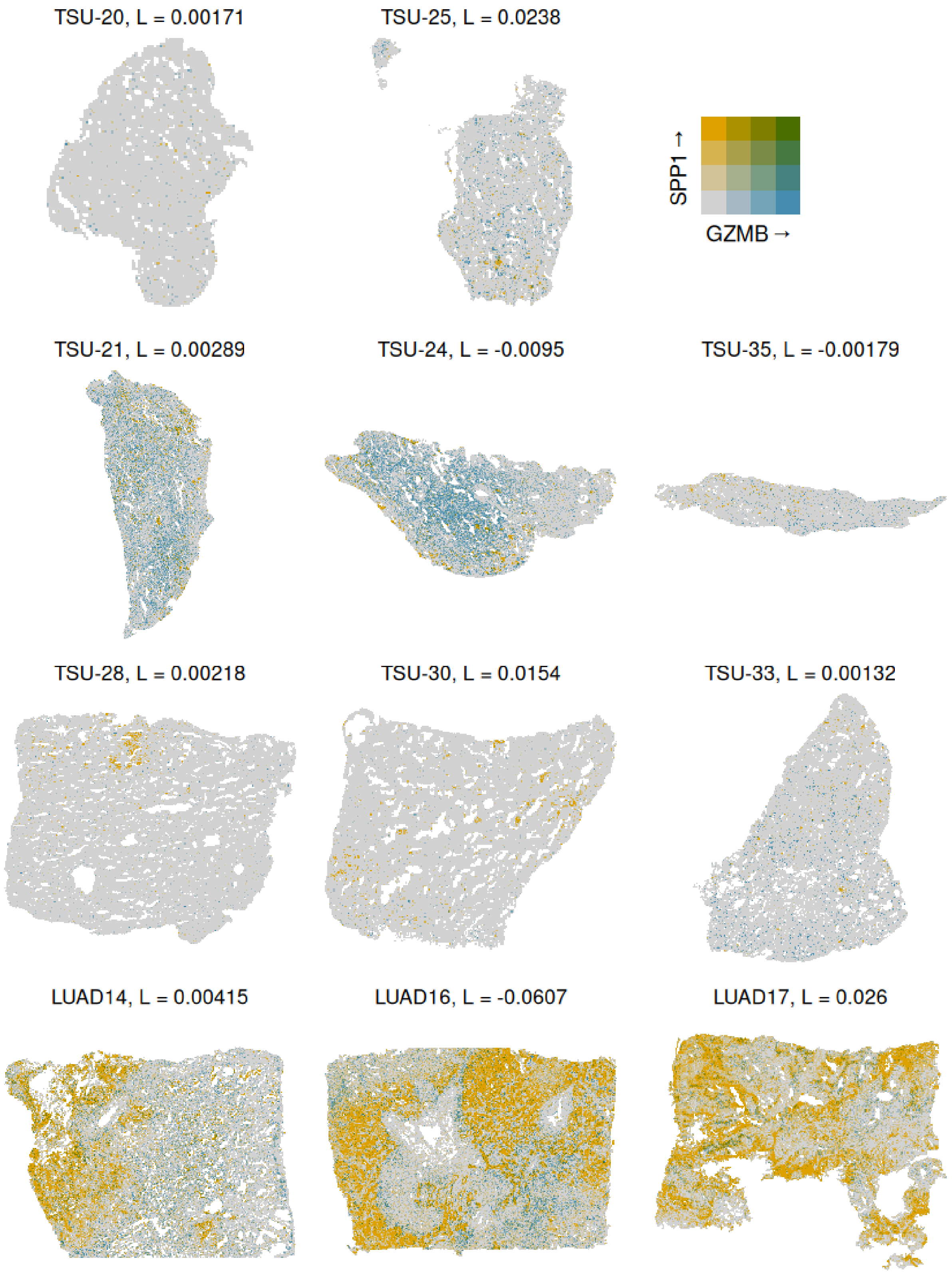
TSU-20, L = 0.00171
TSU-25, L = 0.0238
SPP1 →
GZMB →
TSU-21, L = 0.00289
TSU-24, L = -0.0095
TSU-35, L = -0.00179
TSU-28, L = 0.00218
TSU-30, L = 0.0154
TSU-33, L = 0.00132
LUAD14, L = 0.00415
LUAD16, L = -0.0607
LUAD17, L = 0.026

Supplementary Figure 20.7: Bivariate plot showing spatial co-expression of SPP1 and GZMB in all tissues with 24 μm bins

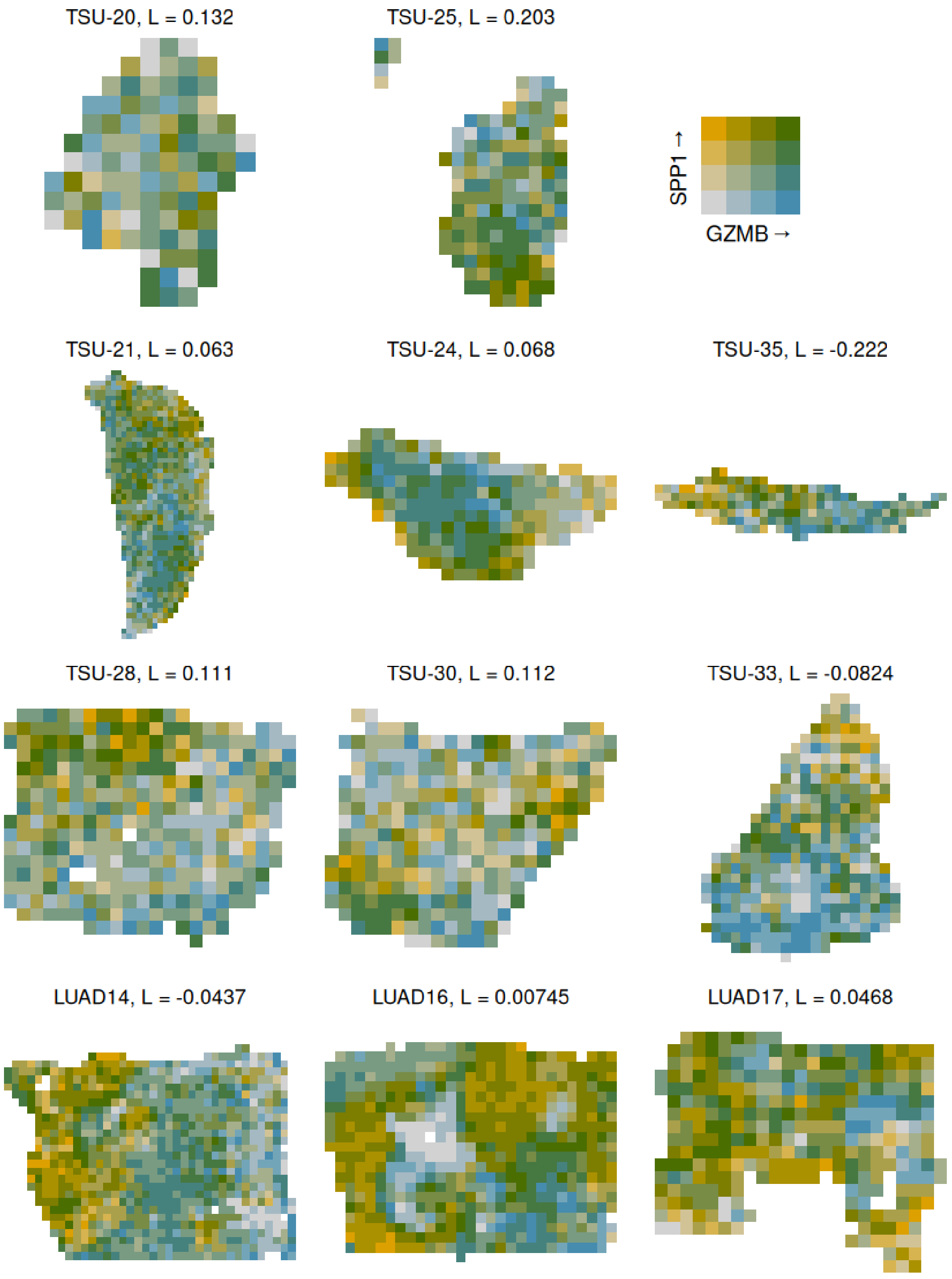

TSU-20, L = 0.132
TSU-25, L = 0.203
SPP1 →
GZMB →
TSU-21, L = 0.063
TSU-24, L = 0.068
TSU-35, L = -0.222
TSU-28, L = 0.111
TSU-30, L = 0.112
TSU-33, L = -0.0824
LUAD14, L = -0.0437
LUAD16, L = 0.00745
LUAD17, L = 0.0468

Supplementary Figure 20.8: Bivariate plot showing spatial co-expression of SPP1 and GZMB in all tissues with 256 μm bins

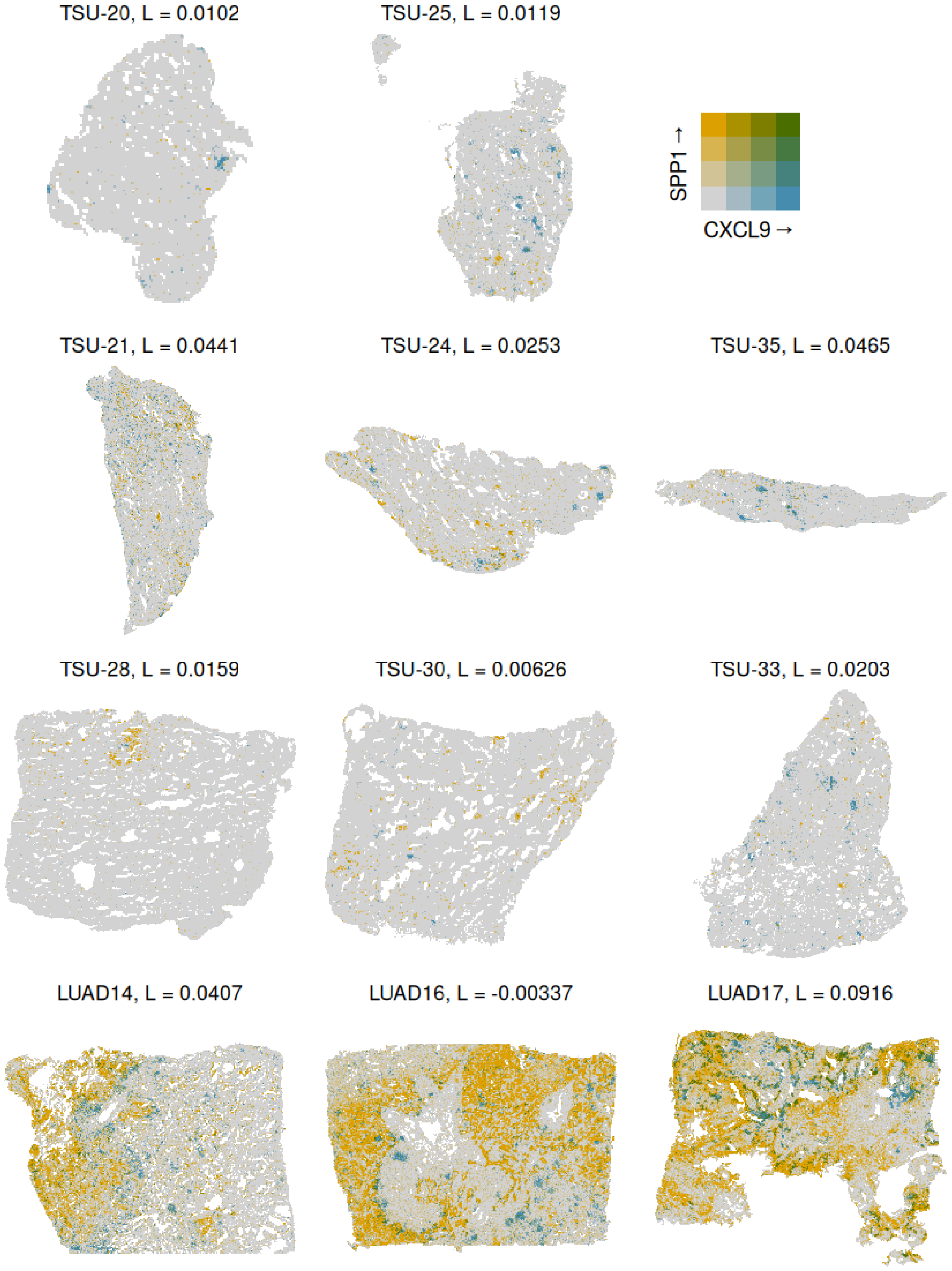
TSU-20, L = 0.0102
TSU-25, L = 0.0119
SPP1 →
CXCL9 →
TSU-21, L = 0.0441
TSU-24, L = 0.0253
TSU-35, L = 0.0465
TSU-28, L = 0.0159
TSU-30, L = 0.00626
TSU-33, L = 0.0203
LUAD14, L = 0.0407
LUAD16, L = -0.00337
LUAD17, L = 0.0916

Supplementary Figure 20.9: Bivariate plot showing spatial co-expression of SPP1 and CXCL9 in all tissues with 24 μm bins

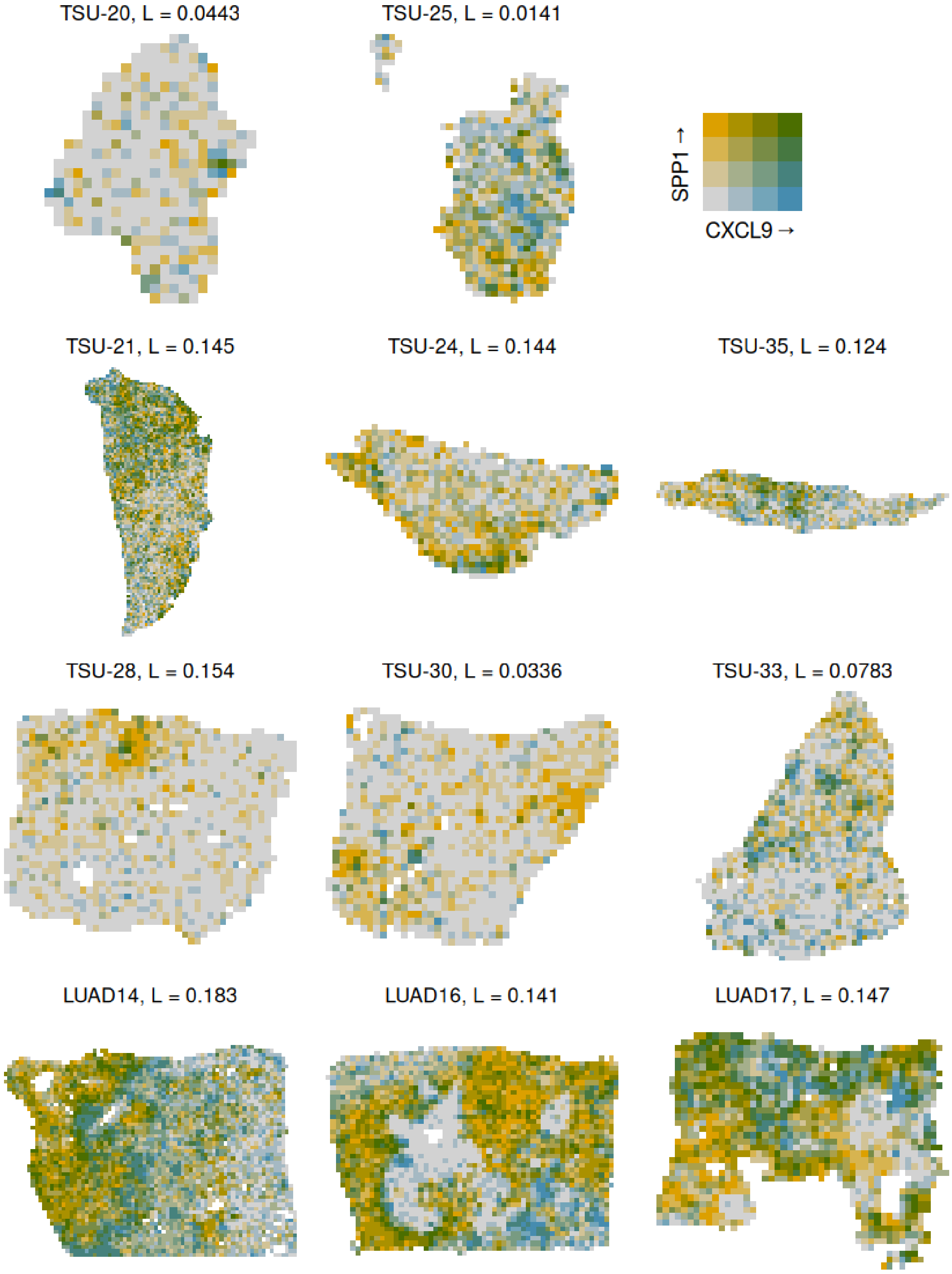
TSU-20, L = 0.0443
TSU-25, L = 0.0141
SPP1 →
CXCL9 →
TSU-21, L = 0.145
TSU-24, L = 0.144
TSU-35, L = 0.124
TSU-28, L = 0.154
TSU-30, L = 0.0336
TSU-33, L = 0.0783
LUAD14, L = 0.183
LUAD16, L = 0.141
LUAD17, L = 0.147

Supplementary Figure 20.10: Bivariate plot showing spatial co-expression of SPP1 and CXCL9 in all tissues with 128 µm bins

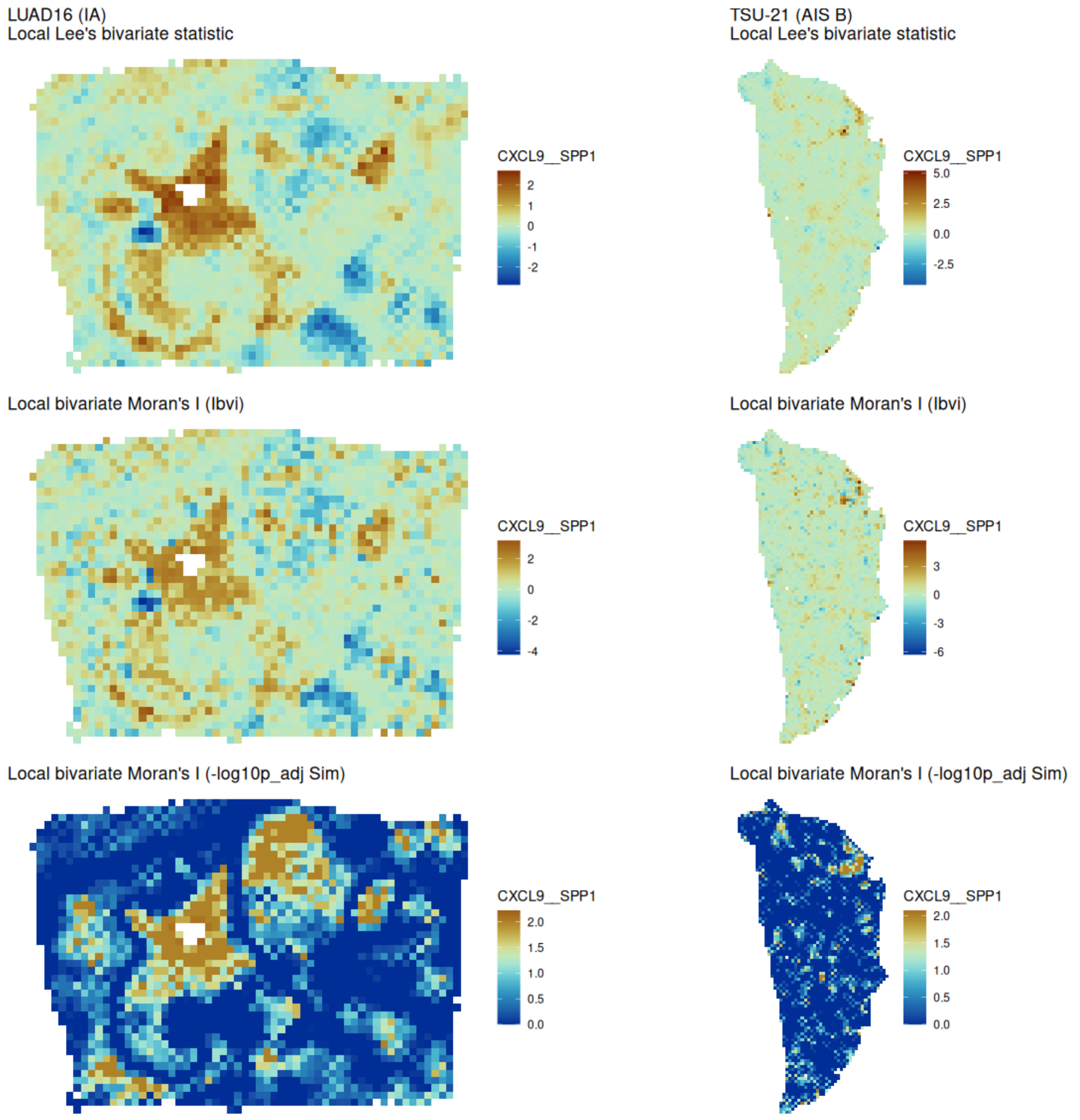


Supplementary Figure 21.1: Local Lee’s L, local bivariate Moran’s I, and p-values of local bivariate Moran’s I of CXCL9 and SPP1 from permutation testing plotted on 128 µm bins, in LUAD16 (IA) and TSU-21 (AIS B).

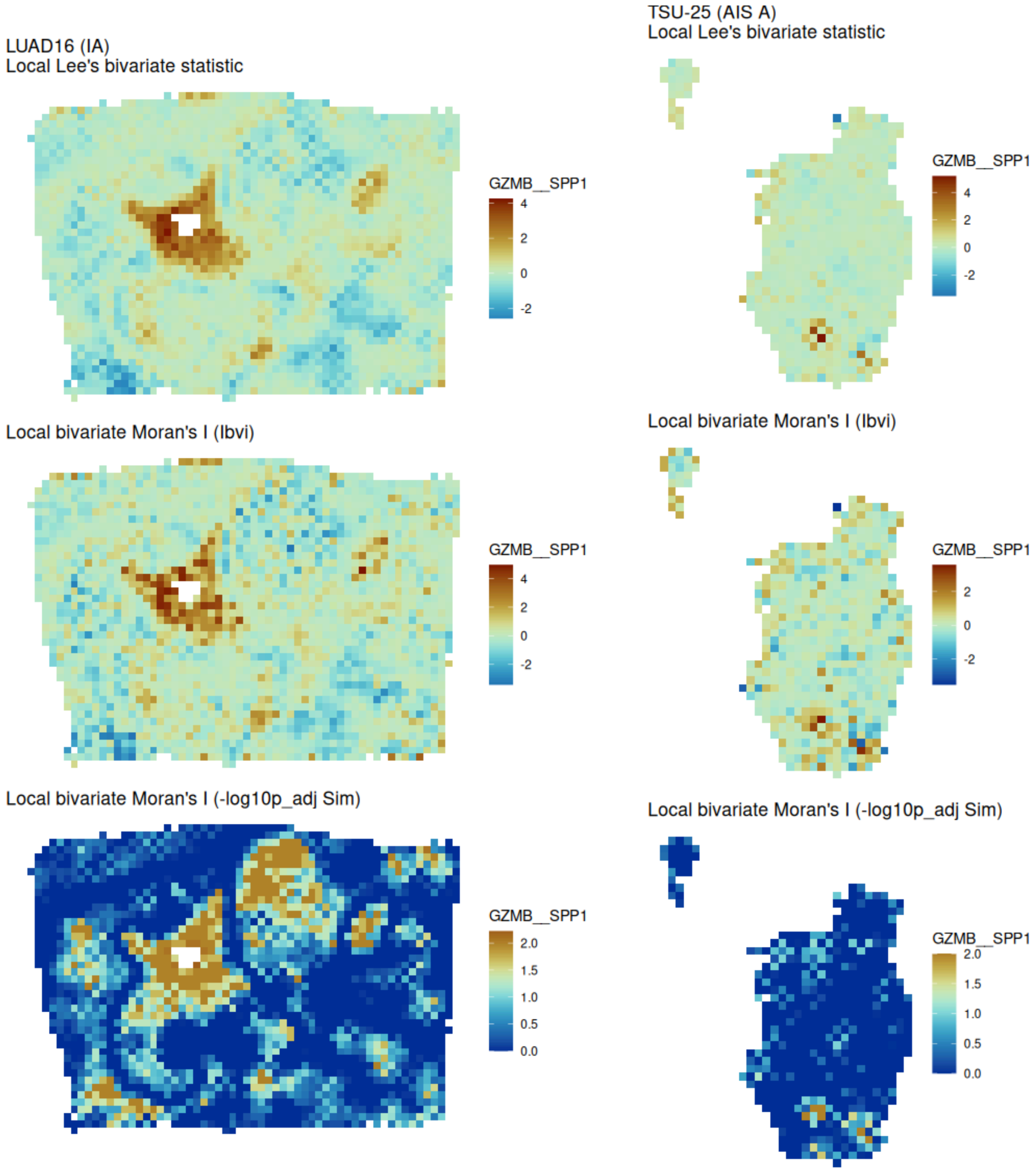


Supplementary Figure 21.2: Local Lee's L, local bivariate Moran's I, and p-values of local bivariate Moran's I of GZMB and SPP1 from permutation testing plotted on 128 µm bins, in LUAD16 (IA) and TSU-25 (AIS A).

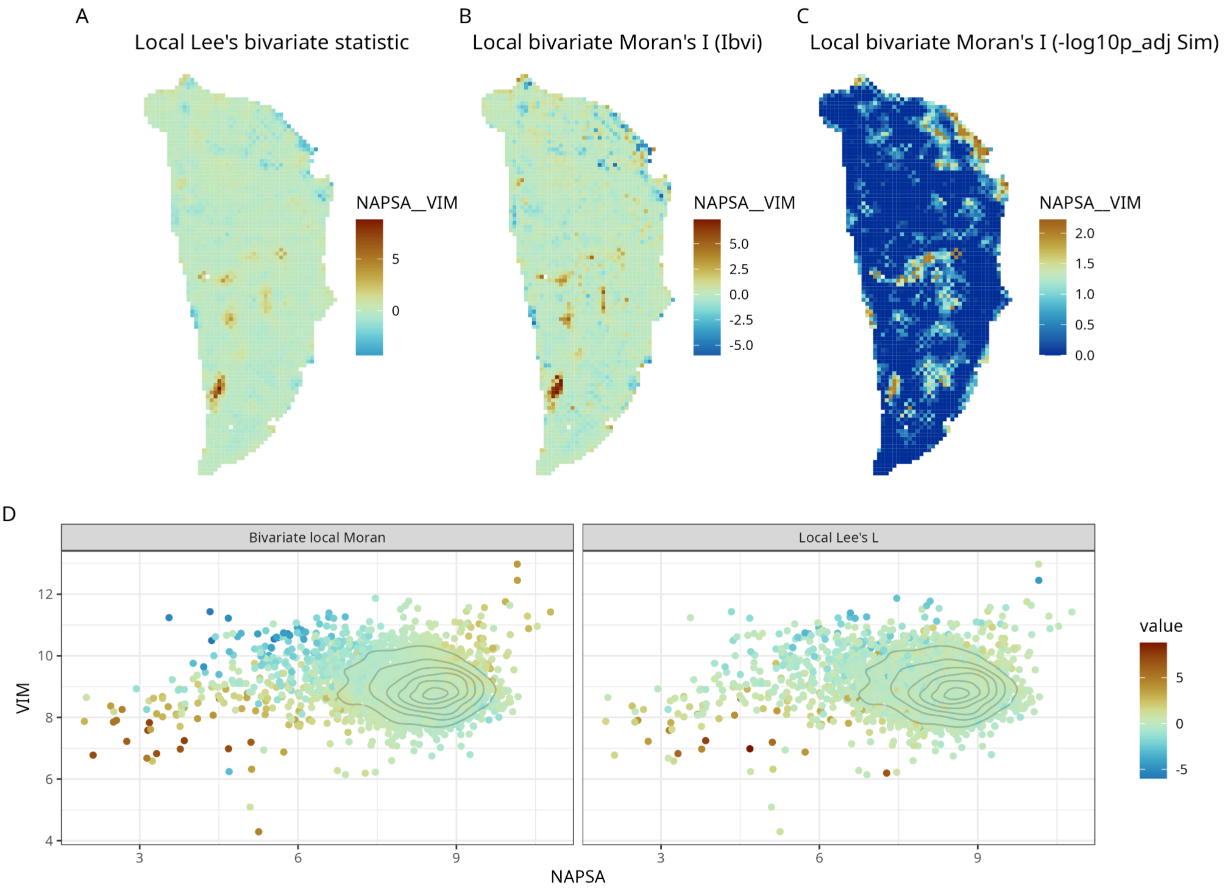


Supplementary Figure 22: Local contributions to bivariate statistic in Supplementary Figure 9.3D. While Lee's L is close to 0 with bin size 256 for NAPA-VIM, there is clearly a spatial structure, so we computed local bivariate spatial statistics to see local contributions to global Lee's L. A) Local Lee's L statistic plotted in space. B) Local bivariate Moran's I plotted in space. C) -log10 adjusted p-value from permutations for local bivariate Moran's I plotted in space (see Methods); warm colors indicate adjusted $p < 0.05$. D) Scatter plot of log normalized expression values of VIM vs. NAPA, colored by local bivariate Moran's I and local Lee's L. Gray contours indicate point density. Bins with high positive values of local Lee's L or local bivariate Moran's I mostly have lower than average expression in both, while bins with more negative values mostly have higher than average values in VIM and lower than average values in NAPSA. This indicates that the near 0 value of global Lee's L is caused by local regions with positive and negative correlations canceling out instead of a lack of spatial structure in co-expression. This points to the importance of local spatial statistics and limitations of this global metric.

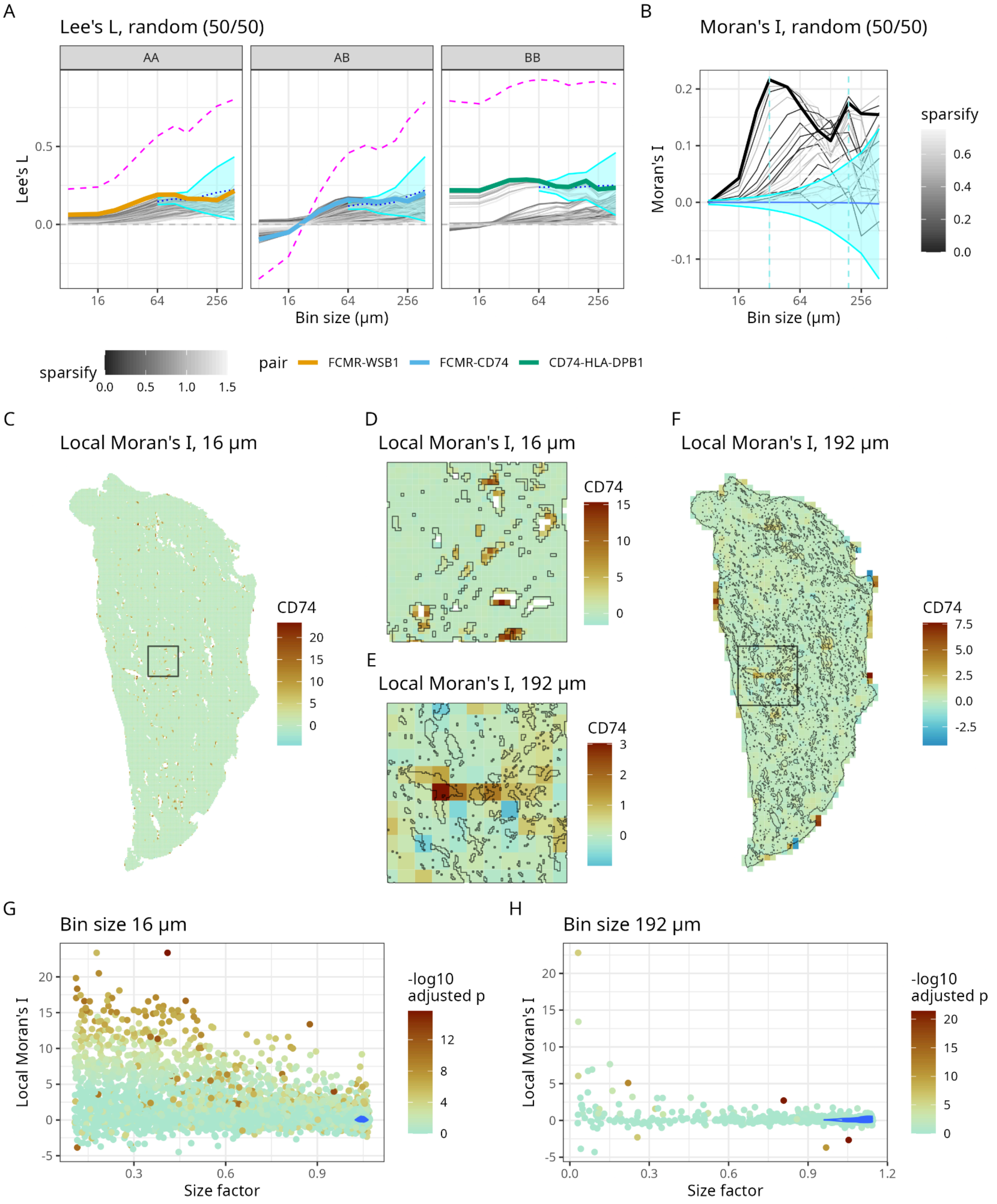


Supplementary Figure 23: Spurious cases of Lee's L and Moran's I. A) Lee's L vs. bin size in a synthetic pattern that does not have a spatial pattern; "cell types" A and B were assigned randomly so the number of A and B 8 μm bins are the same. The magenta dashed lines are Pearson correlations of the same feature pairs highlighted in thick lines, which have the strongest correlation in their respective categories and are the most highly "expressed". For the feature pairs highlighted in thick lines, the cyan ribbon is 2.5 and 97.5 percentiles under null hypothesis, multiplied by the number of bin sizes and number of synthetic features for Bonferroni correction. This interval depends on the non-spatial correlation between the two features and hence differs for each feature pair. Since it's only computationally feasible to compute the mean and variance of Lee's L for smaller numbers of bins, the intervals were only computed for larger bin sizes. The observed values of Lee's L are close to the expected values (thin dotted lines) and well within that interval for bin sizes larger than 64 μm, so for those larger bin sizes, Lee's L is not statistically significant despite values away from 0 due to the high Pearson correlation. B) Moran's I vs. bin size for the non-spatial synthetic pattern. Local Moran's I of the feature whose curve is highlighted in thick line is plotted in panels C-H. C) Local Moran's I values plotted in space on the entire section, with 16 μm bins. D) Zooming into the box marked in C. The thin gray lines mark the boundaries of holes; many bins with high local Moran's I values are at the edges of holes. E) Zooming into the box marked in F. Thin gray lines mark holes for 16 μm bins. Bins with high local Moran's I tend to be around regions with more holes. F) Local Moran's I values plotted in space on the entire section, with 192 μm bins. Thin gray lines mark holes for 16 μm bins. Bins with high local Moran's I tend to be at the edge or in regions with more holes. G) Local Moran's I vs. size factor (proportional to proportion of the bin covered by segmented cells) for 16 μm bins. Blue lines are contours for point density. Each point is a bin and the points are colored by -log10 adjusted p-values of local Moran's I, so warm color indicates adjusted $p < 0.05$. Bins with higher local Moran's I mostly have lower size factor, i.e. they are not fully occupied by cells. H) Same as G but for 192 μm bins.
